# Efficient generation of epitope-targeted *de novo* antibodies with Germinal

**DOI:** 10.1101/2025.09.19.677421

**Authors:** Luis S. Mille-Fragoso, Claudia L. Driscoll, John N. Wang, Haoyu Dai, Talal Widatalla, Jim L. Zhang, Xiaowei Zhang, Bing Rao, Liang Feng, Brian L. Hie, Xiaojing J. Gao

**Affiliations:** Department of Bioengineering, Stanford University, Stanford, CA, USA; Sarafan ChEM-H, Stanford University, Stanford, CA, USA; Stanford Bio-X, Stanford University, Stanford, CA, USA; Arc Institute, Palo Alto, CA, USA; Department of Chemical Engineering, Stanford University, Stanford, CA, USA; Department of Computer Science, Stanford University, Stanford, CA, USA; Stanford Biophysics, Stanford University, Stanford, CA, USA; Department of Structural Biology, Stanford University, Stanford, CA, USA; Department of Molecular and Cellular Physiology, Stanford University, Stanford, CA, USA; Stanford Data Science, Stanford University, Stanford, CA, USA

## Abstract

Obtaining novel antibodies against specific protein targets is a widely important yet experimentally laborious process. Meanwhile, computational methods for antibody design have been limited by low success rates that currently require resource-intensive screening. Here, we introduce Germinal, a broadly enabling generative pipeline that designs antibodies against specific epitopes with nanomolar binding affinities while requiring only low-n experimental testing. Our method co-optimizes antibody structure and sequence by integrating a structure predictor with an antibody-specific protein language model to perform *de novo* design of functional complementarity-determining regions (CDRs) onto a user-specified structural framework. When tested against four diverse protein targets, Germinal successfully designed functional antibodies across all targets and binder formats, testing only 43-101 designs for each antigen. Validated designs also exhibited robust expression in mammalian cells and high sequence and structural novelty. We provide open-source code and full computational and experimental protocols to facilitate wide adoption. Germinal represents a milestone in efficient, epitope-targeted *de novo* antibody design, with notable implications for the development of molecular tools and therapeutics.

## 1. Introduction

Antibodies play a central role in adaptive immunity by binding a remarkable diversity of molecular epitopes on target antigens, often with high specificity (Hozumi and Tonegawa, 1976; Victora and Nussenzweig, 2022). Together with well-characterized biochemical properties and favorable therapeutic profiles, these capabilities have made antibodies widely adopted as general-purpose and high specificity binders in biomedicine, biotechnology, and basic research (Bailly et al., 2020; Köhler and Milstein, 1975).

Antibody generation against arbitrary antigens traditionally requires animal immunization or large library screening campaigns (Fridy et al., 2024; Alexander and Leong, 2024). However, these experimental approaches are constrained by fundamental limitations. First, these processes are laborious and expensive, and do not always yield successful binders (Wilson and Andrews, 2012). Second, once binders are identified, their molecular and structural characterization can be difficult (Cheng et al., 2024; Fekete et al., 2013). Finally, these methods offer no control over which specific region of the antigen the antibody recognizes, known as the epitope, making it difficult to direct binding to functionally important sites or specific target conformations.

Advances in machine learning have led to atomically accurate structure prediction of protein monomers and multimeric complexes with methods such as AlphaFold (Jumper et al., 2021; Evans et al., 2022). Subsequent efforts have also leveraged these structure predictors to design protein sequences that have desirable structural properties, including binding interactions. For example, methods such as RFdiffusion have fine-tuned structure prediction models with diffusion objectives to generate the backbone coordinates of potential protein binders (Watson et al., 2023; Baek et al., 2021). Alternatively, several studies have leveraged gradient-based methods that invert structure prediction to generate proteins and protein binders (Frank et al., 2024; Jendrusch et al., 2025; Goverde et al., 2023; Wicky et al., 2022). Recently, BindCraft inverted AF-M to obtain high experimental success rates for *de novo* miniproteins that bind arbitrary targets (Pacesa et al., 2025; Evans et al., 2022).

However, despite numerous preliminary efforts (Bennett et al., 2025; Shanehsazzadeh et al., 2023; Nabla Bio and Biswas, 2025), robust *de novo* antibody design against specific target epitopes with high success rates remains challenging, especially due to the inherent hyper-variability of their complementarity determining regions (CDRs) and the highly constrained nature of antibody sequence space. For example, efforts to design novel CDRs have leveraged large-scale library screens containing thousands of designs to ultimately yield a handful of binders, often with activity in the micromolar range (Bennett et al., 2025). These experimental success rates require resource-intensive screening campaigns that are inaccessible to most laboratories. In contrast, advances in generative modeling that reduce experimental validation to tens rather than thousands of designs would democratize access to these technologies across molecular biology.

Here, we present Germinal, a broadly enabling generative pipeline for *de novo* antibody design (**Figure 1A**). In particular, Germinal achieves epitope-targeted, *de novo* CDR design with experimental success rates that enable low-n experimental testing. Germinal leverages a computational pipeline that biases protein binders to a specified antibody framework while freely designing the CDRs, a strategy that not only maintains robust expression in human cells, but also preserves the therapeutic developability profiles conferred by known framework regions (FR). To achieve this, we combine backpropagation of AlphaFold-Multimer (AF-M) (Evans et al., 2022) with an antibody-specific protein language model (IgLM; Shuai et al. (2023)), allowing us to produce designs with realistic CDR sequences (**Figure 1B**). Similarly, Germinal uses custom loss functions that ensure correct spatial positioning of the designs to favor CDR over framework binding contacts. We identified lead antibody designs for using a split-luciferase assay or a preliminary surface plasmon resonance (SPR; A.2.11) screen followed by validation by bio-layer interferometry (BLI), enabling robust and efficient identification of high-affinity binders.

**Figure 1.**
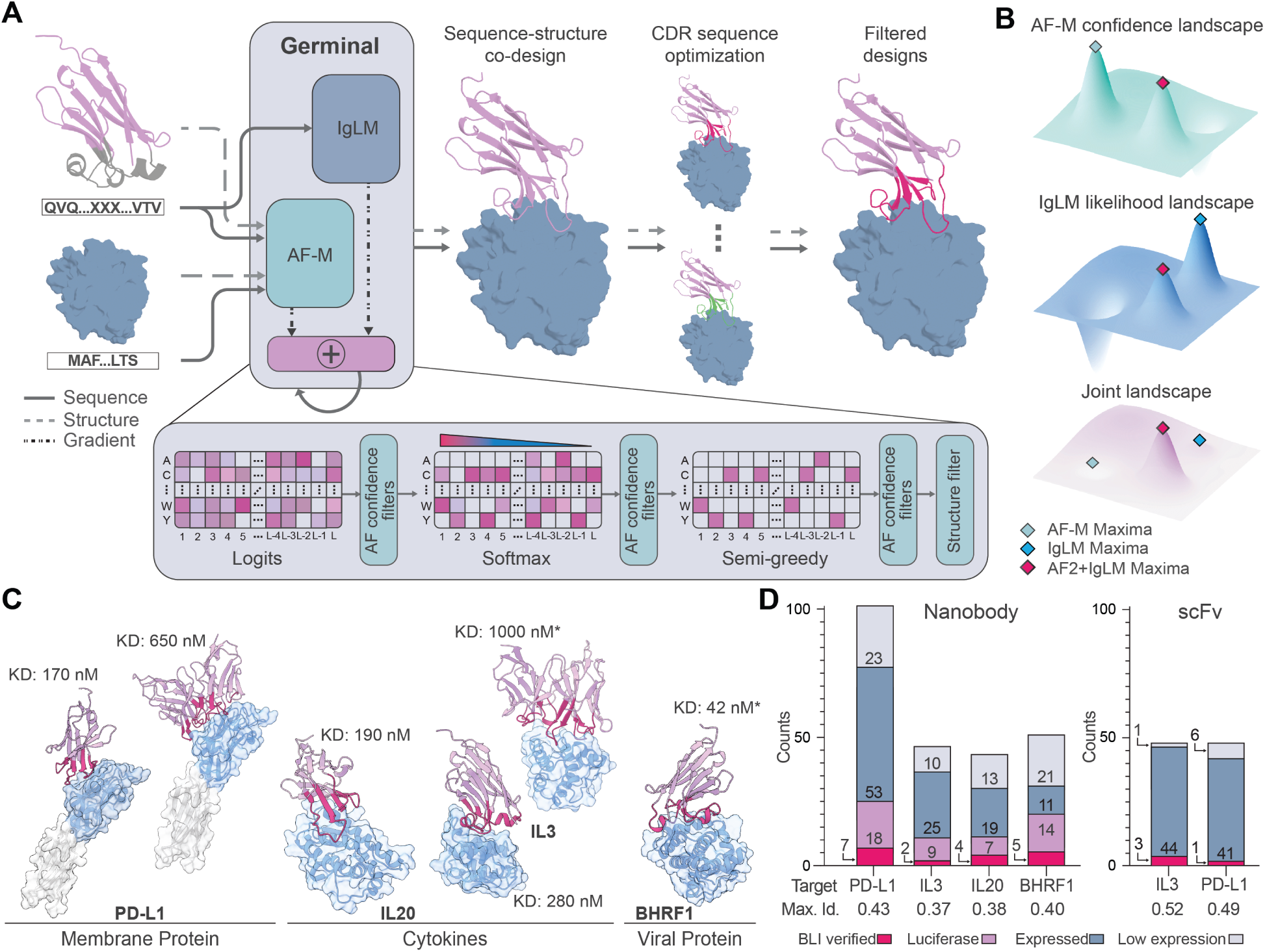
Efficient antibody generation via joint optimization of AlphaFold-Multimer and IgLM with Germinal. (**A**) Overview of the Germinal pipeline. A target and antibody framework are provided as structural templates and sequence inputs while CDRs are freely designed. Gradients from AlphaFold-Multimer (AF-M) and an antibody-specific language model (IgLM) are iteratively combined throughout the design stage to optimize sequences during three phases (Logits → Softmax (temperature annealing) → Semi-greedy). (**B**) Structure and sequence joint optimization landscape. AF-M (structure/complex-compatibility) and IgLM (antibody sequence prior) objectives define distinct optima. Their joint objective steers optimization toward sequences that are simultaneously structurally confident and antibody-like. (**C**) Representative predicted complexes across the four targets. Designed nanobodies or single-chain variable fragments (scFvs) bound to specified epitopes on PD-L1, IL3, IL20, and BHRF1 are shown in pink with CDRs highlighted in magenta. Domains used for design are colored in blue. White domains were excluded from the target structure used during design. Best dissociation constants (𝐾_𝐷_) measured by BLI for each target are annotated. Structures shown are AF3-predicted. Asterisks (*) denote binders derived from an initial Germinal design (see **Section 2.3**). (**D**) Screening results by target for designed nanobodies (*left)* and scFvs (*right)*. Stacked bars summarize the per-library results (43 to 101 designs per target). Designs fall into four categories depending on their expression and binding profiles. Low-expression (gray), expressed but filtered as non-binder in split-luciferase assay (blue), binders exhibiting significant signals in the split-luciferase assay (pink), and BLI-verified binders (magenta). Numbers indicate counts within each category. scFv designs were screened directly in fragment antibody (Fab) format by SPR/BLI, and thus the luciferase-related categories are omitted from the bar plot. “Max. Id.” denotes the highest pairwise CDR sequence identity of any design in the library to the PDB. For scFvs, the Max. Id. was calculated independently for the variable heavy (VH) and variable light (VL) chains and then averaged. AF-M: AlphaFold-Multimer; IgLM: antibody-specific language model; CDR: complementarity-determining region; BLI: bio-layer interferometry; SPR: surface plasmon resonance; 𝐾_𝐷_: dissociation constant.

Targeting epitopes across four distinct antigens (**Figure 1C**), we obtained experimentally validated binders following screens of only tens of *de novo* nanobody or single-chain variable fragments (scFv) designs per target (**Figure 1D**). Subsequent affinity characterization by BLI revealed that validated designs of both formats achieved nanomolar-to-low-micromolar dissociation constants (𝐾_D_) for all antigens tested. Cryo-EM structural determination and alanine mutagenesis at computationally specified hotspot residues confirmed that most designs engage their intended epitopes with atomic-level accuracy. Furthermore, all designs exhibited low polyreactivity comparable to nanobody and antibody controls with previously determined specificity.

To promote broad accessibility, we make our full computational pipeline open-source and freely available to the scientific community, alongside our efficient experimental protocol for initial binder screening. We anticipate that Germinal will greatly reduce the need for the extensive experimental infrastructure developed specifically for antibody discovery. By supporting precise epitope targeting while maintaining favorable antibody properties, Germinal opens new possibilities for the development of antibody molecules for molecular biology and therapeutic design.

## 2. Results

### 2.1. Design of antibodies via dual-objective structure and sequence optimization

Computational design of *de novo* antibodies remains challenging due to their specific structural constraints, the high variability of their binding regions (CDRs), and a lack of antibody training data for structure prediction tools (Ruffolo et al., 2023; Hitawala and Gray, 2024). In natural antibodies, the conserved 𝛽-sheet framework regions serve as a rigid scaffold that positions flexible CDR loops for optimal antigen recognition (North et al., 2011). Although secondary structures do occur in CDRs, flexible loops (where flexible refers to the paucity of secondary structures rather than conformational change upon target binding (Fernández-Quintero et al., 2019; Spoendlin et al., 2025; Liu et al., 2024)) are more prevalent in natural antibodies (Spoendlin et al., 2025; Mitchell and Colwell, 2018). Binder design methods built on these structure predictors, however, are biased toward protein–protein interfaces dominated by secondary structures rather than the more flexible loop conformations characteristic of CDRs (Watson et al., 2023; Pacesa et al., 2025; Lu et al., 2025).

Antibody-specific language models offer a complementary source of information. Trained on vast repositories of antibody sequences (Olsen et al., 2022a), these models learn the distribution of natural antibody sequences and capture implicit structural information from sequence data alone (Shuai et al., 2023; Olsen et al., 2022b; Rao et al., 2021; Verkuil et al., 2022). Consistent with this, recent work has shown that general structure-based models alone struggle to redesign functional CDRs, but perform substantially better when complemented by antibody-specific language models (Alamo et al., 2025). We therefore hypothesized that integrating structure prediction with antibody-specific language models could overcome the twin challenges of data scarcity and structural bias, unlocking efficient *de novo* antibody design. Germinal realizes this integration by combining a protein structure predictor, AF-M (Evans et al., 2022), with an antibody-specific protein language model, IgLM (Shuai et al., 2023) (**Figure 1A**), merging their gradients to co-optimize both fold geometry and antibody naturalness in a joint optimization landscape (**Figure 1B**). Germinal’s generation pipeline consists of three main stages: design, sequence optimization, and filtering.

In the design stage, Germinal leverages the gradients of AF-M to sample sequences that, with high confidence, bind desired epitopes in the predicted structures. Germinal also leverages the gradients of IgLM to bias sampling toward sequences with high probability under the language model, reflecting their likelihood of resembling naturally occurring antibodies. Given AF-M’s direct conditioning on input sequences and the intrinsic coupling of sequence and structure, the antibody language model also implicitly influences the resulting structure. During this stage, Germinal also allows users to explicitly bias samples toward a desired antibody framework sequence (e.g., a framework with a favorable developability profile; **Figure S1**), as well as augmenting structure guidance by providing the framework’s structure as a template to AF-M (**Methods**).

Sampled sequences with high-quality predicted structures are then carried forward to a sequence optimization stage. Similar to other hallucination methods, we use AbMPNN (Dreyer et al., 2023; Dauparas et al., 2022)), a structure-conditioned sequence design model fine-tuned on antibodies, to redesign CDR residues that are not in direct contact with the antigen. This serves two purposes: improving binder stability by optimizing non-interface CDR residues while preserving key paratope–epitope contacts, and generating multiple sequence variants per design structure, effectively expanding the diversity of the candidate pool without requiring additional computational trajectories (**Methods; Table S1**).

Finally, designs are passed through a filtering stage (**Tables S2; S3**), where we use a different structure predictor from the one used during the design stage to provide a separate assessment of design quality; in particular, we used AlphaFold 3 (AF3) due to its superior accuracy on antibody–antigen complexes (Abramson et al., 2024; Hitawala and Gray, 2024). We filter and rank (**Tables S4; S5**) designs based on AF3 confidence scores alongside PyRosetta-derived biophysical and biochemical scores (Chaudhury et al., 2010), yielding a final set of candidates for experimental testing (**Methods**).

We initially found that naive application of AF-M guidance towards antibody binder design resulted in generations with paratopes (the region of the antibody that contacts the target) composed of framework residues or enriched with secondary-structures. To address this, Germinal incorporates three custom loss functions during the design stage (**Figure 2A**) that effectively guide generations toward natural antibody binding conformations in contrast to those favored by naive AF-M. A paratope-specific loss ensures that binding occurs primarily through the designed CDRs (**Figure 2B**). Additionally, Germinal incorporates two secondary-structure losses, an 𝛼-helix loss and 𝛽-strand loss. These losses encourage the generated CDRs to adopt loop conformations, instead of the secondary-structure–rich conformations from vanilla AF-M guidance (visualized in **Figure 2C,D**). Together, these losses guide designs toward binding interfaces dominated by CDRs (Fernández-Quintero et al., 2018; Kunik et al., 2012; Tsuchiya and Mizuguchi, 2016) while promoting their conformational flexibility, a feature that has been linked to enhanced antigen specificity and the capacity to generate broadly neutralizing binders in natural antibodies (Li et al., 2015; Prigent et al., 2018).

**Figure 2.**
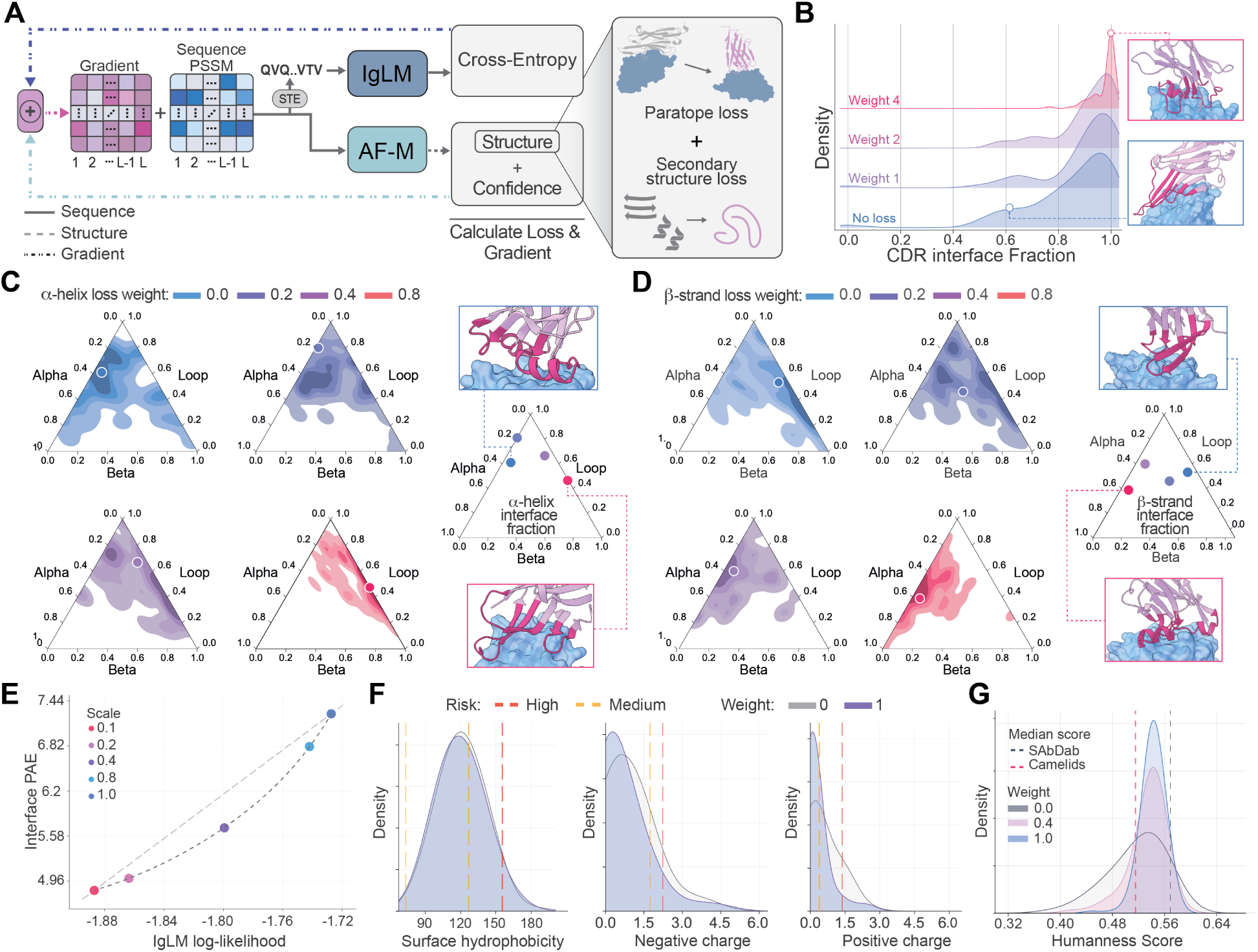
Germinal’s design stage steers generations toward natural antibody-like properties through structural and sequence guidance. (**A**) Schematic of Germinal’s design stage. Structure and confidence losses are integrated with IgLM sequence guidance through gradient merging, and the resulting gradient is used to update the sequence PSSM. A straight-through estimator converts the PSSM into a one-hot encoded sequence for input to IgLM, whereas AF-M operates directly on the PSSM. (**B**) Paratope loss ensures binding occurs primarily through CDR regions rather than framework residues. (**C-D**) Secondary structure losses prevent CDRs from being dominated by 𝛼-helices and 𝛽-strands. Ternary kernel density plots display the secondary structure composition of the interface for the designs. Selected points represent the median interface fraction for designs generated at each weight. (**E**) Pareto frontier between AF-M structural confidence, shown as iPAE, and IgLM sequence likelihood reveals competing optimization objectives. (**F**) Comparison of developability properties from the Therapeutic Nanobody Profiler (TNP) between sequences generated with (weight of 1.0) and without (weight of 0) IgLM guidance. Designs within the amber region are defined as low risk, between amber and red, medium risk, and above red, high risk. (**G**) Comparison of the OASis humanness score amongst sequences generated with varying weighting of IgLM guidance. Shown in gray and pink are the median humanness scores of all nanobodies, and all camelid nanobodies, respectively, in the SAbDab-nano database.

While combining AF-M structure guidance with IgLM sequence guidance is central to our approach, we found that the preference an antibody language model has for a given sequence (IgLM log-likelihood) and the predicted binding confidence (AF-M interface predicted aligned error; iPAE) are competing objectives. This behavior is expected given that antibody-antigen complexes have been challenging to model (Spoendlin et al., 2025; Hitawala and Gray, 2024), as well as the fact that backpropagating through AF-M can lead to adversarial designs with unrealistic poses that improve structure confidence. This trade-off yields a Pareto frontier, underscoring the need for joint optimization (**Figure 2E** and **S2**). As the strength of IgLM guidance increased, we observed improvements in the therapeutic developability and safety profiles for generated sequences as predicted by the Therapeutic Nanobody Profiler (TNP) (Gordon et al., 2025) (**Figure 2F; Figure S3**) and Therapeutic Antibody Profiler (TAP) (Raybould et al., 2019) (**Figure S4**). Additionally, IgLM guidance improved the "humanness" and reduced the immunogenicity risk of generated sequences, as predicted by the OASis tool (Prihoda et al., 2022), which calculates the similarity of a given sequence against a large database of human antibodies (Olsen et al., 2022a) (**Figure 2G**). In total, Germinal enables controllable, multi-objective generation of antibody-like sequences.

### 2.2. Using Germinal to target diverse antigens

We next sought to apply Germinal to four diverse targets: Protein Death Ligand 1 (PD-L1), an immune checkpoint ligand expressed on both tumor and immune cells, which is both a clinically validated therapeutic target and a frequent target of *de novo* binder design efforts; interleukin-3 (IL3) and interleukin-20 (IL20), cytokines involved in immune signaling for which no *de novo* binders have been reported to date; and BHRF1, a viral Bcl-2–like anti-apoptotic protein from the Epstein–Barr virus, as a representative example of a non-human, pathogen-derived target.

After sampling, we observed that 1,584 nanobody trajectories for PD-L1, 1,379 trajectories for IL3, 739 trajectories for IL20, and 1,456 trajectories for BHRF1 had passed the pipeline filters with promising computational metrics (**Figure S5**). We then ranked these designs for experimental screening based on a comprehensive set of metrics (**Table S4; Methods**), ultimately selecting the top 101 nanobody designs for PD-L1, 46 designs for IL3, 43 designs for IL20, and 52 designs for BHRF1 for downstream validation. These designs exhibited low CDR sequence identity (<55%; median of ∼30%; **Figure S6**) to any existing sequence in the PDB (Berman et al., 2000) or OAS (Olsen et al., 2022a). Additionally, the tested designs demonstrated significant structural novelty when compared against experimentally resolved complexes in the PDB (**Figure S7; Table S6; Methods**), with low iAlign interface similarity scores (IS<0.47) indicating that the predicted interfaces do not resemble those of any known complexes. Furthermore, the designs selected for experimental testing exhibited substantial structural and sequence diversity within each library (**Tables S7, S8**).

We also used Germinal to design scFvs against two of these targets, PD-L1 and IL3. The resulting designs passed our filtering criteria at rates comparable to nanobody designs and achieved similarly favorable values across key evaluation metrics (**Figure S5**, **S8**, **S9**). Notably, several of the metrics identified as strong predictors of nanobody binding success were based on interface contact area (e.g., LIA, number of hydrogen bonds, and number of interface residues; **Figure S10**), and were thus naturally elevated in scFv designs due to the presence of two binding chains. Therefore, using the information available from the nanobody design rounds, we updated our filters for scFvs to calibrate for the interface area (**Table S3**) and sampled designs for PD-L1 and IL3. These runs yielded 470 PD-L1 and 324 IL3 designs that passed the pipeline filters, of which 48 designs each were selected for experimental validation following stringent filtering and ranking (**Table S5**). Moreover, consistent with our nanobody results, these scFv designs exhibited both CDR sequence and structural novelty. CDR sequence identities to any sequence in the PDB or OAS remained at ∼40% (<52%; **Figure S6**), and binder–target interfaces differed substantially from known complexes in the PDB (IS ≤ 0.2; **Figure S7; Table S6**) as well as from other designs within the generated set (**Tables S7, S8**).

It is worth noting that all input antigen structures to the pipeline are AF3 predictions rather than experimentally determined structures. This demonstrates that Germinal can generate functional binders using predicted rather than experimentally resolved antigen structures, highlighting its potential applicability to targets for which such data are not available (**Supplementary Discussion**). However, for all four targets tested experimentally, the AF3 predictions used as input were themselves validated against known experimental structures (C𝛼 RMSD < 1 Å). As with other hallucination-based methods, Germinal assumes accurate antigen structure conditioning, and design quality will therefore depend on the fidelity of the input model.

### 2.3. Binding affinity characterization of generated antibodies

To experimentally validate nanobody designs produced by Germinal, we employed a split-luciferase assay based on the NanoBiT system (**Figure S11**) (Dixon et al., 2016) to enable efficient scaling and fast iteration across initial design–validation rounds. This assay served as a smaller-scale, simpler alternative to binding-identification assays such as display-based approaches used by other groups (Nabla Bio and Biswas, 2025; Swanson et al., 2025). This system is built on the large luciferase subunit (LgBiT), which we fused to our binders, and the small luciferase subunit (SmBiT) that we fused to the antigen. LgBiT can be complemented by either a high-affinity peptide (HiBiT, 𝐾_𝐷_ = 0.7 nM) or the very low-affinity peptide (SmBiT, 𝐾_𝐷_ = 190 𝜇M). Addition of HiBiT enables direct quantification of LgBiT-tagged binder expression, while addition of an antigen–SmBiT fusion protein allows detection of potential binders. For all four targets, designed nanobodies were fused to LgBiT and encoded in plasmids that were transiently transfected into HEK293 cells. Alongside our designs, we included a validated nanobody against the same antigen (when available) as a positive control, and a nanobody targeting an unrelated antigen as a negative control (**Methods**). To identify promising candidates, binding measurements (SmBiT–LgBiT) were normalized to the corresponding expression measurements (HiBiT–LgBiT), providing a relative measure of binding potential (**Methods**).

Candidates meeting both expression and binding thresholds were advanced to BLI. As a result, we tested 25 of 101 PD-L1 nanobodies, 11 of 46 IL3 nanobodies, 11 of 43 IL20 nanobodies, and 20 of 52 BHRF1 nanobodies for binding to their respective antigens. We observed detectable binding for 7 of 25 PD-L1 designs, 2 of 11 IL3 designs, 4 of 11 IL20 designs, and 5 of 20 BHRF1 designs (**Figure 3A-D; Figure S12**). Crucially, we obtained nanomolar binders for all four antigens tested.

**Figure 3.**
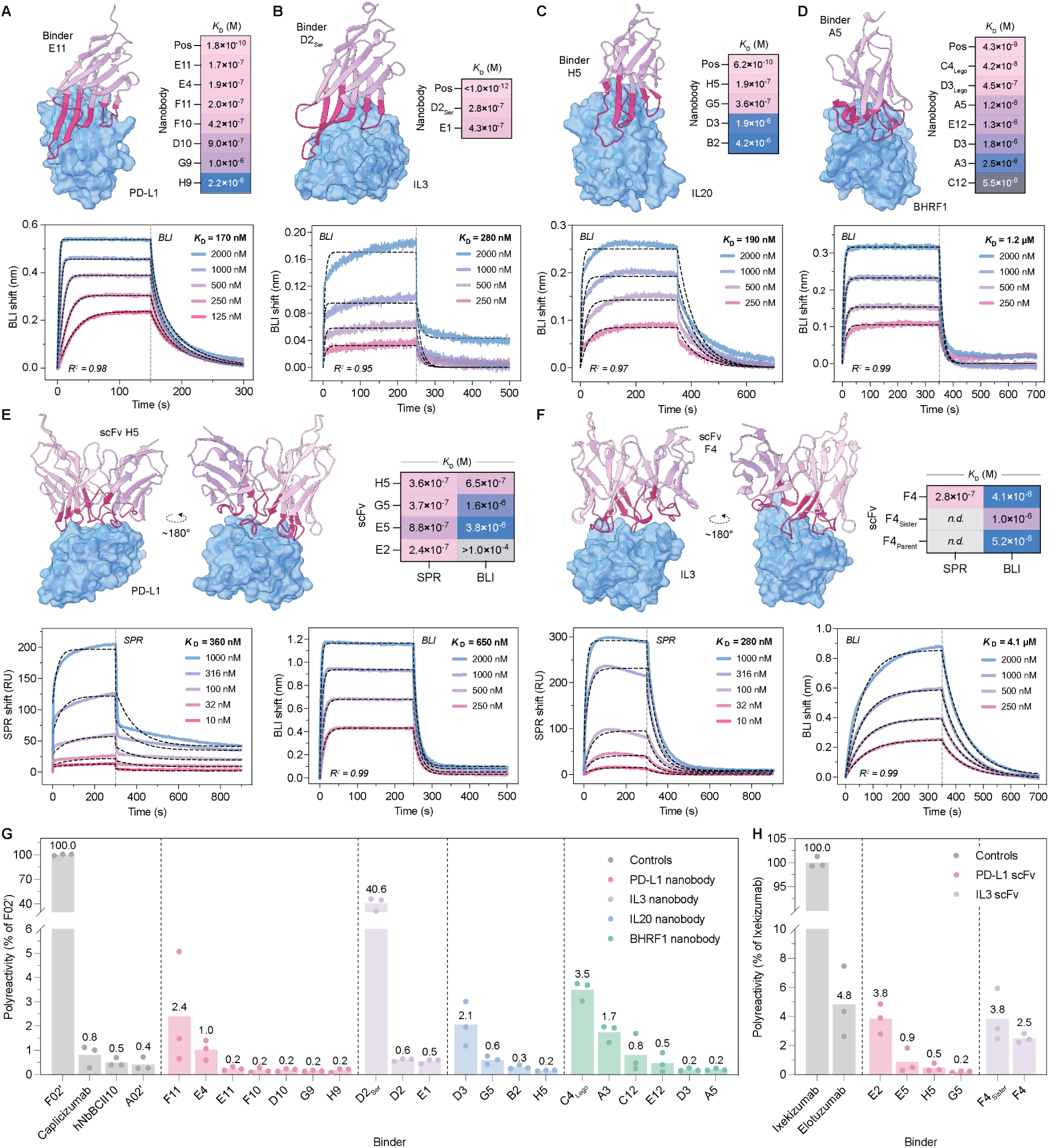
Biophysical characterization of Germinal-designed binders. (**A–D**) For each target, a Germinal-predicted structure of an experimentally validated nanobody (pink cartoon) in complex with the antigen (blue surface) is shown alongside the corresponding BLI sensorgram: PD-L1 (**A**, E11, 𝐾_D_ = 170 nM); IL3 (**B**, D2_Ser_, 𝐾_D_ = 280 nM); IL20 (**C**, H5, 𝐾_D_ = 190 nM); BHRF1 (**D**, A5, 𝐾_D_ = 1.2 μM). Heat maps of BLI-derived 𝐾_D_ values (M) for all nanobody hits for each target are also shown. (**E, F**) Germinal-predicted structures of designed scFv hits against PD-L1 (**E**) and IL3 (**F**), shown from two orientations (180° apart), with heat maps of SPR- and BLI-derived 𝐾_D_ values and representative SPR and BLI sensorgrams for the highest-affinity hit in each panel: PD-L1 scFv H5 and IL3 scFv F4. For all sensorgrams, binding curves are colored by analyte concentration, kinetic fits are shown as black dashed lines, and the derived 𝐾_D_ is shown in bold. All BLI binding were fit with a global 1:1 model unless otherwise specified. (**G–H**) A polyspecificity particle assay was performed to measure the binding of Germinal-designed nanobodies (**G**) or scFvs (**H**) to a polyspecificity reagent (PSR). PSR binding scores are normalized to the polyspecific controls F02’ for nanobodies and Ixekizumab for scFvs. Each design was tested in triplicate (n=3, technical replicates). Dashed vertical lines delineate different targets. *n.d.*, not determined.

Given the success rate of the initial nanobody designs and the consistency of *in silico* predictions between Germinal-designed nanobodies and scFvs, 48 anti-PD-L1 and 48 anti-IL3 scFv designs were reformatted as fragment antibodies (Fabs) and subjected to a preliminary surface plasmon resonance (SPR) affinity screen to identify hits for subsequent biophysical characterization (**Figure S13; Methods**). Fabs were chosen as the screening format due to their closer resemblance to therapeutic IgGs, their clinical relevance, and more predictable retention of binding affinity upon IgG conversion (Steinwand et al., 2014). From this screen, we identified 4 potential PD-L1 binders and 1 IL3 binder.

These hits were subsequently characterized by BLI, both to confirm binding and to establish a baseline for downstream characterization. Of the SPR-validated designs, 3 of 4 anti-PD-L1 designs and the 1 anti-IL3 design had measurable binding activity by BLI (**Figures 3E–F; S14–S15**) and advanced to downstream biochemical characterization, while the remaining anti-PD-L1 design produced inconclusive measurements when tested with BLI. We hypothesize that this is due to discrepancies between SPR and BLI that arise from differences in experimental setup, and thus proceeded only with designs validated by both SPR and BLI. More broadly, despite applying computational filters retrospectively identified from the nanobody screens (**Figure S10**), experimental success rates for scFvs remained comparable to those of the nanobody campaigns, suggesting that these filters may not generalize across antibody formats.

### 2.4. Rescue and optimization of low-expressing designs

After observing low expression levels for a subset of Germinal-generated designs, we explored targeted expression rescue strategies. For the anti-IL3 scFv F4, we evaluated two additional designs that ranked just below the selection cutoff. The first was a parent sequence representing the original Germinal design prior to AbMPNN redesign (F4_Parent_; differing at 6 residues from the hit); the second was a sister sequence sharing the same Germinal seed but carrying a different sampled AbMPNN sequence (F4_Sister_; differing at 2 residues). Both variants expressed at higher levels than the original F4 (**Figure S16**), while retaining comparable or improved binding affinities, and were thus carried forward for downstream characterization (**Figure 3F**).

For the anti-BHRF1 nanobody C4, which consistently yielded insufficient protein for biochemical characterization after purification, we mutated its framework to match the Legobody framework (Wu and Rapoport, 2021) (to yield C4_Lego_; 94.53% sequence similarity), which is nearly identical to those used in synthetic nanobody libraries engineered for high thermal stability (Zimmermann et al., 2018; McMahon et al., 2018). To test whether the legobody framework mutations affect nanobody binding activity, we applied the same re-scaffolding to the BHRF1 nanobody D3 (D3_Lego_), a design that expressed robustly and showed confirmed binding. D3_Lego_ maintained high expression yields and showed improved binding affinity relative to D3, establishing that transfer to the Legobody framework is well-tolerated. Indeed, C4_Lego_ showed improved expression yields in Expi293F cells compared to the original C4 and was found to bind BHRF1 with 42 nM affinity (**Figure S17F; Figure 3D**). Therefore, we proceeded with C4_Lego_ for all downstream experiments.

Separately, the anti-IL3 nanobody D2 contained a solvent-exposed cysteine in CDR3 that likely promoted disulfide-mediated dimerization, as observed by non-reducing gel electrophoresis (**Figures S17F–G**). We hypothesized that this dimerization contributed to D2’s low expression yield and poor-quality BLI measurements (**Figure S12C**). Mutating this residue to serine (D2_Ser_) restored monomeric expression, with improved purity by gel electrophoresis and well-resolved BLI binding curves (**Figures 3B; S12B**). D2_Ser_ was therefore used in all subsequent experiments.

### 2.5. Polyreactivity

We evaluated non-specific binding of our designs using a flow cytometry-based polyspecificity particle assay, in which His- and FLAG-tagged binders are captured on anti-His magnetic beads and assessed for binding to biotinylated solubilized Expi293F membrane proteins (polyspecificity reagent or PSR) (Makowski et al., 2021; Hie et al., 2024). Detection of biotin with streptavidin and FLAG-tag with an anti-FLAG M2 antibody enabled distinction of polyreactive binders from target-specific species, as verified with previously validated positive controls (F02’ for nanobodies and Ixekizumab for scFvs) and non-polyreactive negative controls (A02’ for nanobodies and Elotuzumab for scFvs) (**Figure S18**) (Harvey et al., 2022; Makowski et al., 2021). We found that all Germinal-generated designs exhibited low polyreactivity (<4% of the polyreactive positive control), comparable to the negative control samples (**Figure 3G–H; S19–S20**). This favorable profile was retained in the derivative sequences C4_Lego_ and F4_Sister_. In contrast, D2_Ser_ showed markedly elevated polyreactivity relative to the original D2 design (**Figure 3G**, **Figure S19**), suggesting that although mutation of a CDR cysteine to serine can rescue monomeric expression and target binding, it may also introduce non-specific binding interactions. This result underscores the importance of comprehensive developability assessment beyond binding affinity alone.

### 2.6. Epitope specificity of generated antibodies

To experimentally validate whether Germinal can indeed design epitope-specific antibodies, we determined the structure of a representative anti-PD-L1 scFv (H5) in complex with PD-L1 at 3.9 Å resolution via cryo-EM. Superposition of the experimental structure with the Germinal-predicted model demonstrated close overall agreement, with a global C𝛼 RMSD of 1.25 Å (**Figure 4A**). The predicted model also showed a good fit to the experimental cryo-EM density across all six CDR loops (**Figure 4B,C**), indicating that the predicted backbone geometry is consistent with the experimental map. Comparison at the binding interface confirmed that contacts to the targeted hotspot residues (I37, Y39, E41) were maintained as designed (**Figure 4D**), demonstrating that Germinal can successfully produce designs that bind at the intended epitope.

**Figure 4.**
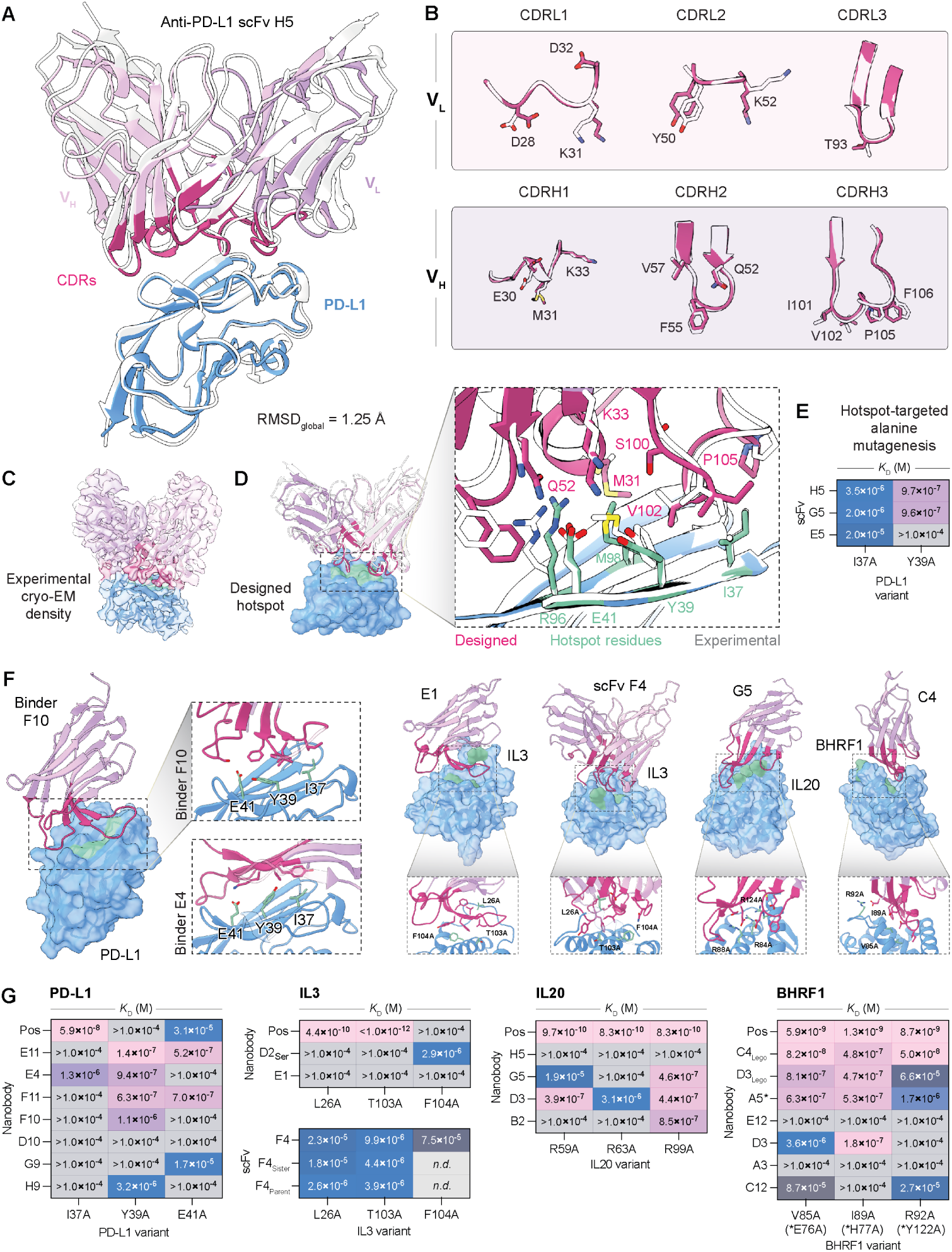
Cryo-EM and mutational mapping of epitopes targeted by Germinal-designed binders. (**A**) Cryo-EM structure of the Germinal-designed anti-PD-L1 scFv H5 (white) overlaid with the AF3-predicted structure (pink), in complex with PD-L1 subdomain I (blue).(**B**) Per-CDR comparison of experimental and predicted loop conformations for V_L_ (top) and V_H_ (bottom), with key interface residues shown as sticks and labeled. (**C**) Experimental cryo-EM density map of the H5 scFv:PD-L1 complex with fitted atomic models overlaid, colored by chain. (**D**) Close-up of the binding interface comparing designed (magenta) and experimental (white) residue conformations, with hotspot residues highlighted in green. (**E**) 𝐾_D_ values (M) for three anti-PD-L1 scFv hits against PD-L1 alanine variants I37A and Y39A. (**F**) Germinal-predicted binding poses for representative binders across all four targets (pink ribbons on blue surfaces), with zoom panels showing binder contacts (≤3.5 Å) at alanine-substituted hotspot residues (green sticks, labeled). (**G**) 𝐾_D_ matrices from alanine mutagenesis at hotspot residues for all four targets (PD-L1: I37A, Y39A, E41A; IL3: L26A, T103A, F104A; IL20: R59A, R63A, R99A; BHRF1: V85A, I89A, R92A). For BHRF1 A5 (denoted by an asterisk (*)), the corresponding hotspot mutations were E76A, H77A, and Y122A. Binding below the limit of detection was assigned 𝐾_D_ >1.0 10^−4^ M. *n.d.*, not determined. All residue numbers refer to the mature protein sequence after signal peptide cleavage (see **Methods** for details).

To further assess whether binders across all four targets engaged their intended epitopes, we measured the effects of alanine substitutions at predicted hotspot residues. For PD-L1, hotspot substitutions I37A and Y39A reduced binding affinity across all three designed scFvs (**Figures 4E, S14**), consistent with the direct contacts observed at these residues in the cryo-EM structure of H5. We performed hotspot mutagenesis across all targets and binder formats, substituting predicted hotspot residues to an alanine: I37, Y39, and E41 for PD-L1; L26, T103, and F104 for IL3; R59, R63, and R99 for IL20; and V85, I89, and R92 for BHRF1 (**Figures 4F, G**) (residue numbers refer to the mature protein sequence after signal peptide cleavage; **Methods**). Substitution of one or more hotspot residues reduced binding affinity by at least two-fold relative to wild-type antigen, with at least one substitution abrogating detectable binding entirely in 17 out of 26 designs (**Figure 4E, G; Figures S14, S15, S21–S24**). Encouragingly, binders with distinct binding modes were differentially affected by individual substitutions, with no single mutation abolishing binding across all designs (**Figure 4F (left), G (left)**). To further confirm that the observed affinity reductions reflected disruption of the binder–antigen interface rather than global destabilisation of antigen folding, monoclonal antibodies with proprietary epitopes (for IL3 and IL20) or binders with overlapping but distinct epitopes (for PD-L1 and BHRF1) were included as positive controls; in each case, control binders retained near wild-type affinity for at least one alanine variant (**Figure 4G**), confirming the structural integrity of the antigen variants. Together, these results demonstrate that Germinal-designed binders engage their computationally specified epitopes with the predicted contacts.

## 3. Discussion

We present an end-to-end pipeline for *de novo* antibody design that achieves nanomolar-to-low-micromolar binders with low-n experimental validation. A scalable, streamlined split-luciferase assay was devised to initially screen and filter designs, efficiently identifying lead nanobodies for subsequent validation by BLI. This pipeline yielded nanobodies against four diverse soluble protein targets, including IL3, for which, to our knowledge, no antibody or other non-natural binder has been reported in the literature or deposited in the PDB aside from its cognate receptor—making it a target for which the pipeline is unlikely to benefit from memorization of known binder conformations. We then extended the computational design methodology to *de novo* scFv antibody fragments, obtaining binders for both PD-L1 and IL3. Importantly, we make our complete methodological details and code publicly available to the scientific community.

Our method differs from previous studies that have leveraged protein language models to guide directed evolution of antibodies toward improved binding affinity, antibody stability, or other therapeutically relevant properties (Hie et al., 2024; Shanker et al., 2024; Widatalla et al., 2024). These efforts have assumed at least weak initial function (e.g., affinity maturation of a weak antibody binder), whereas Germinal designs antibody CDRs completely *de novo* without any starting binder, noting that protein language models may also enable efficient affinity maturation of our generated antibodies. Concurrent with our work, several industry-led efforts (Chai Discovery Team et al., 2025; Nabla Bio, 2025; Latent Labs Team et al., 2025) have described antibody design models and reported double-digit success rates, though their code, model weights, and detailed methods are not publicly available. Open-source pipelines (Swanson et al., 2025; Stark et al., 2025), by contrast, provide greater methodological transparency. mBER, like Germinal, leverages backpropagation-based hallucination with partial structural conditioning for nanobody design, while BoltzGen presents a diffusion-based all-atom generative model capable of designing nanobodies and other binder modalities.However, Germinal directly co-optimizes structural confidence and antibody-likeness, while enabling direct control over binding pose through explicit paratope- and structure-based loss functions. Germinal further couples design to experimental validation of epitope specificity through Cryo-EM structural determination of a binder–antigen complex and epitope-directed alanine mutagenesis, together with a rigorous evaluation of polyreactivity. Separately, Bennett *et al*. recently demonstrated *de novo* design of antibodies with atomic-level structural accuracy using RFDiffusion, though experimental validation required screening tens of thousands of candidates to identify tens of binders (Bennett et al., 2025). While Bennett *et al*. report potentially higher success rates with AF3 filtering, our analysis of their published data (**Supplementary Discussion**) indicates that AF3 filtering would improve their success rate to ~1.2–1.5%. In comparison, Germinal’s experimental success rates represent a several-fold improvement over RFDiffusion even after AF3 filtering. Furthermore, while we did not use ipSAE (Dunbrack, 2025) for scoring our designs, we note that all validated binders except one achieved min_ipSAE scores above the previously suggested threshold of 0.5 (Overath et al., 2025), with the majority exceeding 0.6 (**Figures S25; Table S9**). However, many non-binding designs also exceeded these thresholds, highlighting the difficulty of identifying computational scores that reliably predict binding and generalize across binder formats.

Beyond the initial design pipeline, we note that computational *de novo* antibody design will continue to benefit from rational optimization strategies rooted in traditional protein engineering and biochemistry. In this work, we employed three such approaches: directed framework mutations (via re-scaffolding nanobody designs onto the Legobody framework) to rescue low-expressing candidates (BHRF1 C4_Lego_), mutating a solvent-exposed cysteine to recover monomeric expression (IL3 D2_Ser_), and recovering sister and parent binder variants from the design trajectories (IL3 scFv F4). These represent optional, modular steps that can be layered onto the core pipeline to expand the pool of functional binders or improve the properties of initial hits.

Several aspects of the Germinal model and sampling pipeline also present clear opportunities for future development. First, the method’s performance is fundamentally constrained by the confidence and accuracy of AF-M predictions. Additionally, the need for structure prediction and network backpropagation at each iteration requires extensive inference-time sampling to generate successful designs, rendering it computationally intensive (see Computational resources in **Methods**). Future improvements in optimization algorithms and integration with other generative approaches could significantly reduce this computational burden. Memory constraints also limit the method to smaller protein targets, though most epitopes can be successfully targeted by truncating large proteins to the domain of interest. More broadly, to support wider adoption of Germinal, future work could make the pipeline fully open by substituting components that currently carry restrictive licenses, such as IgLM and AF3, with open-source alternatives including AbLang (Olsen et al., 2022b) and ProteniX (Team et al., 2026).

Similar to other methods (Swanson et al., 2025; Zambaldi et al., 2024; Nabla Bio, 2025), Germinal is cur-rently restricted to favorable protein epitopes, limiting success rates for hard-to-target surfaces (**Supplementary Discussion**). We note that our results demonstrate the ability to target specific epitopes rather than any arbitrary ones, and thus performance will vary depending on the biophysical properties of the selected surface. Similarly, given that AF-M can only model canonical protein amino acids, important classes of targets such as glycans, small molecules, and other non-protein antigens remain beyond its current scope. Further, although we demonstrated that Germinal can produce high-confidence designs against predicted antigen structures, all experimentally validated targets were compared against existing experimentally-determined structures. Therefore, for antigens such as MCF2, a target with no experimentally determined structure (**Supplementary Discussion**), a systematic evaluation of how design success rates depend on antigen model quality remains an important open question. Finally, while Germinal’s loss functions are designed to steer designs toward CDR loop-dominated binding poses, the optimization yields a distribution of interface conformations, and a subset of designs was observed to engage the target through secondary-structure-rich contacts. Interestingly, some of these designs tended to exhibit stronger binding affinities, but also showed higher signal in the polyspecificity assay (**Figure 3G–H**). Although the sample size is too small to draw definitive conclusions, this observation warrants caution when prioritizing designs with secondary-structure-rich interfaces, particularly those involving partial framework regions.

The ability to computationally design epitope-targeted antibodies with high success rates opens transformative possibilities across biotechnology and medicine. Germinal could unlock targeting of previously inaccessible epitopes, including intracellular domains, specific conformational states, and highly conserved regions that fail to elicit natural immune responses. Rapid antibody design could compress therapeutic antibody identification from weeks to days, improving our response to novel pathogens or rapidly evolving disease targets. As our generative pipeline becomes more efficient, systematic design campaigns could produce antibodies against entire proteomes, vastly expanding our repertoire of affinity reagents and therapeutic molecules.

## Supporting information

supp_file1_plasmidmaps

supp_file2_all_seq_for_binders

supp_file3_bli_fits

## 4. Acknowledgments

We thank Connor C. Call, Julia Kazaks, Brian Plosky, and David Li for helpful discussions and support with the manuscript. L.S.M acknowledges support by the Stanford Bio-X fellowship, the Stanford Graduate Fellowship, the Stanford University Sarafan ChEM-H Chemistry–Biology Interface training program and the Stanford Interdisciplinary Graduate Fellowship. T.W. acknowledges support by the NSF Graduate Research Fellowship and the Stanford Graduate Fellowship. X.Z. is supported by the Stanford Interdisciplinary Graduate Fellowship affiliated with ChEM-H. B.L.H. acknowledges funding support from Arc Institute, the Gates Foundation, Stanford Institute for Human-Centered Artificial Intelligence (HAI) Hoffman-Yee Research Grants, V. Gupta, and R. Tonsing. X.J.G. acknowledges funding support from the National Institutes of Health (NIH DP2EB035891), Longevity Impetus grants, Stanford Bio-X Interdisciplinary Initiatives seed grant program, the Rosenkranz Foundation, the Hevolution Foundation, and Genscript Life Science Research Grant.

## 5. Author Contributions

L.S.M., J.N.W., T.W., B.L.H., and X.J.G. conceived the study. L.S.M., C.L.D., J.N.W., H.D., X.Z. designed experiments. B.L.H. and X.J.G. supervised the project. L.S.M. and J.N.W. developed the algorithms and pipeline. L.S.M., J.N.W., and T.W. contributed to the codebase and performed computational analysis of designs. L.S.M. and J.N.W. ran design campaigns and selected designs for experiments. L.S.M., C.L.D., J.N.W., and H.D. cloned plasmids used in this work. L.S.M., C.L.D., H.D. and X.Z. ran luciferase assays. L.S.M., C.L.D., and H.D. performed protein purification and ran BLI validation and alanine mutagenesis experiments. L.S.M. and C.L.D. performed polyspecificity particle assays. L.S.M. and C.L.D. performed the alanine mutagenesis experiments. H.D. performed differential analysis on computational and experimental data. J.L.M. and B.R. determined the cryo-EM structure. L.F. supervised cryo-EM structure determination. L.S.M., J.N.W., C.L.D., T.W., B.L.H., and X.J.G. wrote the manuscript.

## 6. Competing Interests

B.L.H. acknowledges outside interest in Arpelos Biosciences and Genyro as a scientific co-founder. X.J.G. is a cofounder and serves on the scientific advisory board of Radar Tx. L.S.M, J.N.W., C.L.D., H.D., T.W., X.Z., B.L.H, and X.J.G are named on a provisional patent application applied for by Stanford University and Arc Institute related to this manuscript.

## 7. Code and Data Availability

We make code for running the pipeline available at http://www.github.com/SantiagoMille/germinal. We make the following files available as supplementary data:

- **Data S1:** Template plasmid maps used in this work. supp_file1_plasmidmaps.xlsx
- **Data S2:** Designed sequences per target and their corresponding names. supp_file2_all_seq_for_binders.xlsx
- **Data S3:** Derived 𝐾_𝐷_s and associated 𝑅^2^ values from BLI experiments. supp_file3_bli_fits.zip

## A. Methods

### A.1. Germinal algorithm

Germinal performs gradient-based optimization on the joint landscape defined by AF-M and IgLM objectives, merging their gradients into a single update direction. Following previous work (Pacesa et al., 2025), design proceeds through three phases (logit, softmax, and semi-greedy optimization) that progressively anneal continuous logits into one-hot sequence representations (**Figure 2A**). Non-interface CDR residues are then redesigned during sequence optimization using an antibody-finetuned sequence design model. Finally, designs undergo filtering and ranking to select top candidates for experimental testing.

#### A.1.1. Input preparation

To run the design pipeline, we first defined the antigen structure and the target epitope. To reduce computational cost, multi-domain antigens were restricted, whenever possible, to the independently folding domain containing the target epitope. For PD-L1, we used the V-like domain, while for IL3, IL20, and BHRF1, we used the full mature protein sequence excluding signal peptides. Antigens were predicted using AF3 (Abramson et al., 2024) and validated to fold with high confidence and low RMSD when aligned to the experimental structure (RMSD < 1 Å), if available. This procedure was also applied to new antibody frameworks.

Epitope residues on the target were specified in an input settings file, as were CDR lengths for the antibody framework (identified via AbNumber using IMGT numbering) (Dunbar and Deane, 2016; Lefranc et al., 2003). These lengths, together with user-defined starting points, were used to calculate CDR positions. For antigens with known binders (e.g., natural receptors), the epitope corresponding to the reference biologically relevant interaction was designated as the preferred epitope and the one to be consistently tested first. This choice was motivated by: 1) targeting natural interaction interfaces is often of direct therapeutic interest, and 2) in preliminary unconstrained runs, these regions emerged most frequently among successful designs. However, for some targets the known epitope was not the most productive and an alternative was used instead. See **Supplementary Discussion** for a retrospective analysis of epitope-dependent design success rates across targets.

#### A.1.2. Initialization

Before the design stage, the antibody framework and antigen structures are merged into a unified PDB containing two chains, which serves as the starting complex. The sequence position-specific scoring matrix (PSSM) is initialized by setting framework positions to their one-hot encodings and CDR positions to one of three different initializations depending on user choice: (i) uniform random noise, (ii) Gumbel noise centered around the template framework sequence, or (iii) zeros. This flexibility allows users to control the trade-off between broad sampling and guided starting points. Gumbel initialization was adopted as the default based on empirical performance. This initialization, together with the starting conditions and random seed, determines the diversity of the design trajectories.

During AF-M initialization, the antigen and antibody framework structures are also supplied as templates to AF-M. Importantly, only the framework regions (FRs) of the antibody structure are used as templates, with all CDR residues masked in the input feature matrices (e.g., atom positions, side chains, sequence). This ensures that AF-M provides structural guidance based only on stable framework features without biasing the optimization of CDR loops to the natural CDRs of the framework structure.

#### A.1.3. Sequence bias

To bias sequences toward a chosen framework, we use a bias weight to modulate between framework preservation and exploration. This term is applied to framework region (FR) residues only (i.e., all residues other than CDRs). Specifically, at every iteration, this bias is added to the logits corresponding to the original framework sequence before the current binder is used as input to AF-M and IgLM, essentially shifting the probability distribution to favor the original framework sequence. Unlike hard-fixing, this approach still permits mutations in FR positions if gradients strongly favor them. Based on parameter sweeps (**Figure S26**), we selected a value of 10, which minimized framework mutations while allowing enough flexibility for confidence optimization, and hence successful trajectories.

#### A.1.4. Design stage phases

- **Logits**: In this phase, the sequence is represented as continuous logits for each position and amino acid type. Gradient-based optimization is performed directly on these logits using the combined gradients from AF-M (structural confidence) and IgLM (sequence naturalness). The logits allow for smooth optimization in continuous space while maintaining differentiability.
- **Softmax**: Here, the sequence logits of the binder are normalized to sequence probabilities using the softmax function. Gradients are similarly merged and applied to update this representation of the sequence.
- **Semi-greedy optimization**: In this final phase, the softmax probabilities from the previous stage are used to sample mutations, and mutations with the best loss are fixed.

#### A.1.5. Optimization strategy

Gradients from AF-M and IgLM are merged at each iteration of the logit and softmax phases using either a weighted sum, Multiple Gradient Descent Algorithm (MGDA) (Sener and Koltun, 2019) or a weighted version of PCGrad (Yu et al., 2020) (see ***Gradient merging***). In the semi-greedy phase, optimization shifts from continuous gradients to discrete sequence updates. At each of the 10 semi-greedy iterations, five candidate sequences are sampled, and the one that best balances AF-M structural confidence with IgLM sequence likelihood—calculated as a weighted sum of the AF-M losses and IgLM log-likelihood—is accepted.

For all three stages, the IgLM weight is adjusted according to a four-step schedule. During the logits phase, the IgLM weight is linearly annealed from 𝑣_1_ to 𝑣_2_. In the softmax phase, the weight is set to 𝑣_3_. Finally, in the semi-greedy phase, the weight is set to 𝑣_4_. This annealing schedule empirically provided more robust performance than fixed IgLM weights, and is the default setting in Germinal.

Additionally, because Germinal does not include a discrete one-hot optimization phase, we check for sequence convergence at the end of the softmax phase. Specifically, for each CDR position we: (i) compute the maximum probability across all 20 amino acids and (ii) average these values across all CDR positions. The resulting value represents the average positional certainty in the most likely amino acid per position. To ensure that the model is confident enough in the sequence to justify discretization, only sequences exceeding an empirically determined threshold (0.1) for this metric are advanced to the semi-greedy stage.

At the end of each phase, the current best design is evaluated against AF-M confidence metrics (pLDDT > 0.8, ipTM > 0.68 and iPAE < 0.35) and designs advance to the next phase only if they satisfy these thresholds. At the end of the design stage, remaining sequences are co-folded with AF3 for independent evaluation prior to ***Sequence optimization***. Designs must satisfy strict AF3 confidence thresholds (ipTM > 0.75, PAE < 8, pTM > 0.85, and pLDDT > 0.85), exhibit a CDR interface percentage greater than 60%, and be predicted to bind the specified epitope to proceed.

Importantly, the final AF-M loss used in Germinal is computed as a weighted sum of multiple structural and interface metrics. A full list of the loss terms and their weightings, as well as biases added during optimization are shown below (see ***Loss and bias terms***). Additionally, during each iteration, AF-M is run using three recycles, with the final recycle taken as the prediction for each iteration.

#### A.1.6. Gradient merging

Central to Germinal is the gradient merging algorithm, which reconciles updates from AF-M and IgLM objectives. Before merging is applied, gradients from both models are normalized to their respective ℓ_2_-norms. This ensures optimization of the two objectives is not biased due to magnitude differences between the two gradients. The resulting gradient is then re-normalized to have unit ℓ_2_-norm again and scaled by the effective sequence length (the number of non-zero positions in the AF-M gradient) as originally done by ColabDesign. Three strategies were explored used for gradient merging:

- **Weighted Sum**: gradients are merged by a direct weighted sum where the AF-M weight is fixed to 1, and the IgLM weight 𝜆 is set by a user-defined parameter.

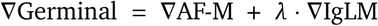

- **Weighted PCGrad**: As described in (Yu et al., 2020), PCGrad mitigates destructive interference between gradients by resolving conflicts in direction. Specifically, one gradient is projected onto the orthogonal complement of the other if the dot product is negative, thereby preventing cancellation of progress along critical dimensions. In our implementation, the previously normalized vectors are scaled by a user-defined weight before applying PCGrad.
- **Multiple Gradient Descent Algorithm**: MGDA (Sener and Koltun, 2019) formulates gradient merging as a multi-objective optimization problem, where the goal is to compute a Pareto-optimal descent direction. Concretely, MGDA solves for the convex combination of AF-M and IgLM gradients that minimizes the squared norm of the merged update:

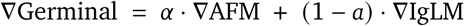

with 𝛼 ɛ [0, 1] chosen to balance objectives without privileging one a priori. This is achieved by solving a small quadratic program at each step. In practice, MGDA is more computationally intensive than PCGrad or weighted sums, but it provides a principled way to balance objectives when their contributions vary dynamically during optimization.

#### A.1.7. Loss and bias terms

Below, we list all biases, weights, and custom loss terms utilized in Germinal.

We use the following biases:

- **Sequence bias:** Higher values enforce greater similarity to the framework sequence.
- **IgLM scale:** Scaling factor applied to normalized IgLM gradients relative to AF-M gradients.

We use the following ColabDesign loss functions (weights):

- **pLDDT loss:** Emphasizes high per-residue confidence in AF-M predictions.
- **ipLDDT loss:** Emphasizes high per-residue confidence at the predicted interface.
- **iPAE loss:** Penalizes high alignment error between predicted and target structures at the interface.
- **PAE loss:** Penalizes high alignment error of the predicted complex.
- **ipTM loss:** Favors predictions with high predicted TM-score, encouraging correct geometry at the interface.
- **pTM loss:** Favors predictions with high predicted TM-score, encouraging correct overall geometry.
- **Inter-protein contacts:** Ensures that the designed antibody has enough contacts to the antigen chain within a certain distance. The number of contacts and distance can be specified by the user. If a hotspot is provided, the loss favors contacts with the specified positions.
- **Intra-protein contacts:** Encourages each residue in the designed sequence to form a sufficient number of contacts with other residues within the same protein. The user can specify both the minimum number of contacts required and the distance threshold used to define a contact.

We use the following custom loss functions:

- 𝛼 **-helical loss:** This loss is a variant of the helix loss from (Pacesa et al., 2025), restricted to CDR positions. We discourage 𝛼-helical geometry within the CDRs by penalizing {𝑖, 𝑖+3} pairs whose predicted distances fall in the 2-6.2 Å range characteristic of 𝛼-helices. Using AlphaFold’s distogram, we compute a binary cross-entropy penalty per residue pair that is large when probability mass lies inside this window and small otherwise. We also apply a positional mask to restrict evaluation to CDR residues, averaging the resulting penalties. Minimizing this loss reduces helix-like contacts in the CDRs and biases designs toward flexible, loop-like conformations.
- 𝛽 **-strand loss:** Analogous to the helix term, this loss suppresses 𝛽-strand–like geometry within the selected CDR positions by penalizing {𝑖, 𝑖+3} pairs whose predicted distances fall in a 𝛽-like window (9.75–11.5 Å). Similar to the helix loss, we compute a binary cross-entropy penalty per pair with respect to this window using the distogram. To restrict this loss to CDR residues, we apply a positional mask that includes all CDR including the position directly before and directly after each CDR. We sort the probability of each pair constituting a 𝛽-strand and the top three most likely pairs were averaged as the final 𝛽-strand loss. Minimizing this loss reduces 𝛽-strands in the targeted regions, complementing the 𝛼-helix loss to favor loop-like conformations.
- **Paratope loss:** We developed a paratope loss to achieve two goals: (1) encourage the CDR residue to bind to the specified epitope and (2) discourage the framework binding to the target. For the first goal, we define a loss *L*_CDR_, which operates on two sets of residues: target_id, which includes all specified hotspot residues on the target (or all residues in the target if no hotspot is specified), and pos_id, which include all CDR residues. For each pair of residues between target_id and pos_id, we calculate a categorical cross-entropy (CCE) loss using the distogram distance probabilities and a user-defined distance threshold. We then average the top 𝑘 closest predicted contacts for each epitope residue, across all residues in target_id to yield a final scalar loss. For the second goal, we calculate the same loss with different sets of residues. Here, we set target_id to all target residues and define fw_id as the set of framework residues on the antibody. Following the same loss calculation yields *L*_framework_ (**Figure 2B**). We then define the final paratope loss as:

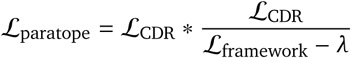

where 𝜆 is a user-defined parameter and is referred to as an “offset” value in the codebase. Note that both _CDR_ and *L*_framework_ will decrease when the corresponding region gains more contacts with its target. Thus, the combined term *L*_paratope_, which is proportional to *L*_CDR_ and inversely proportional to *L*_framework_, increases when framework contacts are abundant and decreases when CDR residues are predicted to bind to the specified epitope, steering contacts toward the intended CDR pose via minimization of the loss.

#### A.1.8. Sequence optimization

Design candidates that cleared initial filters are passed through AbMPNN (Dreyer et al., 2023; Dauparas et al., 2022), a variant of ProteinMPNN fine-tuned on OAS (Olsen et al., 2022a) and SAbDab (Dunbar et al., 2014). AbMPNN was used to generate 40 sequence variants per design, of which the top 4 (by AbMPNN log-likelihood) were retained for re-folding with AF3 and final filtering. Optimization was restricted to CDR residues with all heavy atoms >3 Å from the antigen, preserving key paratope–epitope contacts while allowing AbMPNN to improve the stability of the surrounding loop scaffold. This is consistent with the improvement in binder_score, a PyRosetta energy term for which lower values indicate greater predicted stability, observed across AbMPNN-redesigned sequences relative to their parent designs (**Table S1**). Beyond stability, this step also serves as a practical means of diversifying the candidate pool, as the AbMPNN-redesigned sequences exhibit only modest changes in other key scores (**Figure S27**). Since each successful trajectory effectively seeds multiple distinct yet high-quality candidate sequences, it increases the yield of testable designs without additional sampling and can de-risk a given structural proposal, as different sequences may exhibit distinct developability properties that increase the likelihood that at least one will have a favorable profile. This process also implicitly acts as a self-consistency check, as designs from the same seed frequently scored highly and proved functional when tested experimentally. In future iterations of the pipeline, this self-consistency behavior could serve as an additional ranking criterion.

Finally, while the sequence optimization step may not be strictly required in all cases — for example, IL3 scFv F4 was experimentally successful both with and without AbMPNN optimization (**Figure S15**) — broader experimental validation will be needed to determine how generally it can be omitted.

#### A.1.9. IgLM Implementation

Slight modifications were made to the original implementation of IgLM (Shuai et al., 2023) in order to incorporate the PSSM sequence representation during design. All inputs to IgLM are one-hot amino acid sequences; at each iteration, we convert the raw PSSM to a probability distribution using a softmax operation with a temperature of 0.6, then use a straight-through estimator to obtain discrete sequence representations. We condition each sequence (split into VH and VL for scFvs) with a chain token (<HEAVY> or <LIGHT>) and a species token (e.g., <HUMAN>) (Vincke et al., 2009). Loss is calculated by taking the mean negative log-likelihood over the entire sequence, which is then backpropagated to the raw PSSM to yield the update gradient for the iteration.

#### A.1.10. Design filtering

For filtering, designs were evaluated against a set of stringent thresholds derived from structural prediction confidence scores and PyRosetta-derived metrics (Chaudhury et al., 2010). Only sequences passing all thresholds advanced to the next stage. Seeking to utilize an independent structure prediction model from AF-M, and given its superior accuracy in modeling antibody-antigen complexes (Hitawala and Gray, 2024), we selected AlphaFold 3 (AF3) (Abramson et al., 2024) as the final co-folding model. All AF3 predictions were performed without structural templates and only using MSAs for the target. The metrics and thresholds applied for filtering are detailed in **Table S2**.

#### A.1.11. Design ranking

After filtering, we proceed to an optional, yet recommended, ranking step, to reduce the number of designs to be experimentally validated. Ranking is conducted using a Borda count–based ranking algorithm, which integrates multiple structural and sequence-based metrics, each weighted by relative importance (**Table S4**).

Hard threshold filtering followed by Borda ranking ensures the top high-confidence, well-structured, and framework-preserving candidates are chosen. Retrospective analysis further highlighted the importance of several metrics (**Figure S10**).

#### A.1.12. Computational resources

All sampling was performed on NVIDIA H100 GPUs. Although exact runtimes vary heavily by antigen, typical sampling runs require 200–500 H100-GPU hours to yield 200–400 designs that pass pipeline filters.

#### A.1.13. Structural novelty analysis

To determine the structural novelty of designed binder–target interfaces, we first performed a broad structural search using Foldseek-Multimer (van Kempen et al., 2024), which enables fast and sensitive comparison of large protein complexes. Specifically, we used easy-multimersearch to query every BLI- or SPR-verified hit against the PDB for each target:

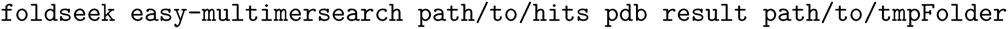

PDB structures deposited after the AlphaFold-Multimer training cutoff date were excluded. To quantify interface-level structural similarity against all Foldseek hits, we additionally computed Interface Similarity (IS) scores using iAlign (Gao and Skolnick, 2010), which performs structure-based comparison of protein–protein interfaces. For each verified hit, iAlign was run against all multi-chain Foldseek matches (with the same post-cutoff exclusions applied).

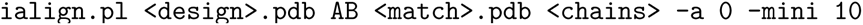

For each campaign, the match with the highest IS-score was selected and the design complex was aligned to the matched PDB structure by superposition of the target chain for visualization.

#### A.1.14. Sequence novelty analysis

To determine the sequence novelty of the designs, the MMseqs2 toolkit (Mirdita et al., 2019) was used to assess the sequence similarity to all entries in the PDB (Berman et al., 2000) and Observable Antibody Space (OAS) (Olsen et al., 2022a). The MMseqs2 command used is:

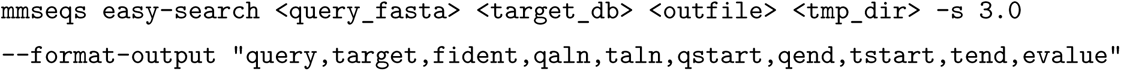

The pairwise alignments are post-processed to compute CDR-specific identity by extracting the alignment positions corresponding to the IMGT-derived (Lefranc et al., 2003) CDR positions and computing the number of matching positions divided by total CDR positions.

#### A.1.15. Structural and sequence diversity analysis

To assess the structural diversity of designed binder–target interfaces within each campaign, we computed the Interface Similarity (IS) scores using iAlign (Gao and Skolnick, 2010) between all pairs of designs within each campaign. Specifically, for each campaign, every design complex was compared against every other on chains A and B (target and binder), and the pairwise IS-scores were recorded. This was done for all tested designs as well as verified hits. Similarly, to assess the sequence diversity of designed binders, we computed CDR sequence identity between all pairs of designs within each campaign, and between all verified hits within each campaign.

#### A.1.16. Therapeutic Nanobody Profiler analysis

To evaluate designs with the Therapeutic Nanobody Profiler (TNP), the Github repository found at github.com/ oxpig/TNP was utilized. Since there is no Github to our knowledge for the Therapeutic Antibody Profiler (TAP), the web server was utilized (https://opig.stats.ox.ac.uk/webapps/sabdab-sabpred/sabpred/tap). The thresholds for green, amber, and red (shown as blue, purple and pink, respectively, in **Figure S3, S4**) risk levels are derived from the automatic assignment from the profiler. Humanness scores were calculated using the promb package (https://github.com/MSDLLCpapers/promb) following methodology proposed in BioPhi.

### A.2. Experimental characterization

#### A.2.1. Framework selection

To identify a robust scaffold for CDR grafting, we evaluated both nanobody and scFv frameworks for their expression and tolerance to CDR replacement. For nanobodies, we screened three frameworks: the humanized nanobody hNbBCII-FGLA, its camelid predecessor cAbBCII, and an alpaca nanobody that has been used in prior in-house experiments. These scaffolds were chosen due to their known stability and successful use as universal grafting frameworks (Vincke et al., 2009). For scFvs, we tested an anti-IL2 scFv from our collection (Fab-F5111) (Trotta et al., 2018), an scFv version of trastuzumab (Byun et al., 2023), and a previously reported scFv framework (FW1.4) (Borras et al., 2010). All nanobody scaffolds were grafted with PD-L1-nb (Zhang et al., 2017) CDRs (PDB: 5JDS), and all scFvs were grafted with F5111 CDRs to assess the effect of framework on binding and expression activities. Among the nanobody grafts, only hNbBCII and cAbBCII expressed robustly and retained binding activities (**Figure S1**). We selected hNbBCII for downstream work based on its prior successful use (Bennett et al., 2025). Among the scFv grafts, only the trastuzumab scaffold yielded appreciable expression, but none of the scFv grafts showed a significant binding signal. We retained trastuzumab given its previous successful use in other computational antibody design pipelines (Bennett et al., 2025).

Notably, CDR lengths were not modified for any design and were instead kept identical to those of the original framework. This decision was motivated by test experiments in which most designs with non-native CDR lengths failed to express, and by previous work showing that grafting CDR sequences onto a mismatched framework substantially reduces both thermostability and binding affinity (Nakakido et al., 2024; Kinoshita et al., 2022). In nanobodies, the lengths of CDR1 and CDR2 are highly conserved across structures deposited in the PDB (Mitchell and Colwell, 2018). Whilst the length of CDR3 is variable, its length carries a structural consequence: whether or not CDR3 makes intramolecular contacts with the framework depends on both CDR3 length and framework identity (Nakakido et al., 2024; Kinoshita et al., 2022).

#### A.2.2. Plasmids and cloning

Constructs were cloned by PCR methods using either NEBNext High-Fidelity Polymerase (New England Biolabs; #M0544) followed by Gibson assembly of fragments (New England Biolabs; #E2621S), or Phusion Flash High-Fidelity Master Mix (Thermo Fisher; #F548L) followed by In-Fusion fragment assembly (Takara Bio). In all cases, constructs were assembled in a three-piece In-Fusion strategy, in which the mammalian expression vector backbone was split into two fragments by PCR with pUC-ori primers, and homology regions were designed to overlap with the synthetic DNA fragments obtained from Twist Biosciences.

DNA sequences of designed binders and targets (PD-L1, IL3, IL20, and BHRF1 (**Figure 1C**)) were ordered from Twist Biosciences with Gibson cloning adaptors for insertion into a pCMV mammalian expression vector. Target proteins were cloned with their native N-terminal signal peptide, with the exception of BHRF1, which was cloned with the IL2 signal peptide (MYRMQLLSCIALSLALVTN). All designed binders were cloned with the IL2 signal peptide. For split-luciferase assays, all target proteins were fused to a C-terminal SmBit peptide and His tag, while all designed binders were fused to a C-terminal LgBit protein. For BLI experiments, target proteins were cloned with a C-terminal AviTag (GLNDIFEAQKIEWHE) and His tag, except for IL3, which was cloned in both C-terminal and N-terminal AviTag formats due to the proximity of its termini to the target epitope; designed binders were fused to a C-terminal His tag only. For polyspecificity assays, nanobodies were cloned with an N-terminal FLAG tag (DYKDDDDK) and a C-terminal His tag, while Fabs were cloned with a C-terminal FLAG tag on the light chain and a C-terminal His tag on the heavy chain. For Cryo-EM experiments, target antigens were cloned with a C-terminal His tag only, and PD-L1 Fab H5 was cloned with a C-terminal His tag on the heavy chain only. Plasmids were propagated in Turbo Competent Cells (New England Biolabs; #C2984) for high-yield transformation and outgrowth, and purified using standard plasmid preparation protocols. All constructs were validated by whole-plasmid Nanopore sequencing. A full list of tested binder sequences is provided in **Data S2**.

#### A.2.3. Expi293 protein expression

All target proteins for split-luciferase assays and lead designed binders were expressed in Expi293F cells (Thermo Fisher; #A14635). For BLI experiments, target proteins possessing an AviTag were expressed in Expi293F in the presence of a BirA enzyme, enabling biotinylation of targets in vivo. Cells were maintained in a 2:1 mixture of FreeStyle 293 Expression Medium (Thermo Fisher; #12338018):Expi293 Expression Medium (Thermo Fisher; #A1435102) and grown at 37 °C with 8% (v/v) CO2 and shaking at 125 rpm. After ≥ 3 passages, cells were diluted to 2 106 cells/mL and grown for 20-24 h, when they were diluted to 3 106 cells/mL for transfection in a final volume of 30-50 mL (designed binders) or 100-250 mL (targets).

Transfection mixtures were prepared with the following components per mL of expression culture: 0.5 𝜇g midi-prepped DNA, 1.3 𝜇L FectoPRO transfection reagent (Polyplus; #101000007), and 100 𝜇L expression media. After incubating for 10 min, the transfection cocktails were added to expression cultures. Cultures were immediately supplemented with glucose (0.45% (w/v) final concentration) and valproic acid (2 mM final concentration), and grown for 4-5 days post-transfection, when cell viability fell below 50%. Secreted protein was harvested from cells by centrifugation at 4000 × g at 4 °C, followed by filtration through a 0.22 𝜇M syringe filter (Thermo Fisher; 723-2520). 10× phosphate buffered saline (PBS; 1.37 M NaCl, 27 mM KCl, 100 mM Na2HPO4, 18 mM KH2PO4, pH 7.4) and 1.63 M NaCl were added to filtered supernatant each to a final dilution of 1:10 and, if necessary, supernatants were pH-adjusted to pH 7.6 using 1 M Tris-HCl pH 9.0.

#### A.2.4. Immobilized metal ion affinity chromatography purification of His-tagged proteins

All His-tagged proteins were purified using HisPur Cobalt resin (Thermo Scientific; #89965). Cobalt resin was washed twice in binding buffer (high-salt phosphate buffered saline [HS-PBS; 300 mM NaCl, 2.7 mM KCl, 10 mM Na2HPO4, 1.8 mM KH2PO4, pH 7.4], 10 mM imidazole) before adding to clarified supernatants and incubating for 1 h at 4 °C with end-over-end rotation. Purification mixtures were transferred to chromatography columns for gravity flow purification. Resin was first washed with binding buffer, followed by wash buffer (HS-PBS, 30 mM imidazole), before proteins were eluted with elution buffer (HS-PBS, 200 mM imidazole). Eluants were immediately buffer exchanged into PBS pH 7.4 and concentrated using centrifugal concentrators with 5 or 10 kDa molecular-weight cut-off (Millipore; #UFC901024), by four rounds of centrifugation (4,000-15,000 × g, 10 min, 4 °C) followed by resuspension in PBS pH 7.4. Protein concentrations were determined from A_280_ using their extinction coefficients predicted from ExPASy ProtParam. Protein purity was assessed by SDS-PAGE (**Figure S17**, **Figure S16**), flash-frozen in liquid nitrogen, and stored at −80 °C.

#### A.2.5. Size-exclusion chromatography purification of antigens

For antigens whose Co-NTA eluates contained visible contaminants by SDS-PAGE, preparations were further purified by size-exclusion chromatography on an ÄKTA pure FPLC system (Cytiva) equipped with a Superdex 200 Increase 10/300 GL column (Cytiva) pre-equilibrated in degassed PBS pH 7.4. A 0.5 mL sample was injected per run and antigen purity and monodispersity within peak fractions were confirmed by SDS-PAGE (**Figure S28A**). Fractions containing target species were pooled (**Figure S28B**), concentrated, flash-frozen in liquid nitrogen, and stored at −80 °C.

#### A.2.6. HEK293T protein expression

Transient transfections were performed on all of the binder plasmids using jetOPTIMUS DNA transfection Reagent (Avantor) in 24-well plates, with each plate incorporating internal controls. When available, validated binders (PD-L1 binder, KN035 [PDB: 5JDS]; BHRF1 binder, GDM_BHRF1_35 (Zambaldi et al., 2024)) were included as positive controls. Binders not associated with a given target (a PD-L1 binder, KN035, for IL20 and BHRF1; a TNF𝛼 binder [PDB: 5M2J] for PD-L1 and IL3) were included as negative controls. This layout ensured that every plate contained both experimental and control conditions for consistent data normalization and comparison. Each binder plasmid was transfected in two biological replicates, with each replicate consisting of 450 ng of binder plasmid and 5 ng of CMV-driven mCherry plasmid (as co-transfection marker). A total of 990 ng binder plasmid DNA was added to 110 𝜇L of transfection buffer with co-transfection marker mixed. 0.88 𝜇L of transfection reagent was then added, well mixed, and incubated for 9 minutes. Following incubation, 50 𝜇L of the transfection complex was added dropwise to each well of Human Embryonic Kidney (HEK) 293 cells (ATCC CRL-1573) at 80-90% confluency. The cells were cultured in Dulbecco’s Modified Eagle Medium (DMEM), supplemented with 10% fetal bovine serum (Fisher Scientific; #FB1299910), 1 mM sodium pyruvate (EMD Millipore; #TMS-005-C), 1× penicillin–streptomycin (Genesee catalog, #25-512), 2 mM l-glutamine (Genesee; #25-509) and 1× MEM non-essential amino acids (Genesee; #25-536), under standard conditions (37 °C, 5% CO_2_). 6 to 12 hours after transfection, 300 𝜇L of DMEM was removed from each well and replaced with 200 𝜇L of fresh DMEM media to reduce reagent effects and concentrate final protein concentration. All split-luciferase assays were carried out at least 36 hours after changing the media.

#### A.2.7. HiBit-LgBit NanoLuc complementation reporter assay for binder expression

Binder expression was quantified by luminescence of the Hibit-LgBit complementation system (Nano-Glo HiBiT Lytic Detection System; Promega) measured with plate reader (Tecan Infinite® M Plex, multimode microplate reader; #30190085). 200 𝜇L of media were first collected from each well in 24-well plates and split equally between 2 wells in 96-well plates. During all experiments, media from the first biological replicate well were distributed to the first two technical replicates of split-luciferase assay, and media from the second biological replicate added to the third technical replicate. Each split-luciferase assay reaction was prepared in a total volume of 100 𝜇L, consisting of 49 𝜇L Nano-Glo® buffer, 44.9 𝜇L phosphate-buffered saline (PBS, pH 7.4), 1 𝜇L Nano-Glo® substrate, 0.1 𝜇L HiBit (20 nM) and 5 𝜇L conditioned media. First luminescence reading was recorded immediately after addition of samples and second measurement was performed after 4 minutes of incubation. Longer incubation time leads to signal saturation, causing the plate reader to end the assay.

To validate that the luminescence signal provides a reliable and quantitative measurement, we leveraged a known PD-L1 binder, KN035 (Zhang et al., 2017) (PDB: 5JDS, 𝐾_𝐷_ = 3 nM), at varying concentrations. We measured luminescence directly from the supernatants after addition of HiBiT (expression) and confirmed a strong linear relationship between luminescence and KN035 concentration upon addition of 20 nM of HiBiT to quantify expression (**Figure S11B** *top*). Encouragingly, measured expression levels from the HiBiT assay (**Figure S11C** *left*) correlated well with protein yields obtained from independent recombinant expression of nanobodies in Expi293F cells (**Figure S29**).

#### A.2.8. SmBit-LgBit NanoLuc complementation reporter assay

Binder-target interaction and specificity was quantified by luminescence of the SmBiT–LgBiT complementation system (Nano-Glo Live Cell Assay System, Promega) on the plate reader (Tecan Infinite® M Plex, multimode microplate reader; #30190085). Media from the same well used in the Hibit-LgBit assay were used to maintain consistency. Each reaction was prepared in a total volume of 100 𝜇L, consisting of 49 𝜇L Nano-Glo® buffer, 39 𝜇L phosphate-buffered saline (PBS, pH 7.4), 1 𝜇L Nano-Glo substrate, 20 nM target protein (from 2 𝜇M stock solution), and 10 𝜇L conditioned media. To assess specificity, parallel reactions were assembled with the same volume of conditioned media but substituting the target protein with 20 nM off-target protein (IL2 for PD-L1 binders; PD-L1 for BHRF1, IL20, and IL3 binders; all from 2 𝜇M stock solutions). First luminescence was recorded right after all the binders were added and second measurement was performed after 8 minutes of incubation.

To verify that binding activity could be represented by SmBiT–LgBiT signals normalized by HiBiT–LgBiT (expression) signals, purified PD-L1 KN035 binder at different concentrations (0.01 nM, 0.1 nM, 1 nM, 10 nM, 100 nM) was tested using the same experimental setup as the split-luciferase assays used for all binders with the exception of the 5 𝜇L of media, which were replaced by direct addition of purified protein at the specified dilutions. HiBiT–LgBiT measurements were not collected at 100 nM due to signal saturation. Both the HiBiT–LgBiT and SmBiT–LgBiT assays exhibited log-linear responses across the tested concentration range (**Figure S11B**), supporting the use of SmBiT–LgBiT normalized by HiBiT–LgBiT as a valid measure of binding activity.

#### A.2.9. Identification of nanobody binder leads for further characterization

We selected candidates for further characterization if they met two criteria: (i) expression luminescence was above 150,000 RLU (**Figure S11C** *left*) and (ii) we observed an increase in normalized binding luminescence between two time points (**Figure S11C** *right*, **S11D**, and **S30; Methods**).

For each split-luciferase assay condition, three technical replicates were averaged to obtain a representative luminescence value for both expression (HiBiT–LgBiT assay) and binding (SmBiT–LgBiT assay). In the initial screens of PD-L1 and IL3, binders were first defined based on binding activity, using a threshold of at least a five-fold increase in normalized binding signal relative to the negative control. A set of PD-L1 nanobodies with different expected activities was also tested on BLI to gain insights into both their expression and binding behaviors. Based on these initial tests, only samples with HiBiT–LgBiT luminescence values above 150,000 arbitrary luminescence units (ALU) were considered expressed. Binding activity for downstream validation was subsequently determined using more refined criteria following our initial PD-L1 experiments: we defined a binder as either (i) an increase in the binding ratio (on-target/off-target) between T1 and T2 greater than 0.5, or (ii) an increase in the on-target binding ratio (on-target T2 / on-target T1) greater than 1.05.

#### A.2.10. Bio-layer interferometry (BLI)

Target proteins with AviTag and SmBiT removed and binder proteins with LgBiT removed, were used for BLI analysis. All reactions were run on an Octet RED96 at 30 ^◦^C, and samples were run in PBS pH 7.4 supplemented with 0.2% BSA and 0.05% Tween 20 (assay buffer). For each of the binder proteins, a titration of five concentrations was prepared by 2-fold serial dilution from the highest concentration tested. Lead nanobody designs were assayed for binding to biotinylated target proteins immobilized onto Octet Streptavidin Biosensors (Sartorius; 18-5019). Sensors were pre-incubated in an assay buffer for 10 minutes, before incubation in wells containing the assay buffer to determine the baseline signal. Sensors were then immersed in wells containing biotinylated target protein (200 nM in assay buffer) for 300 s or until a 1 nm (for PD-L1, IL3, and IL20) or 0.6 nm (BHRF1) BLI shift was reached. Loaded sensors were again washed and baselined in wells containing assay buffer alone, before immersion in nanobody protein solutions for association. Dissociation was monitored by transferring the sensors back into assay buffer alone. For each binder titration, a reference channel was prepared by loading sensors with target protein, but associating in assay buffer alone, to account for any change in signal due to dissociation of target protein from the biosensor tips. Association and dissociation binding curves were fit in Octet System Data Analysis Software version 9.0.0.15 using a global fit 1:1 monovalent model to determine apparent 𝐾_𝐷_s, with signals baseline-corrected by subtraction of the reference channel. Binding curves for monoclonal IgG control antibodies against IL3 and IL20 were fit to 1:2 bivalent binding models. BLI hits were identified if the design displayed clear association kinetics (BLI shift > 0.07 nm for the highest tested concentration). 𝑅^2^ values for each derived apparent 𝐾_𝐷_ are found in **Data S3**.

#### A.2.11. Surface Plasmon Resonance (SPR)

Affinity quantification of designed scFvs by SPR were carried out by the contract research organization Adaptyv Bio. SPR experiments were conducted using a Carterra LSA XT instrument, and at least two independent binding experiments (n = 2) were performed per design. Briefly, designed scFvs were reformatted into fragment antibody (Fab) format, by fusing the designed variable heavy (VH) domain to human IgG1 CH1 (Uniprot P01857, positions 1-103), and fusing the designed variable light (VL) domain to human Ig 𝜅 light chain (Uniprot P01834, positions 1-107). Fab designs were expressed with C-terminal Twin-Strep tags in a prokaryotic in vitro translation system and normalized post-expression using an affinity-based quantification assay. SPR sensor chip surfaces were functionalized by covalently attaching Strep-Tactin XT to a carboxymethylated surface using EDC/NHS coupling, before twin-Strep-tagged Fabs were printed onto the chip using 96-channel bidirectional flow (750 s capture, 600 s equilibration). A PD-L1 dilution series (10–1000 nM, half-log dilutions) was prepared using running buffer (10 mM HEPES, 150 mM NaCl, 3 mM EDTA, 0.05% Tween-20, pH 7.4) and injected onto the chip following a single-cycle kinetics format. Each cycle included 60 s baseline, 300 s association, and 600 s dissociation phases. Surfaces were regenerated between cycles with glycine-HCl, pH 1.5. 𝑅^2^ values were not provided for SPR kinetic fits.

Sensorgrams were processed in Adaptyv Fitting software with baseline and reference subtraction. Data were globally fit to a 1:1 Langmuir model to extract 𝑘_on_, 𝑘_off_, and 𝐾_𝐷_ values. Fabs were classed as hits if they exhibited clearly measurable interaction signal with successful model fits and a calculated 𝐾_𝐷_ ≤ 10 𝜇M. Binders producing association signals >300% above negative controls but lacking reliable fits were manually classified based on response magnitude. The raw data for SPR experiments and corresponding kinetic fits for each replicate for all validated binders can be found in Data S3.

#### A.2.12. Epitope-targeted alanine mutagenesis

Designed epitopes were interrogated by screening binding activity against alanine substitution variants at selected hotspot residues (PD-L1: I37A, Y39A, E41A; IL3: L26A, T103A, F104A; IL20: R59A, R63A, R99A; BHRF1: V85A, I89A, R92A). Residue numbering for all mutants corresponds to the mature protein sequence (i.e., after signal peptide cleavage) and thus differs from full-length UniProt numbering by the length of the signal peptide: +17 for PD-L1, +28 for IL3, +25 for IL20, and +1 for BHRF1. Variant antigens were expressed, purified, and assayed for binding against designed binders by BLI as described for their wild-type counterparts. All BLI data were collected and analyzed using Octet System Data Analysis Software (v9.0.0.15), with signals baseline-corrected by reference channel subtraction.

#### A.2.13. Polyspecificity reagent preparation

Polyspecificity reagent (PSR) was prepared from Expi293F cells as described previously (Makowski et al., 2021; Harvey et al., 2022), with minor modifications. Briefly, 10^9^ cells were pelleted at 550×𝑔 for 3 min, washed once in ice-cold PBS supplemented with 1 mg/mL BSA (Sigma-Aldrich, A4737; PBS-B), and resuspended in ice-cold Buffer B (50 mM HEPES, 150 mM NaCl, 2 mM CaCl_2_, 5 mM KCl, 5 mM MgCl_2_, 10% v/v glycerol, pH 7.2) supplemented with protease inhibitor cocktail. Cells were homogenized (3 × 30 s) and sonicated (3 × 30 s) on ice. The lysate was centrifuged at 40,000 × 𝑔 for 1 h at 4 °C; the supernatant (soluble cytosolic protein fraction) was retained, and the membrane-enriched pellet was resuspended in Buffer B with protease inhibitor using a Dounce homogenizer (30 strokes). Protein concentration was determined using the DC Protein Assay Kit I (Bio-Rad, 5000111). The membrane fraction was diluted to approximately 1 mg/mL in solubilization buffer (50 mM HEPES, 150 mM NaCl, 2 mM CaCl_2_, 5 mM KCl, 5 mM MgCl_2_, 1% n-dodecyl-β-D-maltopyranoside) with protease inhibitor and rotated end-over-end overnight at 4 °C. The suspension was centrifuged at 40,000 × 𝑔 for 1 h at 4 °C to yield the soluble membrane protein (SMP) fraction in the supernatant. Protein concentration was re-determined and adjusted to 0.7 mg/mL. Biotinylation was performed by addition of 20 μL of freshly prepared 10 mM EZ-Link Sulfo-NHS-SS-Biotin (Thermo Fisher Scientific, 21331) per mg SMP, with incubation at 25 °C for 45 min with gentle agitation. Reactions were quenched by addition of 50 μL of 25 mM Tris-HCl pH 8.0 per mL of reaction. Finally, biotinylated PSR was dialyzed into PBS (pH 7.4) overnight at 4 °C using 6–8 kDa MWCO tubing, aliquoted, and stored at −80 °C.

#### A.2.14. Polyspecificity assay

Non-specific binding of purified His- and FLAG-tagged hit binders was assessed using a flow cytometry-based polyspecificity particle assay adapted from Makowski *et al*. (Makowski et al., 2021) and applied as described previously (Hie et al., 2024). Magnetic beads coated with an NTA-Cobal-based ligand (His-Tag Isolation and Pulldown Dynabeads; Thermo Fisher Scientific, 10103D) were washed twice in flow buffer (PBS, pH 7.4 supplemented with 1 mg/mL BSA, 4 °C) and diluted to 540 μg/mL. His- and FLAG-tagged nanobodies or Fabs were diluted to 5 μg/mL or 20 μg/mL, respectively, in flow buffer. Beads (30 μL) were incubated with 85 μL of binder solution in a 96-well plate overnight at 4 °C with rocking. Bead-bound binders were washed twice with flow buffer using a magnetic plate stand, then incubated with 50 μL of 0.1 mg/mL biotinylated PSR for 20–30 min at 4 °C with rocking, followed by one wash in flow buffer. Beads were subsequently incubated for 15 min at 4 °C with streptavidin-APC (BioLegend, 405207; 1:1000), anti-FLAG M2 mouse monoclonal antibody (Sigma-Aldrich, F1804; 1:1000), and goat anti-mouse secondary antibody conjugated to Alexa Fluor Plus 488 (Thermo Fisher Scientific, PIA32723TR; 1:1000). All detection antibody dilutions were prepared in flow buffer. Beads were washed once more, resuspended in 200 μL flow buffer, and analyzed on an Attune NxT flow cytometer (Invitrogen). APC fluorescence reported PSR binding as a measure of polyspecificity, and Alexa Fluor 488 fluorescence confirmed binder loading on beads. Flow cytometry data were analyzed in FlowJo v10.10.1 (BD Biosciences)

#### A.2.15. Expression and purification of anti-Fab nanobody

The anti-Fab nanobody was expressed and purified as previously described (Bloch et al., 2021). Briefly, sequences encoding the anti-Fab nanobody were cloned into a pET26b+ vector containing an N-terminal polyhistidine tag and transformed into BL21(DE3) *E. coli*. Cultures were grown overnight at 37 °C with shaking at 250 rpm in Luria–Bertani (LB) medium supplemented with 50 μg/mL kanamycin, then used to inoculate large-scale cultures grown under the same conditions to OD_600_ 0.6–0.8, at which point expression was induced with 1 mM IPTG for 3–4 h. Cells were harvested by centrifugation at 4,000 × 𝑔 for 15 min and pellets stored at −80 °C or immediately used for purification.

For purification, cell pellets were resuspended in purification buffer (20 mM HEPES, pH 7.4, 150 mM NaCl, 20 mM imidazole, protease inhibitor cocktail) and lysed by sonication. Lysates were clarified by centrifugation at 40,000 × 𝑔 for 45 min and incubated with HisPur Ni-NTA resin (Thermo Fisher Scientific) pre-equilibrated with purification buffer for 1 h at 4 °C. Resin was loaded onto a gravity-flow polypropylene column, washed extensively with purification buffer, and bound protein was eluted with 300 mM imidazole. The eluate was concentrated and further purified by size-exclusion chromatography using a Superdex 200 Increase 10/300 column (Cytiva) pre-equilibrated with imidazole-free purification buffer. Peak fractions corresponding to the anti-Fab nanobody were collected, concentrated, and flash-frozen in liquid nitrogen for storage at −80 °C.

#### A.2.16. Complex assembly

Purified anti-Fab nanobody and anti-PD-L1 Fab were mixed in a 3:1 molar ratio and incubated on ice for 15 min. The mixture was concentrated and subjected to size-exclusion chromatography as described above. Peak fractions corresponding to the nanobody–Fab complex were collected and concentrated (**Figure S31**). Purified PD-L1 was then mixed with the nanobody–Fab complex in a 3:1 molar ratio, incubated on ice for 15 min, and further purified by size-exclusion chromatography. Peak fractions corresponding to the ternary antigen–binder complex were collected and concentrated to approximately 1.0 mg/mL for cryo-EM studies.

#### A.2.17. Cryo-EM sample preparation and data collection

3 μL of purified complex was applied to glow-discharged holey carbon grids (Quantifoil R1.2/1.3 Au, 300 mesh) and vitrified in liquid ethane using a Vitrobot Mark IV instrument (Thermo Fisher Scientific). Grids were imaged on a 300 kV Titan Krios G2 transmission electron microscope equipped with a Selectris imaging filter and Falcon 4i direct electron detector (Thermo Fisher Scientific). Movies were acquired using EPU software (Thermo Fisher Scientific) at a nominal magnification of 130,000×, corresponding to a physical pixel size of 0.98 Å/px, with a defocus range of 1.0 to 2.5 μm. A total electron dose of 50 e^−^/Å^2^ was fractionated across 40 frames.

#### A.2.18. Cryo-EM data processing

A total of 2,613 movies were collected for the PD-L1 binder complex. All processing was performed in cryoSPARC v4.7.1 (Punjani et al., 2017). Movies were preprocessed using patch motion correction and patch CTF estimation. Particles with diameters of 120–140 Å were initially picked using the blob picker and extracted at 4× binning. Two rounds of 2D classification were performed to remove low-quality particles, followed by *ab initio* 3D reconstruction and heterogeneous refinement, yielding classes consistent with a Fab–nanobody–PD-L1 complex. A randomized particle subset was used to train a Topaz model (Bepler et al., 2019) for additional picking, followed by further 2D classification at 4× binning. Particles in high-quality classes were re-extracted without binning and subjected to iterative heterogeneous and non-uniform refinements. The resulting stack was merged with the blob-picked pool, duplicates removed, and subjected to further heterogeneous and non-uniform refinements, yielding an initial ∼4.1 Å reconstruction with clear Fab, nanobody, and PD-L1 density.

To improve resolution, an additional round of template-based and Topaz picking was performed in parallel. Particles from each method were processed through 2D classification and heterogeneous refinement, then merged with duplicates removed. The final stack of 94,644 particles was subjected to non-uniform refinement, yielding a ~3.9 Å reconstruction of the full nanobody–Fab–PD-L1 complex. Masked local refinement at the Fab–PD-L1 interface yielded a final reconstruction at 3.93 Å. Local resolution was estimated in cryoSPARC, with all resolutions reported based on the gold-standard Fourier shell correlation (FSC) criterion of 0.143. A schematic depicting this process, along with a representative cryo-EM micrograph and selected 2D class averages are shown in (**Figure S32**).

#### A.2.19. Model building, refinement, and structural analysis

The predicted structure of Fab-bound PD-L1 was fitted into experimental cryo-EM density maps using UCSF ChimeraX (Goddard et al., 2018). Fitted structures were manually rebuilt in Coot (Emsley and Cowtan, 2004), refined using Phenix (Liebschner et al., 2019), and validated using MolProbity (Davis et al., 2007). Final maps and models were visualized in UCSF ChimeraX.

## B. Supplementary Discussion

### B.1. Role of AF3 Filtering in RFDiffusion

Bennett *et al*. reported a retrospective analysis demonstrating how AlphaFold3 filtering would have improved experimental success rates in their nanobody design campaigns (Bennett et al., 2025). Because Bennett *et al*. did not release their original data, we reanalyzed their reported ROC curves using two independent methods to estimate the success rate that would be achievable by RFDiffusion with AF3 iPTM filtering. Based on standard statistical assumptions, we estimate that the expected hit rate would be approximately 1.2-1.5%. In contrast, our nanobody design success rates range from 4-12% against both “traditional” targets (PD-L1 and BHRF1), for which multiple other methods have successfully designed *de novo* binders, and “challenging” targets (IL3 and IL20), for which previous *de novo* design campaigns have been unsuccessful (Chai Discovery Team et al., 2025; Nabla Bio and Biswas, 2025). Therefore, we estimate that Germinal’s success rate is several-fold higher than that of Bennett *et al*., even when accounting for potential improvements from AF3 filtering.

We employed two independent approaches to estimate the potential success rate achievable by Bennett *et al*. if AF3 was used for filtering. Our first approach to estimating the AF3-enabled success rate is based on the published ROC curve, where we selected the most optimistic operating point, which corresponds to an approximately 0.38 true positive rate (TPR) at 0.03 false positive rate (FPR). Using the reported binder prevalence (𝑝 ≈ 0.12% given the 22 successful binders out of 18,190 total designs across the four targets in their analysis), we calculated an estimated hit rate (or positive predictive value) of approximately 1.5% as:

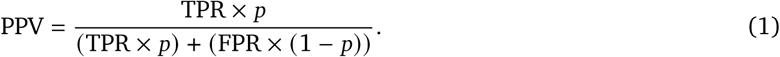

For our second approach to estimating the AF3-enabled success rate, we modeled the AF3 iPTM scores for binders and non-binders as two normal distributions with equal parameters 𝜇 (mean) and 𝜎 (standard deviation) chosen to reproduce the published AUC of 0.86:

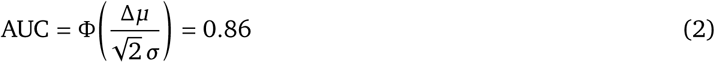

where Φ denotes the cumulative density function (CDF) of the normal distribution. By normalizing the scale to 𝜎 = 1, we obtain

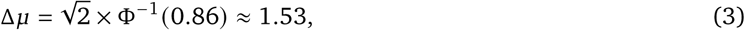

where Φ^−1^ denotes the inverse CDF.

Under this model, we estimated the TPR and FPR at a stringent selection threshold corresponding to the top 4% of designs (iPTM > 0.85 as reported in Bennett *et al*.) as:

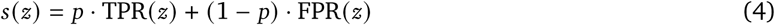

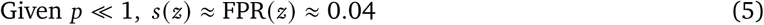

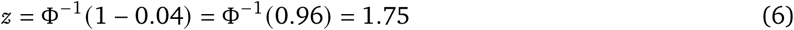

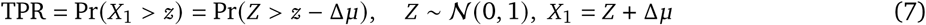

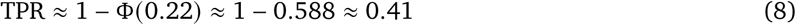

Lastly, we compute the PPV as in (1). This approach yielded an estimated hit rate of approximately 1.2%.

### B.2. Epitope Selection for Known Targets

Design success in *de novo* binder generation is known to vary substantially depending on the target and the specific epitope region. For mini-protein binders, success rates have been shown to range from 9% to 88% across targets, with complete failure on certain proteins such as TNF𝛼 (Zambaldi et al., 2024). Surface properties including high polarity, convex curvature, and glycosylation have been identified as factors that reduce designability (Cao et al., 2022; Yang et al., 2024). More recently, systematic epitope-level benchmarking has revealed that even on the same target, different epitope regions can yield dramatically different success rates, with some epitopes producing no viable designs at all (Zambaldi et al., 2024; Nabla Bio, 2025). In the antibody design setting, JAM-2 reported binders to 30–70% of user-specified epitopes on half of their tested targets, underscoring that not all surface regions are equally amenable to computational antibody design even with state-of-the-art methods.

Motivated by these observations, we adopted a strategy of running preliminary unconstrained campaigns—in which no epitope residues were specified—to identify the most productive epitope for each target prior to full-scale runs. As described earlier, for antigens with known binders (e.g., natural receptors), we designated the epitope corresponding to the reference biologically-relevant interaction as the preferred epitope, both because targeting natural interaction interfaces is often of direct therapeutic interest, and because these regions emerged most frequently among designs passing in silico filters during preliminary runs. However, to quantify how strongly epitope choice influences pipeline productivity, we ran a retrospective comparison across candidate epitopes for IL3 and IL20, two targets with known binding partners. For each target, all epitope conditions used identical pipeline parameters, varying only the hotspot residues. Each condition was run for 48 hours. Differences in the number of started trajectories reflect variation in trajectory length depending on whether a design passes through the full pipeline or terminates early due to low confidence scores.

For IL3, three epitope sets were evaluated (Table B.2.1). The natural receptor binding site (HT1; used in this work) yielded 26 accepted designs from 12 seeds, compared to 6 designs (2 seeds) for an alternative PDB crystal contact interface (HT3) and zero for an uncharacterized surface patch (HT2). Early filter pass rates differed by over an order of magnitude: 36.4% (HT1) vs. 2.4% (HT2) and 1.2% (HT3), indicating that epitope choice dominates pipeline throughput. Average scores for trajectories that pass the hallucination stage and are redesigned with AbMPNN are shown in Table B.2.2.

For IL20, two epitope sets were evaluated (Table B.2.3). The natural receptor binding site (HT2) produced zero designs as a result of no designs passing early filters across 1,804 attempts. Meanwhile, an uncharacterized patch (HT1; used in this work) yielded 9 accepted designs from 6 seeds. Average scores for HT1 trajectories that pass the hallucination stage and were redesigned with AbMPNN are shown in Table B.2.4.

Together, these results suggest that while known binding interfaces provide a reasonable starting point for epitope selection, consistent with these regions representing intrinsically favorable protein–protein interaction surfaces, design productivity can vary substantially across epitope regions and targets, and not all surface regions are equally amenable to computational binder design. We therefore recommend running preliminary design campaigns to identify the most productive epitopes for a given target before committing to a full-scale design run. Importantly, despite engaging similar epitope regions to known binders, the designed nanobodies adopt structurally distinct binding modes, as evidenced by the low Foldseek TM-scores and iAlign IS-scores relative to known complexes (**Figure S7**).

**Table B.2.1.** IL3 epitope screening results. HT1 corresponds to the known natural receptor binding site.

| Condition | Residues | Type | Attempts | Hallucination pass rate | Accepted | Seeds |
| --- | --- | --- | --- | --- | --- | --- |
| HT1 | D12, L26, F103, F104, K105 | Natural receptor site | 1,469 | 36.4% | 26 | 12 |
| HT2 | R46, E50, K70, L73 | Uncharacterized | 3,063 | 2.4% | 0 | 0 |
| HT3 | E13, H17, S67, N71 | Alternative interface | 3,193 | 1.2% | 6 | 2 |

**Table B.2.2.** Average scores for IL3 designs that completed the hallucination stage and before AbMPNN.

| Metric | HT1 ($n = 504$) | HT2 ($n = 65$) | HT3 ($n = 29$) |
| --- | --- | --- | --- |
| iPSAE | $0.229 \pm 0.259$ | $0.090 \pm 0.158$ | $0.393 \pm 0.285$ |
| ipTM | $0.485 \pm 0.244$ | $0.337 \pm 0.206$ | $0.618 \pm 0.252$ |
| pTM | $0.676 \pm 0.119$ | $0.604 \pm 0.094$ | $0.743 \pm 0.123$ |
| pDockQ2 | $0.181 \pm 0.231$ | $0.099 \pm 0.186$ | $0.262 \pm 0.274$ |
| Interface $\Delta G$ | $-49.5 \pm 10.5$ | $-41.2 \pm 9.5$ | $-49.2 \pm 10.8$ |
| Interface hydrophobicity | $58.7 \pm 10.9$ | $59.3 \pm 10.1$ | $54.2 \pm 11.8$ |
| Interface H-bonds | $5.2 \pm 2.7$ | $4.3 \pm 2.0$ | $6.2 \pm 2.9$ |
| Shape complementarity | $0.652 \pm 0.063$ | $0.618 \pm 0.070$ | $0.636 \pm 0.059$ |
| Surface hydrophobicity | $0.419 \pm 0.047$ | $0.413 \pm 0.034$ | $0.434 \pm 0.040$ |
| AbMPNN score | $0.708 \pm 0.103$ | $0.699 \pm 0.089$ | $0.648 \pm 0.120$ |

**Table B.2.3.**
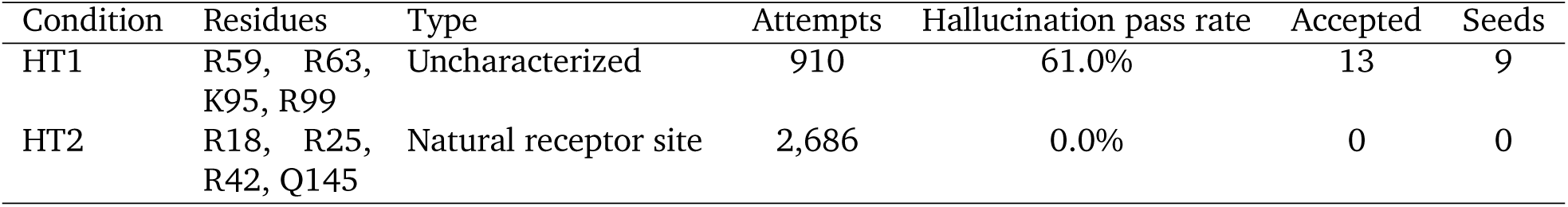
IL20 epitope screening results. HT2 corresponds to the natural receptor binding interface.

**Table B.2.4.** Average scores for IL20 designs that completed the hallucination stage and before AbMPNN redesign (mean ± s.d.). HT2 corresponds to the known natural receptor binding site, yet it produced no designs that passed early filters.

| Metric | HT1 ($n = 539$) |
| --- | --- |
| ipSAE | $0.237 \pm 0.246$ |
| ipTM | $0.790 \pm 0.036$ |
| pTM | $0.855 \pm 0.017$ |
| pDockQ2 | $0.255 \pm 0.270$ |
| Interface $\Delta G$ | $-44.4 \pm 9.5$ |
| Interface hydrophobicity | $58.9 \pm 9.2$ |
| Interface H-bonds | $7.6 \pm 2.7$ |
| Shape complementarity | $0.632 \pm 0.066$ |
| Surface hydrophobicity | $0.427 \pm 0.034$ |
| AbMPNN score | $0.698 \pm 0.114$ |

### B.3. Targeting Proteins Without Known Experimental Structures

MCF2 (MCF.2 cell surface antigen) has no experimentally determined structure in the PDB. We selected it as a biologically relevant test case because it is a proto-oncogene and therefore represents a potentially interesting target for binder-based therapeutic strategies such as targeted degradation. Among proteins lacking experimental structural data, it also provided a practical starting point for structure-guided design, as its predicted DH domain appeared compact and high-confidence, without extensive low-confidence loops, disordered regions, or linker-like segments. We used an AlphaFold3-predicted structure of the DH domain as input and tested two hotspot-guided epitope sets alongside a no-hotspot control (Table B.3.1). All three conditions were run for 48 hours.

**Table B.3.1.** Targeted epitopes for anti-MCF2 antibody design. All epitopes produced viable designs in comparable quantities.

| Condition | Residues | Accepted | Seeds |
| --- | --- | --- | --- |
| HT1 | A22, A26, A79 | 22 | 12 |
| HT2 | A33, A38, A127, A131 | 28 | 12 |
| No hotspot | — | 34 | 12 |

All three conditions produced viable designs across at least 12 unique seeds, demonstrating that Germinal can generate high-confidence binder designs even in the absence of an experimentally determined target structure. Average trajectory-level scores for designs that successfully completed the design stage are shown in Table B.3.2, and scores for designs that passed all pipeline filters are shown in Table B.3.3. Notably, MCF2 designs exhibit structural confidence and Rosetta metrics comparable to those obtained for IL3 and IL20, two targets for which Germinal designs were experimentally validated, suggesting that the computational quality of MCF2 designs is on par with those that yielded functional binders, though experimental validation remains to be performed.

**Table B.3.2.** Average scores for MCF2 redesign candidates (mean ± s.d.).

| Metric | HT1 ($n = 323$) | HT2 ($n = 311$) | No hotspot ($n = 141$) |
| --- | --- | --- | --- |
| iPSAE | $0.320 \pm 0.251$ | $0.379 \pm 0.244$ | $0.380 \pm 0.267$ |
| ipTM | $0.585 \pm 0.206$ | $0.630 \pm 0.195$ | $0.619 \pm 0.219$ |
| pTM | $0.777 \pm 0.085$ | $0.795 \pm 0.082$ | $0.790 \pm 0.092$ |
| pDockQ2 | $0.330 \pm 0.282$ | $0.410 \pm 0.289$ | $0.377 \pm 0.279$ |
| Interface $\Delta G$ | $-43.2 \pm 9.7$ | $-44.3 \pm 10.4$ | $-50.9 \pm 14.0$ |
| Interface hydrophobicity | $47.4 \pm 8.9$ | $44.8 \pm 11.0$ | $49.5 \pm 10.7$ |
| Interface H-bonds | $7.2 \pm 2.7$ | $7.2 \pm 2.8$ | $6.9 \pm 2.9$ |
| Shape complementarity | $0.625 \pm 0.066$ | $0.635 \pm 0.061$ | $0.640 \pm 0.062$ |
| Surface hydrophobicity | $0.414 \pm 0.036$ | $0.404 \pm 0.039$ | $0.423 \pm 0.035$ |
| AbMPNN score | $0.721 \pm 0.118$ | $0.724 \pm 0.117$ | $0.647 \pm 0.142$ |

**Table B.3.3.** Structural and interface scores for accepted MCF2 designs across hotspot conditions (mean ± SD).

| Metric | HT1 ($n = 22$) | HT2 ($n = 28$) | No hotspot ($n = 34$) |
| --- | --- | --- | --- |
| ipSAE | $0.628 \pm 0.095$ | $0.589 \pm 0.057$ | $0.684 \pm 0.071$ |
| ipTM | $0.805 \pm 0.042$ | $0.782 \pm 0.025$ | $0.817 \pm 0.039$ |
| pTM | $0.874 \pm 0.020$ | $0.862 \pm 0.012$ | $0.876 \pm 0.021$ |
| pDockQ2 | $0.715 \pm 0.092$ | $0.706 \pm 0.075$ | $0.639 \pm 0.137$ |
| Interface $\Delta G$ | $-52.3 \pm 10.3$ | $-48.1 \pm 6.4$ | $-54.7 \pm 9.6$ |
| Interface hydrophobicity | $53.6 \pm 8.8$ | $54.6 \pm 7.0$ | $54.8 \pm 5.7$ |
| Interface H-bonds | $8.0 \pm 2.4$ | $8.4 \pm 2.1$ | $8.5 \pm 2.7$ |
| Shape complementarity | $0.667 \pm 0.031$ | $0.663 \pm 0.054$ | $0.626 \pm 0.032$ |
| Surface hydrophobicity | $0.378 \pm 0.023$ | $0.376 \pm 0.014$ | $0.379 \pm 0.014$ |
| AbMPNN score | $0.679 \pm 0.146$ | $0.706 \pm 0.095$ | $0.605 \pm 0.124$ |

## C. Supplementary Tables

**Table S1.** Effect of AbMPNN in design metrics. Parent vs AbMPNN-generated sequences.

| Metric | Parent (mean ± sd) | AbMPNN (mean ± sd) | $\Delta$ |
| --- | --- | --- | --- |
| pLDDT | 0.886 ± 0.029 | 0.890 ± 0.023 | +0.004 |
| ptm | 0.833 ± 0.072 | 0.836 ± 0.058 | +0.003 |
| i_ptm | 0.774 ± 0.135 | 0.773 ± 0.114 | -0.000 |
| i_pae | 0.871 ± 0.141 | 0.873 ± 0.118 | +0.002 |
| pae | 5.811 ± 1.713 | 5.754 ± 1.479 | -0.057 |
| pDockQ | 0.645 ± 0.231 | 0.665 ± 0.168 | +0.020 |
| interface_hbonds | 7.602 ± 2.899 | 7.549 ± 2.189 | -0.054 |
| interface_dG | -51.559 ± 11.856 | -51.155 ± 8.967 | +0.404 |
| interface_dSASA | 1756.989 ± 323.357 | 1733.830 ± 291.392 | -23.159 |
| iglm_ll | -1.866 ± 0.063 | -1.762 ± 0.042 | +0.104 |
| interface_nres | 20.094 ± 4.459 | 19.903 ± 4.167 | -0.190 |
| **binder_score** | **-408.835 ± 14.757** | **-417.079 ± 11.046** | **-8.244** |
| interface_shape_comp | 0.695 ± 0.043 | 0.686 ± 0.040 | -0.010 |

**Table S2.** Set of confidence metrics and PyRosetta-derived scores used as pipeline filters for nanobodies.

| Metric | Condition | Details |
| --- | --- | --- |
| Binder near the epitope | True | Intersection of predicted interface residues and epitope residues. Should be at least 3 residues. |
| CDR3-epitope contacts | > 1 | Number of CDR3 residues that are part of the interface residues. |
| Clashes after relaxation | = 0 | Rosetta metric. |
| Surface hydrophobicity | ≤ 0.4 | Rosetta metric. |
| Interface hydrophobicity | ≥ 0.45 | Rosetta metric. |
| pDockQ2 | > 0.23 | Calculated according to the author's implementation (Zhu et al., 2023). |
| Interface shape complementarity | ≥ 0.6 | Rosetta metric. |
| Binder RMSD | < 6 | RMSD between the final predicted structure (AF3) and the initial design output by the pipeline (AF-M). |
| Interface hydrogen bonds | ≥ 3 | Rosetta metric. |
| CDR Interface Percentage | > 50% | Number of residues at the interface that are CDR residues over total interface residues. |
| pLDDT | ≥ 80 | AF3 metric. |
| pTM | ≥ 0.80 | AF3 metric. |
| ipTM | ≥ 0.75 | AF3 metric. |
| pAE | < 8 | AF3 metric. |

**Table S3.**
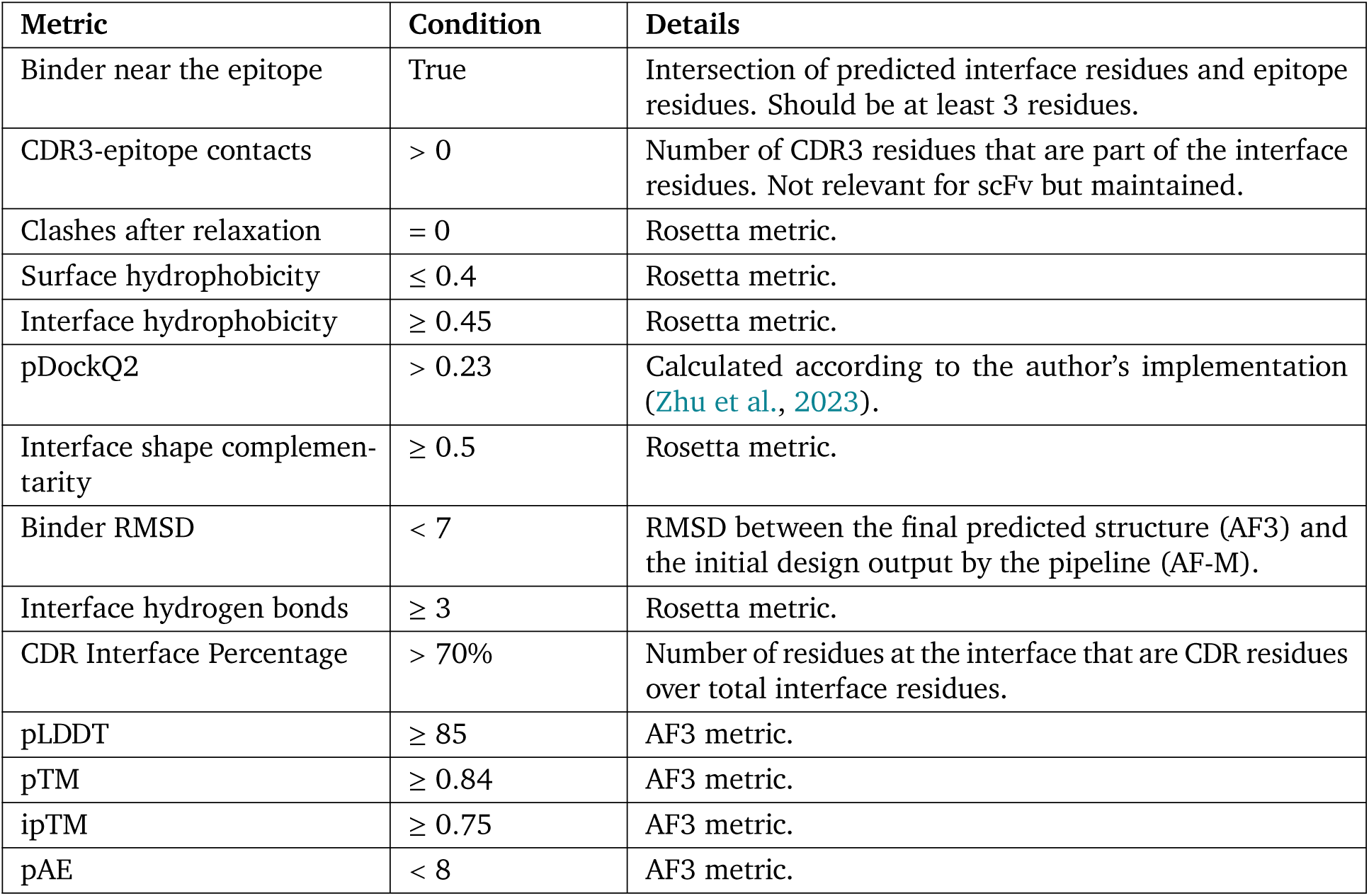
Set of confidence metrics and PyRosetta-derived scores used as pipeline filters for scFvs.

**Table S4.** Set of metrics used to filter and rank nanobody designs for testing.

| Metric | Condition | Ranking weight | Notes |
| --- | --- | --- | --- |
| IgLM log-likelihood | $\geq -1.85$ | 0.10 | IgLM's confidence in the sequence. |
| pAE | $\leq 6$ | | AF3 metric. |
| pTM | $\geq 0.83$ | | AF3 metric. |
| ipTM | $\geq 0.75$ | | AF3 metric. |
| Binder score | $\leq -390$ | 0.10 | Rosetta energy score of the binder. |
| Interface shape complementarity | $\geq 0.6$ | 0.10 | Rosetta metric. |
| Interface hydrophobicity | $\geq 50\%, \leq 60\%$ | 0.10 | Rosetta metric. |
| Interface hydrogen bonds | $\geq 4$ | 0.15 | Rosetta metric. |
| # of framework mutations | $< 3$ | | |
| CDR-specific SAP score | $< 36$ | 0.10 | Rosetta metric. |
| pDockQ2 | $\geq 0.45$ | 0.10 | |
| CDR3-epitope contacts | $\geq 4$ | 0.10 | |
| CDR Interface Percentage | $\geq 0.5$ | | |
| Interface dG | $< -35$ | | Rosetta energy score of the interface. |
| AbMPNN log-likelihood |  | 0.10 | AbMPNN's confidence in its designed residues. |

**Table S5.** Set of metrics used to filter and rank scFv designs for testing.

| Metric | Condition | Ranking weight | Notes |
| --- | --- | --- | --- |
| IgLM log-likelihood |  | 0.15 | IgLM's confidence in the sequence. |
| pAE | < 6.75 |  | AF3 metric. |
| pTM | $\geq 0.85$ | | AF3 metric. |
| ipTM | $\geq 0.75$ | | AF3 metric. |
| pLDDT | $\geq 0.87$ | | AF3 metric. |
| Binder score |  | 0.15 | Rosetta energy score of the binder. |
| Interface shape complementarity | > 0.4 |  | Rosetta metric. |
| Interface hydrophobicity | $\geq 42\%, \leq 59\%$ | 0.10 | Rosetta metric. |
| Interface hydrogen bonds | > 9 | 0.15 | Rosetta metric. |
| Interface number of residues | $\geq 20$ | 0.15 | Rosetta metric. |
| # of framework mutations | < 2 |  | Number of framework residues that are different from the original. |
| CDR-specific SAP score |  | 0.10 | Rosetta metric. |
| pDockQ2 | $\geq 0.33$ | 0.10 | Calculated according to the author's implementation (Zhu et al., 2023). |
| CDR Interface Percentage | > 0.7 |  | Number of residues at the interface that are CDR residues over total interface residues. |
| Interface dG |  | 0.15 | Rosetta energy score of the interface. |
| Interface dSASA |  | 0.15 | Rosetta energy score of the interface. |

**Table S6.** Structural novelty of hit designs. Maximum IS-score across all Foldseek PDB100 matches (deposited before 2021-09-30).

| Campaign | Design | IS<sub>max</sub> | PDB |
| --- | --- | --- | --- |
| PD-L1 Nanobody | G9 | 0.236 | 5XXY |
|  | H9 | 0.257 | 5XXY |
|  | E11 | 0.192 | 6ARQ |
|  | F11 | 0.200 | 7V24 |
|  | E4 | 0.191 | 6AOD |
|  | D10 | 0.247 | 6X4T |
|  | F10 | 0.248 | 5XXY |
| PD-L1 scFv | E5 | 0.184 | 4FFY |
|  | G5 | 0.176 | 1XIW |
|  | H5 | 0.186 | 1P4I |
| IL3 Nanobody | E1 | 0.189 | 5K9K |
|  | D2 | 0.173 | 5UV8 |
| IL3 scFv | *F4<sub>Parent</sub>* | 0.200 | 5UWC |
|  | *F4<sub>Sister</sub>* | 0.197 | 5UWC |
|  | F4 | 0.200 | 5UWC |
| BHRF1 Nanobody | D3 | 0.364 | 3PK1 |
|  | A5 | 0.198 | 6GWQ |
|  | C4 | 0.293 | 6YLI |
|  | E12 | 0.253 | 6WH0 |
|  | A3 | 0.224 | 3PGF |
|  | C12 | 0.466 | 7P33 |
| IL20 Nanobody | B2 | 0.219 | 3KYI |
|  | H5 | 0.203 | 5M4Y |
|  | G5 | 0.214 | 5M4Y |
|  | D3 | 0.194 | 5M4Y |

**Table S7.** Pairwise structural diversity (iAlign IS-score) of designed antibodies. Higher values indicate greater structural similarity between designs. All three IL3 scFv share the same seed ans structure and thus have IS-scores > 0.75.

| Campaign | Subset | N | Pairs | Median | Mean |
| --- | --- | --- | --- | --- | --- |
| PD-L1 Nanobody | All | 101 | 5050 | 0.224 | 0.248 |
|  | Hits | 7 | 21 | 0.255 | 0.375 |
| PD-L1 scFv | All | 48 | 1128 | 0.242 | 0.270 |
|  | Hits | 3 | 3 | 0.228 | 0.430 |
| IL3 Nanobody | All | 46 | 1035 | 0.198 | 0.220 |
|  | Hits | 2 | 1 | 0.146 | 0.146 |
| IL3 scFv | All | 50 | 1275 | 0.254 | 0.276 |
|  | Hits | 3 | 3 | 0.770 | 0.768 |
| BHRF1 Nanobody | All | 52 | 1275 | 0.221 | 0.241 |
|  | Hits | 6 | 15 | 0.225 | 0.233 |
| IL20 Nanobody | All | 43 | 861 | 0.197 | 0.229 |
|  | Hits | 4 | 6 | 0.445 | 0.514 |

**Table S8.** Pairwise CDR sequence identity of designed antibodies. Higher values indicate greater sequence similarity between designs.

| Campaign | Subset | *N* | Pairs | Median | Mean |
| --- | --- | --- | --- | --- | --- |
| PD-L1 Nanobody | All | 101 | 5050 | 0.108 | 0.112 |
|  | Hits | 7 | 21 | 0.162 | 0.255 |
| PD-L1 scFv | All | 48 | 1128 | 0.200 | 0.202 |
|  | Hits | 3 | 3 | 0.240 | 0.473 |
| IL3 Nanobody | All | 46 | 1035 | 0.081 | 0.104 |
|  | Hits | 2 | 1 | 0.135 | 0.135 |
| IL3 scFv | All | 50 | 1275 | 0.180 | 0.180 |
|  | Hits | 3 | 3 | 0.560 | 0.693 |
| BHRF1 Nanobody | All | 52 | 1275 | 0.081 | 0.099 |
|  | Hits | 6 | 15 | 0.081 | 0.101 |
| IL20 Nanobody | All | 43 | 861 | 0.108 | 0.117 |
|  | Hits | 4 | 6 | 0.203 | 0.311 |

**Table S9.**
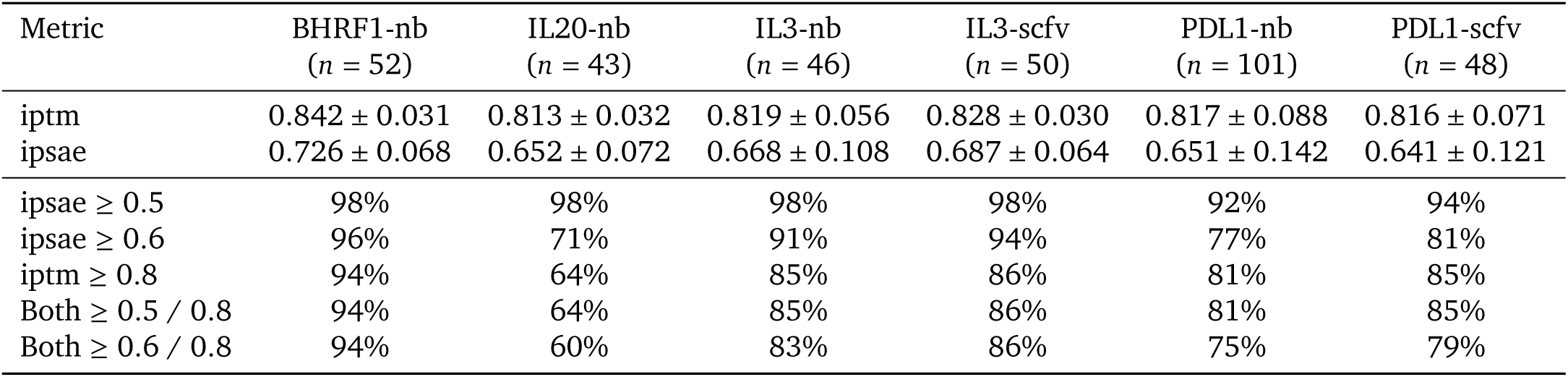
ipTM and min_ipSAE scores for Germinal designs across targets (mean ± SD), with passing rates at two ipSAE thresholds.

## D. Supplementary Figures

**Figure S1.**
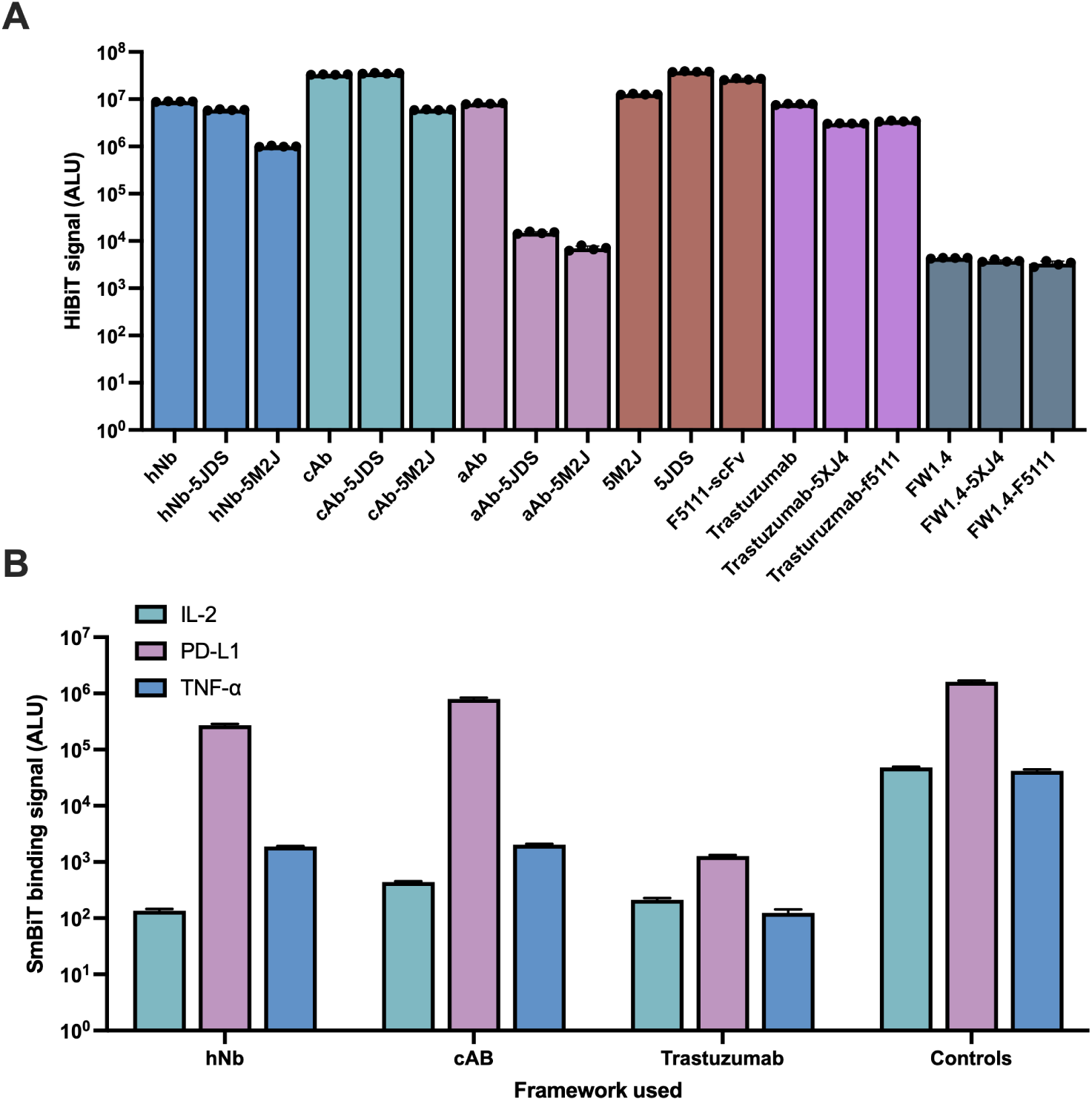
Experimental validation of different nanobody and scFv frameworks. (**A,B**) Different combinations of engrafted CDRs were tested on different antibody frameworks to select the one with highest expression (**A**) and capacity to withstand changes to their CDRs while maintaining binding (**B**). Three nanobody frameworks: hNbBCII (PDB: 3EAK), cAbBCII (PDB: 3DWT) and aAb), and two scFv frameworks: trastuzumab (PDB: 6ZQK) and FW1.4 (Borras et al., 2010) were first tested for expression. For nanobodies, CDRs from a PD-L1 (PDB: 5JDS) and a TNF-𝛼 (PDB: 5M2J) nanobody were used. Similarly, for scFvs, CDRs from F5111 (Trotta et al., 2018), an IL2 Fab, and a PD-L1 scFv (PDB: 5XJ4) were used. Designs were only tested if they showed substantial expression levels (> 10^6^). Bar plots in (**B**) show each of the two engrafted design per framework against its intended antigen, plus a negative control (i.e., hNb-5JDS exclusively against PD-L1, hNb-5M2J exclusively against TNF-𝛼, and a negative control wild-type hNb against an unrelated antigen, IL2). For nanobodies, IL2 was used as control while for scFv TNF-𝛼 was used as negative control.

**Figure S2.**
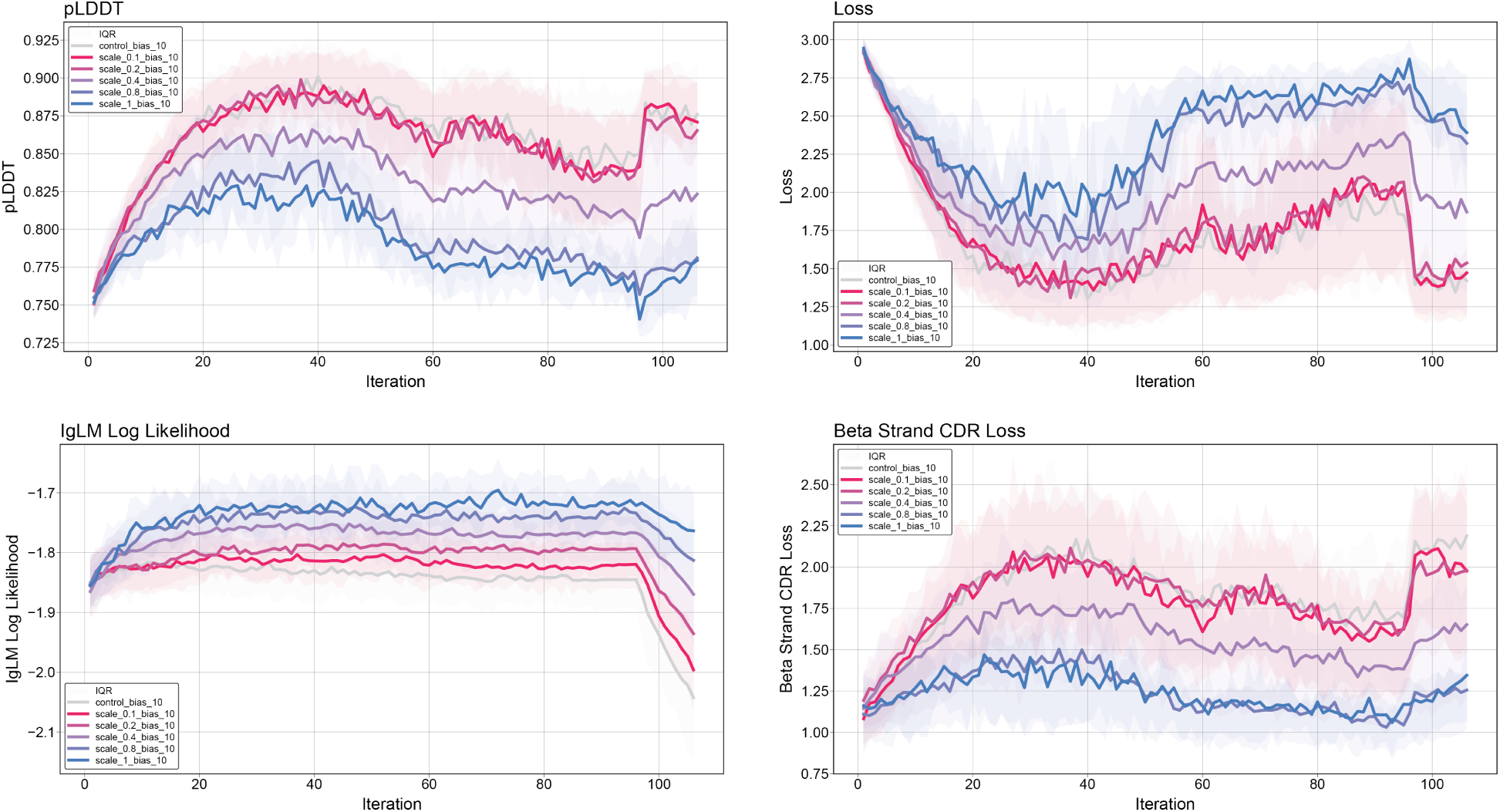
Effect of IgLM guidance over entire design trajectories. The effect of incorporating IgLM-based guidance during the design trajectories across 110 iterations is shown, evaluated under different bias scaling factors (0.1, 0.2, 0.4, 0.8, 1.0) compared to a control without IgLM guidance.

**Figure S3.**
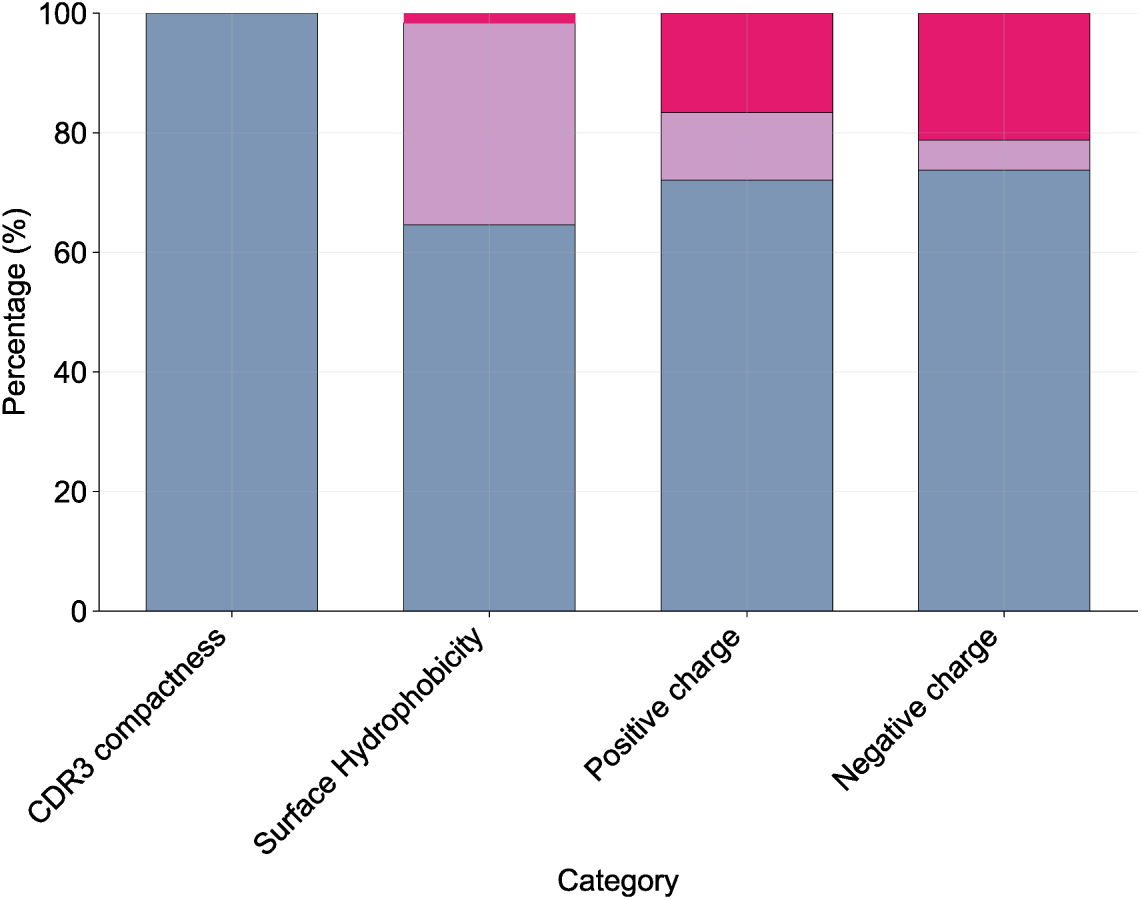
Therapeutic Nanobody Profiler metrics for nanobody hits. Stacked bar chart showing the percentage distribution of low (blue), medium (purple), and high (pink) risk across all BLI-verified nanobody across all target proteins (BHRF1, IL20, IL3, and PD-L1). Designs which failed to be evaluated by TNP were excluded. The CDR length metrics are omitted due to the use of a fixed framework for which CDR length does not change among designs.

**Figure S4.**
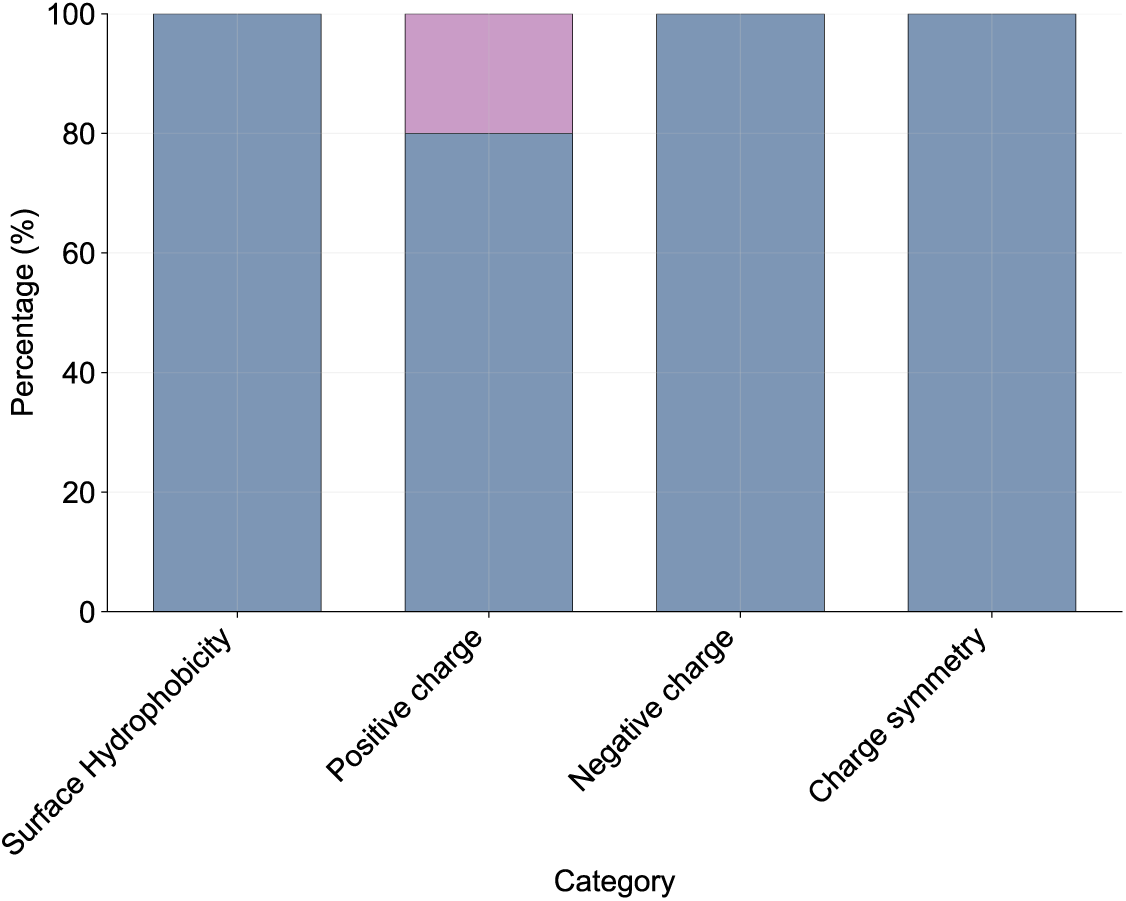
Therapeutic Antibody Profiler metrics for scFv candidates. Stacked bar chart showing the percentage distribution of low (blue), medium (purple), and high (pink) risk across a set of PD-L1 scFv candidates selected for experimental validation. CDR length metrics are omitted because a fixed framework was used, so CDR lengths do not vary across designs.

**Figure S5.**
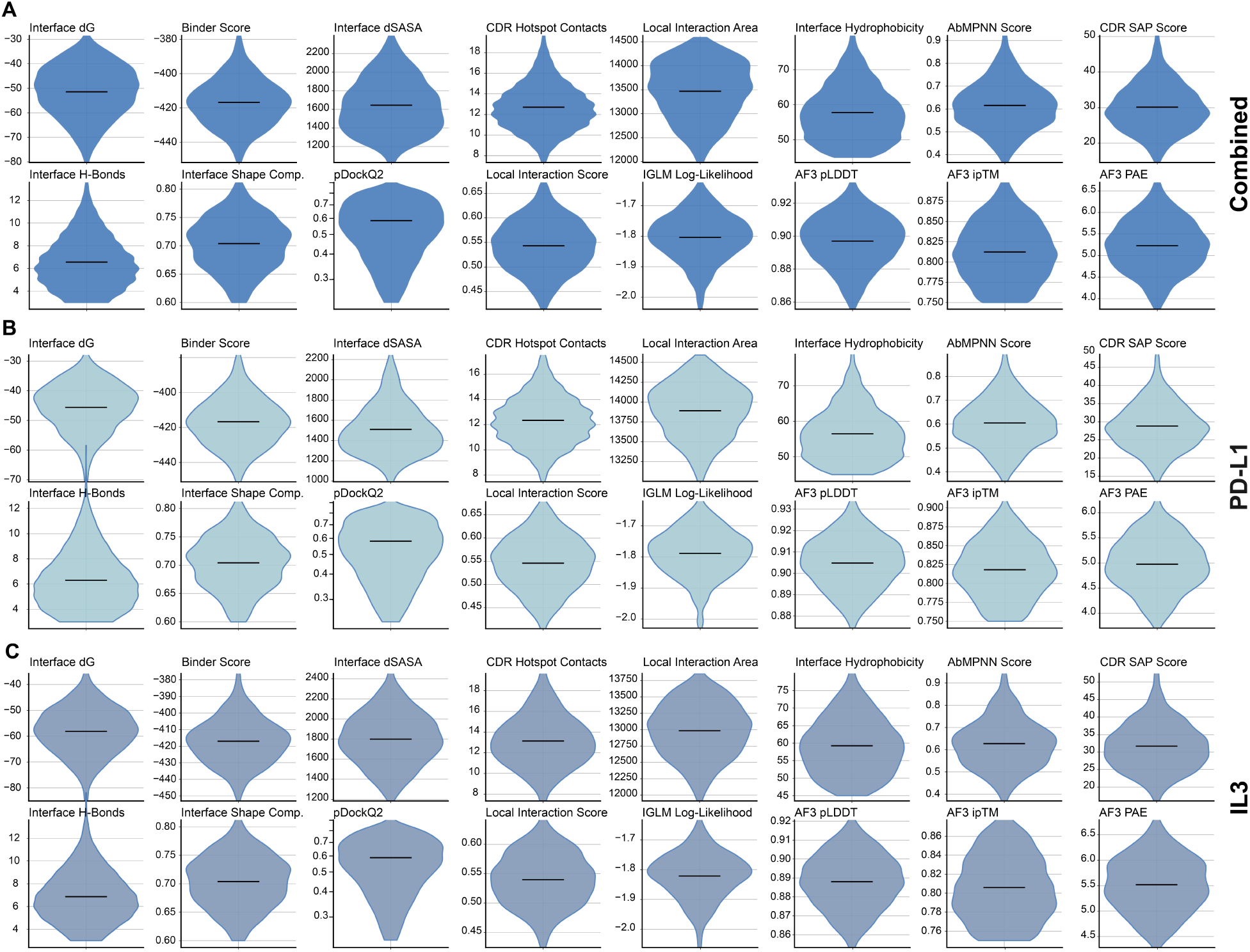
Distribution of key computational metrics for all sampled nanobody designs for PD-L1 and IL3. (**A**) Distributions across all successful designs pooled from different targets. (**B**, **C**) Specific distributions for PD-L1 (**B**) and IL3 designs (**C**).

**Figure S6.**
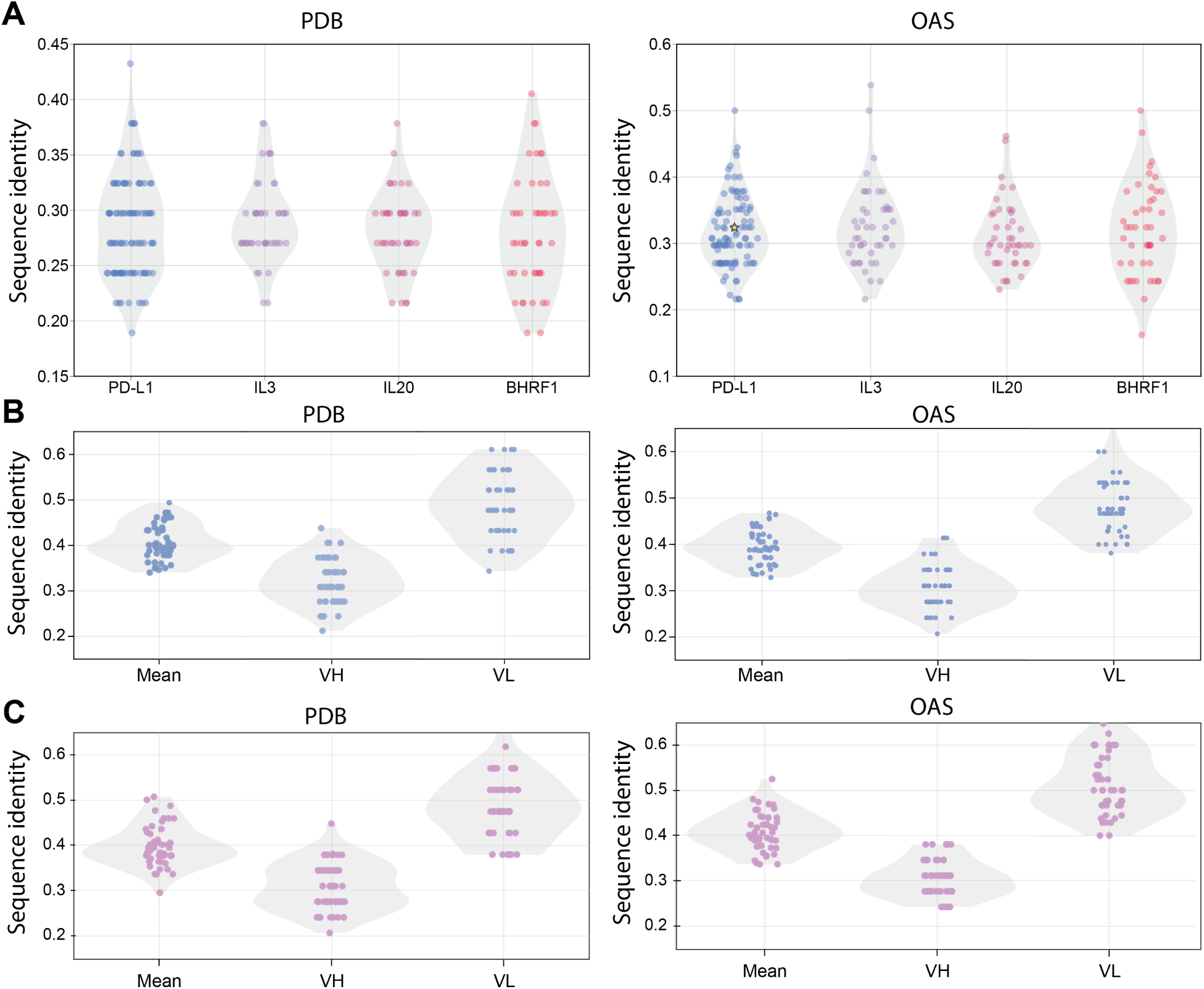
CDR sequence similarities to antibodies in PDB and OAS. Alignments were created against the entire query sequence using mmseqs with a sensitivity of 3.0. The top hits were extracted and CDR similarity was calculated for each. Sequence identity is reported for the CDR positions only. (**A**) Sequence identity for experimentally validated nanobody designs for all four targets against PDB and OAS. The yellow star indicates the sequence similarity of the positive control PD-L1 binder, KN035, against OAS. Sequence identity for experimentally validated PD-L1 (**B**) and IL3 (**C**) scFv designs against PDB and OAS. Each chain was calculated separately.

**Figure S7.**
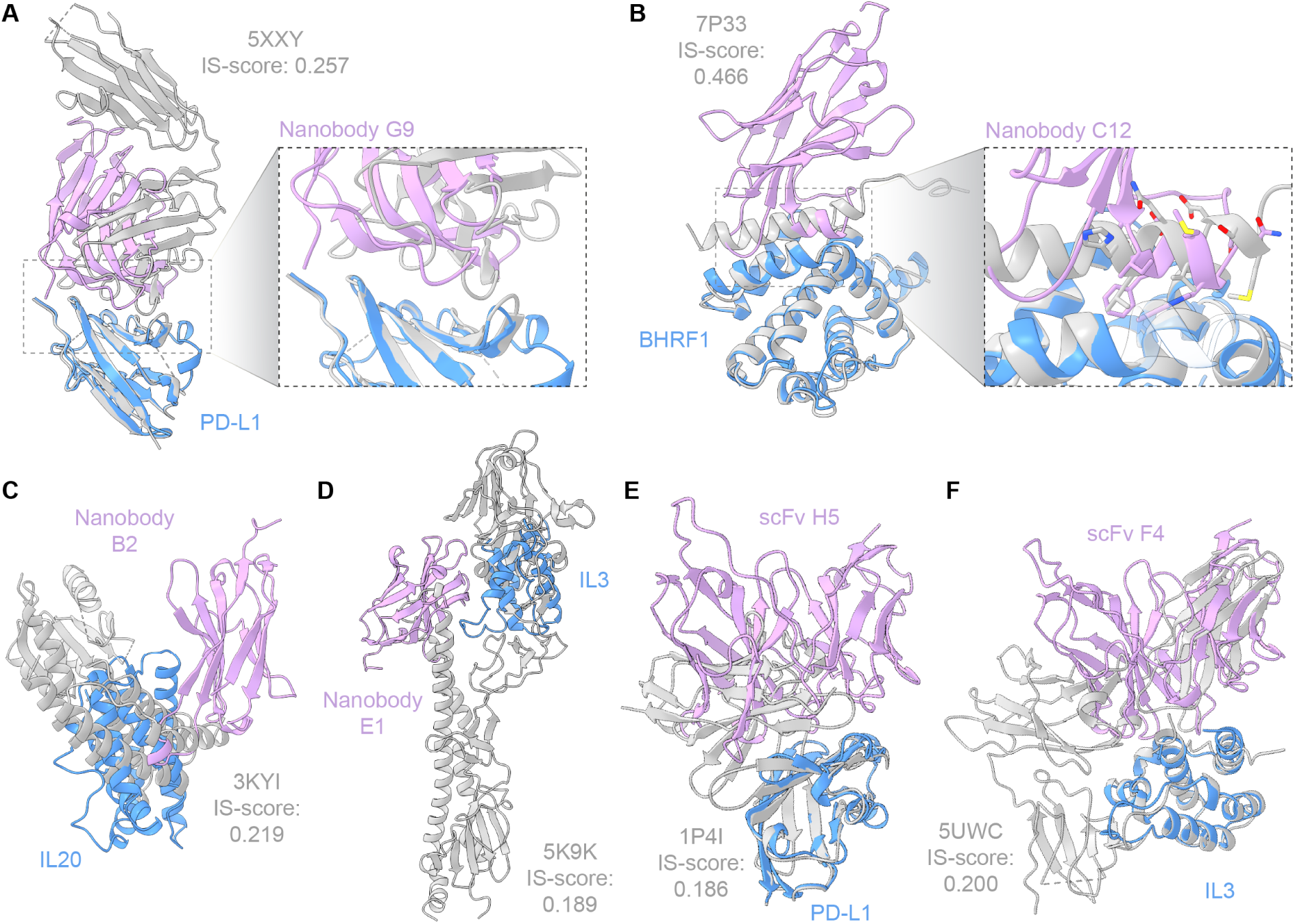
Structural similarity analysis of designed binder–target interfaces against known protein structures. (**A–F**) For each designed binder, the predicted binder–target interface was queried against the Protein Data Bank using iAlign. The top match by Interface Similarity (IS) score is shown (grey cartoon; PDB ID and IS-score indicated) following interface-based structural alignment to the Germinal-predicted binder (pink cartoon)–target (blue cartoon) complex, with a zoom panel showing the aligned interface. PD-L1 nanobody G9 vs. PDB 5XXY (IS-score: 0.257) (**A**); BHRF1 nanobody C12 vs. PDB 7P33 (IS-score: 0.466) (**B**); IL20 nanobody B2 vs. PDB 3KYI (IS-score: 0.219) (**C**); IL3 nanobody E1 vs. PDB 5K9K (IS-score: 0.189) (**D**); PD-L1 scFv H5 vs. PDB 1P4I (IS-score: 0.186) (**E**); IL3 scFv F4 vs. PDB 5UWC (IS-score: 0.200) (**F**). PDB structures deposited after the AlphaFold-Multimer training cutoff date were excluded from the search.

**Figure S8.**
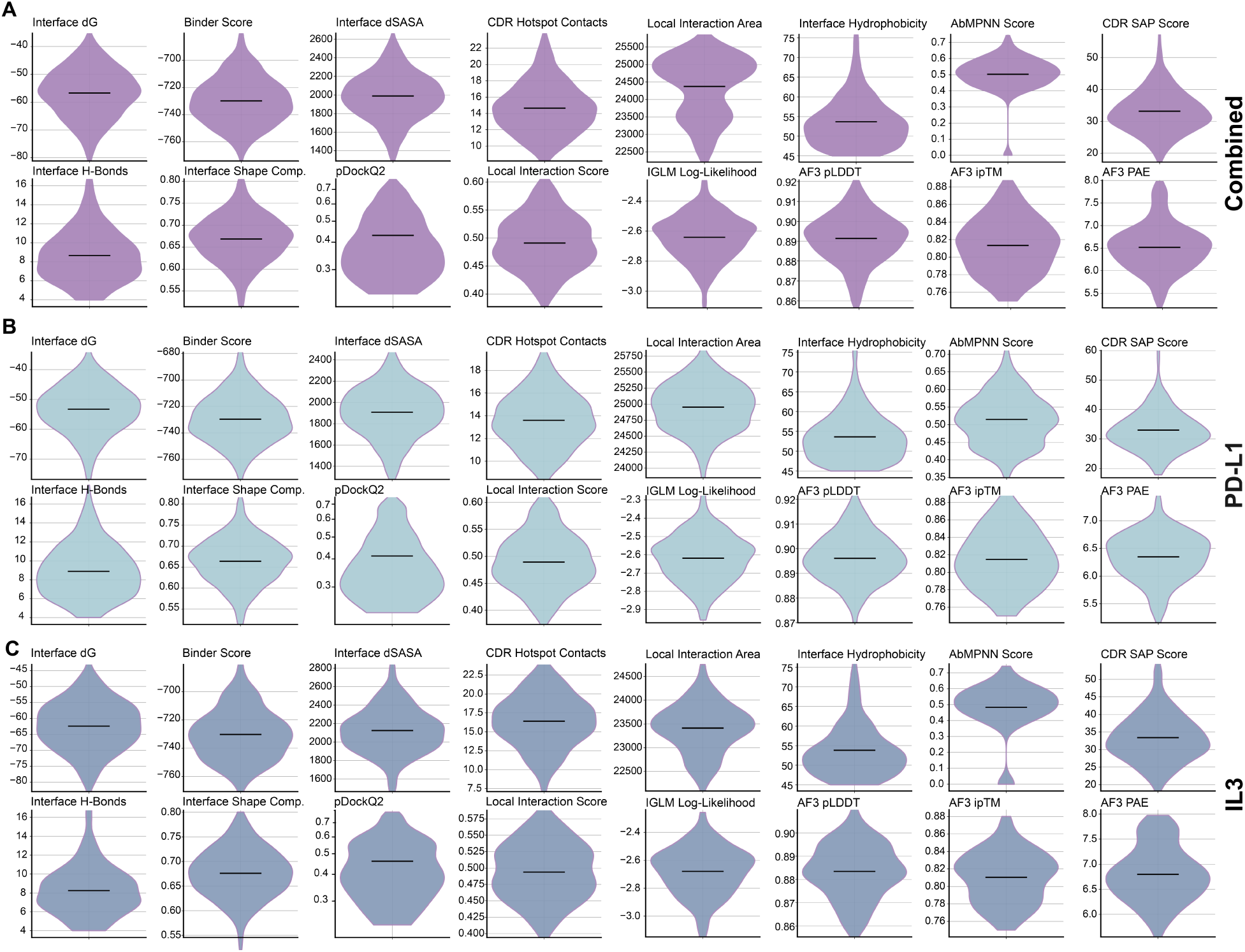
Scores distributions of key computational metrics for sampled scFv designs. (**A**) Distributions across all successful designs pooled from different targets. (**B,C**) Specific distributions for PD-L1 (**B**) and IL3 (C) designs.

**Figure S9.**
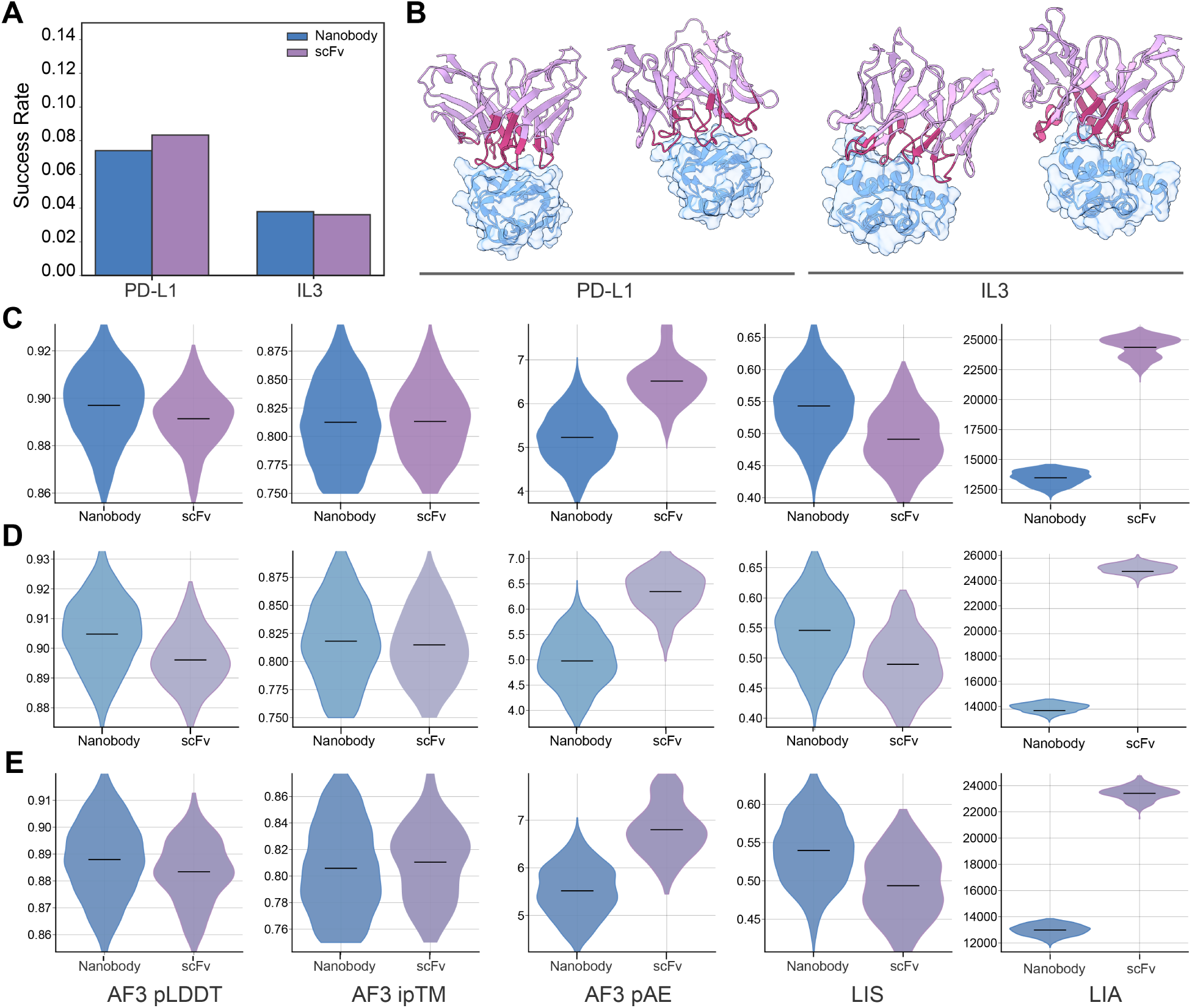
Design of scFv binders using Germinal. (**A**) The success rates (defined as the number of passing designs divided by all started trajectories) are shown for nanobody and scFv formats targeting PD-L1 and IL3. (**B**) Representative predicted binding structures of scFv binders targeting PD-L1 (left) and IL3 (right) are shown in complex with their respective antigens. (**C**) The distributions across all successful nanobody and scFv designs pooled from different targets. (**D,E**) Specific distributions for PD-L1 (**D**) and IL3 designs (**E**).

**Figure S10.**
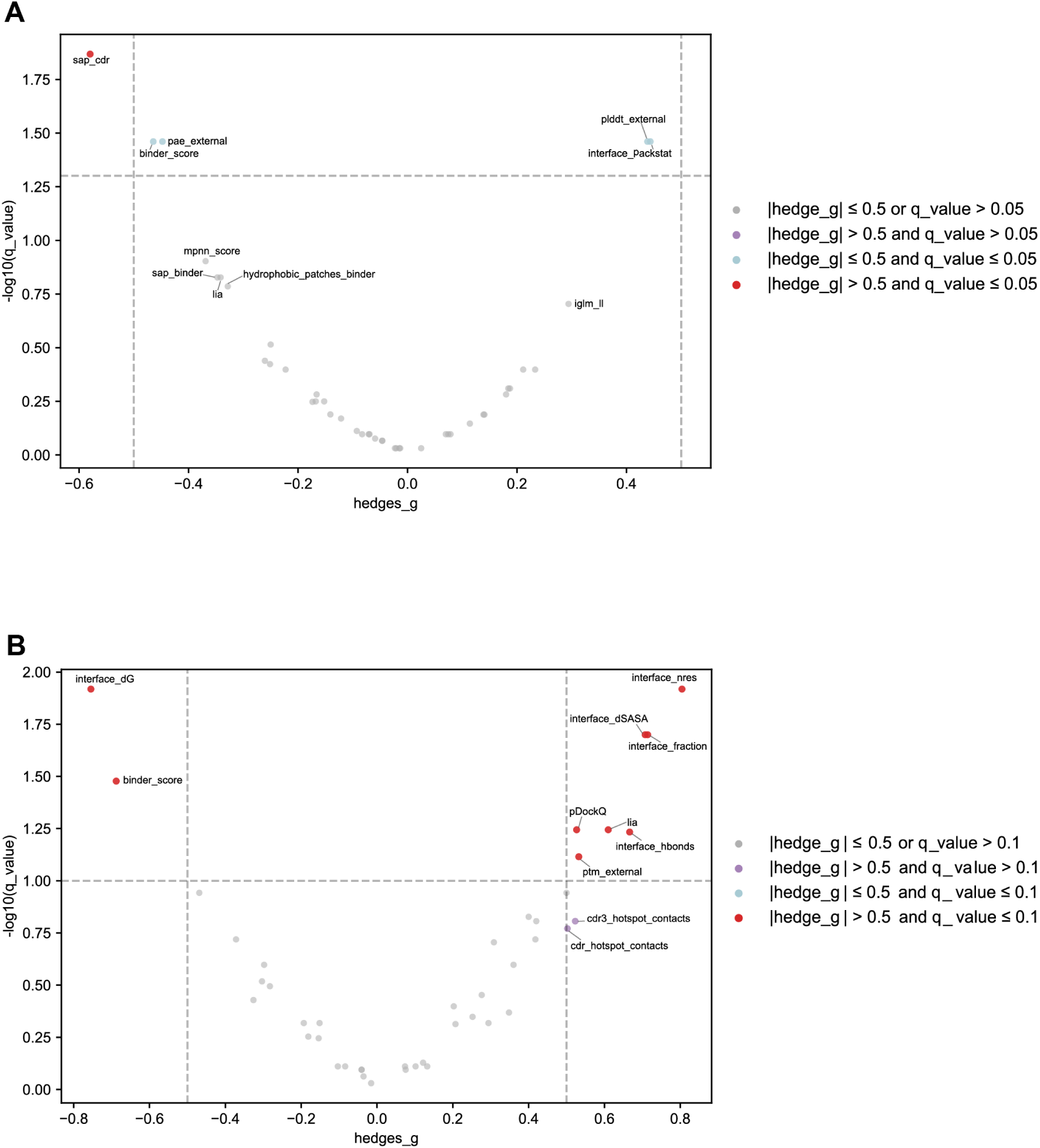
Identification of key computation metrics using differential analysis. (**A**) Two groups of designs were defined for expression analysis based on the HiBiT RLU values: top expressors (HiBiT read RLU > 1,000,000) and non-expressors (HiBiT read RLU < 150,000). Differential analysis was performed between these two groups, and metrics with Hedges’ 𝑔 > 0.5 and 𝑞 < 0.05 were considered significant (red). (**B**) Differential analysis was performed between BLI-verified binders and all other experimentally tested designs to identify key metrics that affect binding. Metrics with Hedges’ 𝑔 > 0.5 and 𝑞 < 0.1 were considered significant (red).

**Figure S11.**
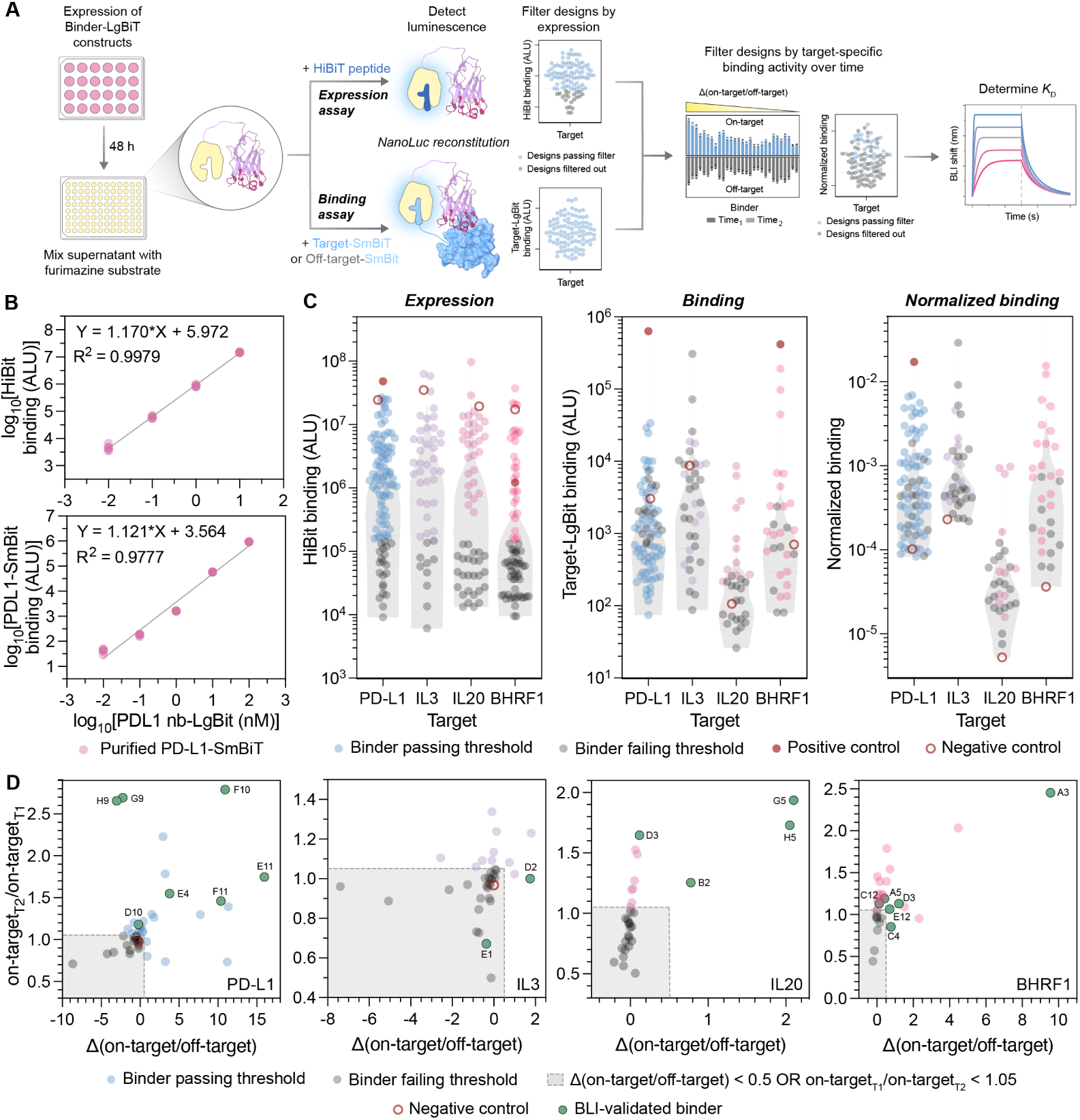
Screening expression and binding of designed binders using a split-luciferase assay. (**A**) Experimental workflow used to identify lead binders. Binder-LgBiT fusions were assayed via luminescence measurements for (i) expression by adding HiBiT peptide and (ii) binding by adding either desired target fused to SmBiT protein (‘on-target’ binding), or an off-target protein-SmBiT fusion (‘off-target’ binding). Designs with favorable expression and binding activity in this assay were further characterized with BLI. (**B**) *Top:* expression standard curve. log_10_–log_10_ linear relationship between HiBiT-LgBiT produced luminescence reads and binder concentration (fit and 𝑅^2^ shown). *Bottom:* binding standard curve. log_10_–log_10_ linear relationship between PD-L1–SmBiT luminescence reads and concentration of a known PD-L1 nanobody–LgBiT (PDB: 5JDS; reported 𝐾_𝐷_ ≈ 3 nM; fit and 𝑅^2^ shown). (**C**) *Left:* designed binder expression, shown as raw luminescence (arbitrary luminescence units, ALU). Designs shown in gray fall below the expression threshold (raw HiBiT signal > 150,000 ALU). *Middle:* Binding activity for all designs that passed the expression threshold, shown as raw luminescence. *Right:* expression-normalized binding estimates used for identification of lead binder candidates. The positive control nanobody against PD-L1, or miniprotein against BHRF1, is shown as a solid red circle, while negative control nanobody against TNF𝛼 (for PD-L1 and IL3 screens) or against PD-L1 (for IL20 and BHRF1 screens) is indicated by a hollow red circle. Designs shown in gray failed to meet the binding threshold. (C) High expressing binders were then selected if the difference in the [normalized on-target]/[normalized off-target binding] ratio between T_1_ and T_2_ was greater than 0.5, or if the ratio of normalized on-target binding (on-target T_2_/on-target T_1_) was greater than 1.05 (**Methods**). Passing candidates, shown as colored circles, were subsequently analyzed by biolayer interferometry (BLI). Green circles represents designs which were validated as hits by BLI. Candidates that fall below the binding threshold (shaded gray region) are shown as gray circles. The average normalized binding value (0.34), [normalized on-target]/[normalized off-target binding] ratio (7.2), and ratio of normalized on-target binding (456) of the BHRF1 positive control exceeded the range of the tested designs and these values were thus omitted from the plots for clarity.

**Figure S12.**
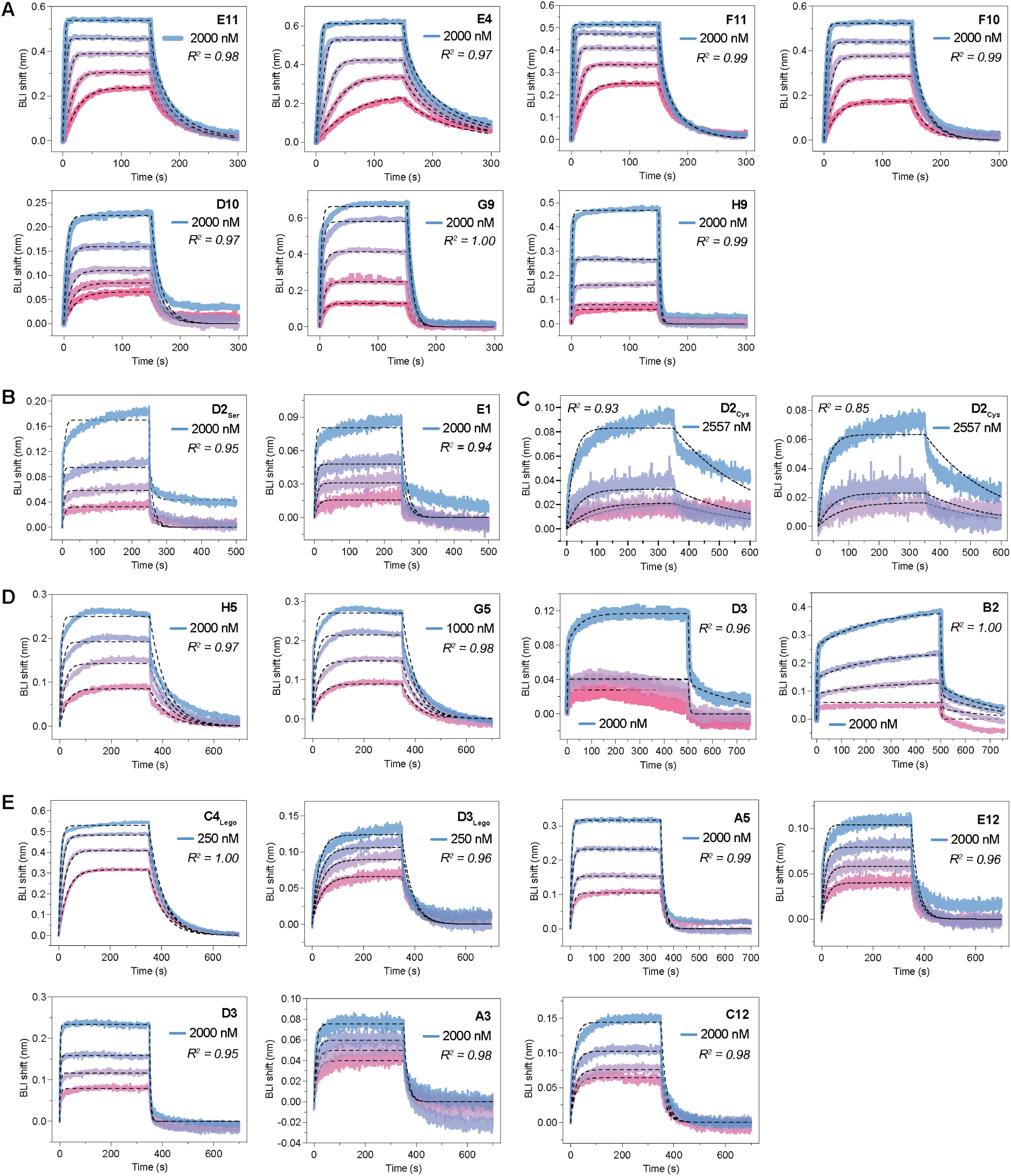
BLI sensorgrams for validated nanobodies against all four target antigens. (**A-E**) Sensorgrams for all 7 PD-L1 designs (**A**), IL3 E1 and D2_Ser_(**B**), IL3 D2_Cys_ design (**C**), all 4 IL20 designs (**D**), and all 7 BHRF1 designs, including D3_Lego_ and C4_Lego_ (**E**). D2_Ser_ is a variant of the designed IL3 binder D2 in which the CDR cysteine is mutated to serine; D2_Cys_ is the originally designed D2 sequence. For each BLI experiment, a titration of 4–5 concentrations was prepared by 2-fold serial dilution from the highest concentration tested, indicated on each plot. For all BLI plots, binding curves are shown as colored lines, and kinetic fits are shown as black dashed lines. All curves were fit with a global 1:1 binding model. 𝑅^2^ values are indicated on each plot.

**Figure S13.**
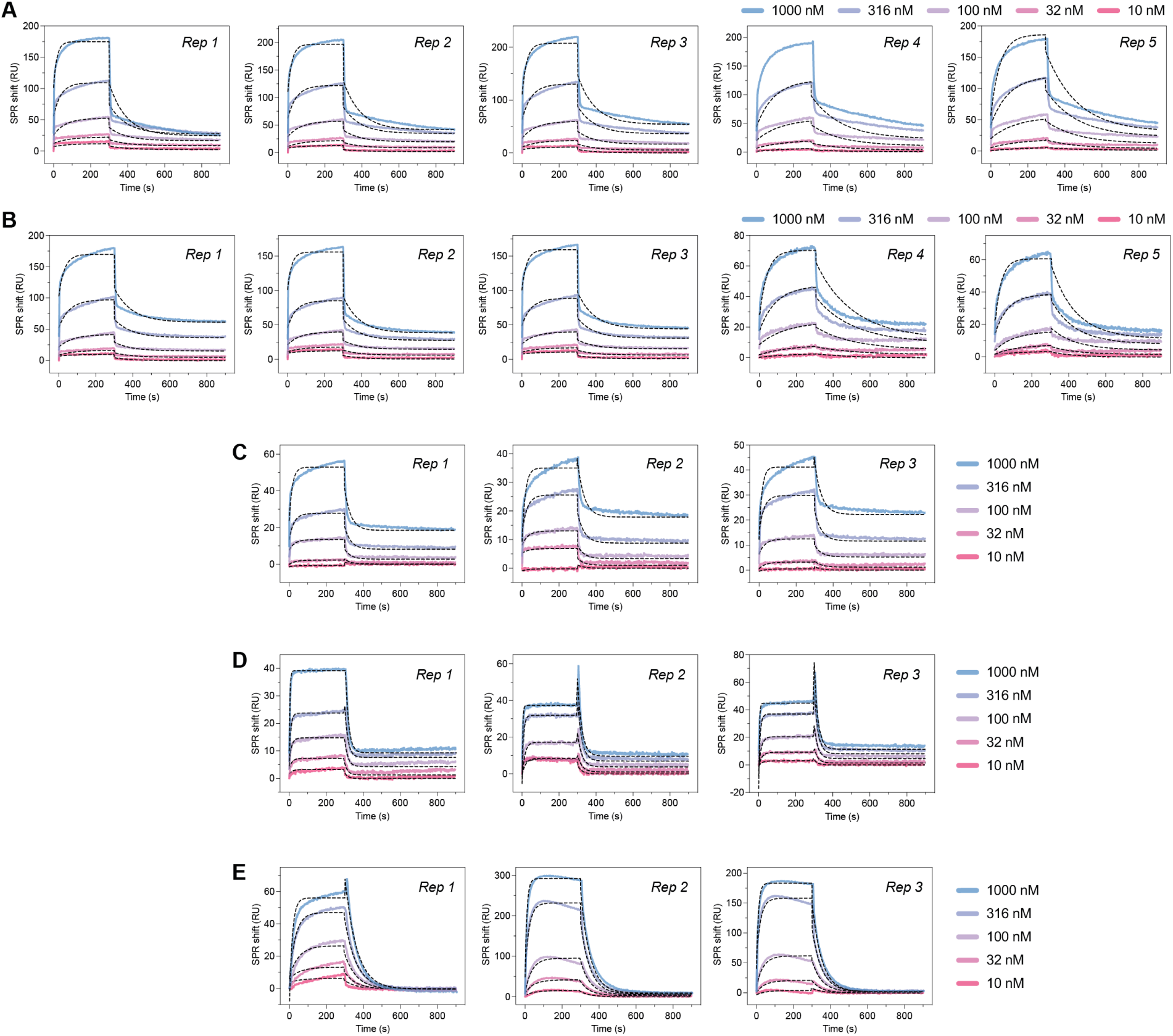
Replicate SPR sensorgrams for designed scFv hits against PD-L1 and IL3. SPR sensorgrams for independent replicates of the anti-PD-L1 scFvs H5 (**A**), G5 (**B**), E5 (**C**), and E2 (**D**), or for the anti-IL3 scFv F4 (**E**). For all experiments, a five-point half-log serial dilution series was prepared from a top concentration of 1000 nM. Binding curves are shown as colored lines and kinetic fits as black dashed lines.

**Figure S14.**
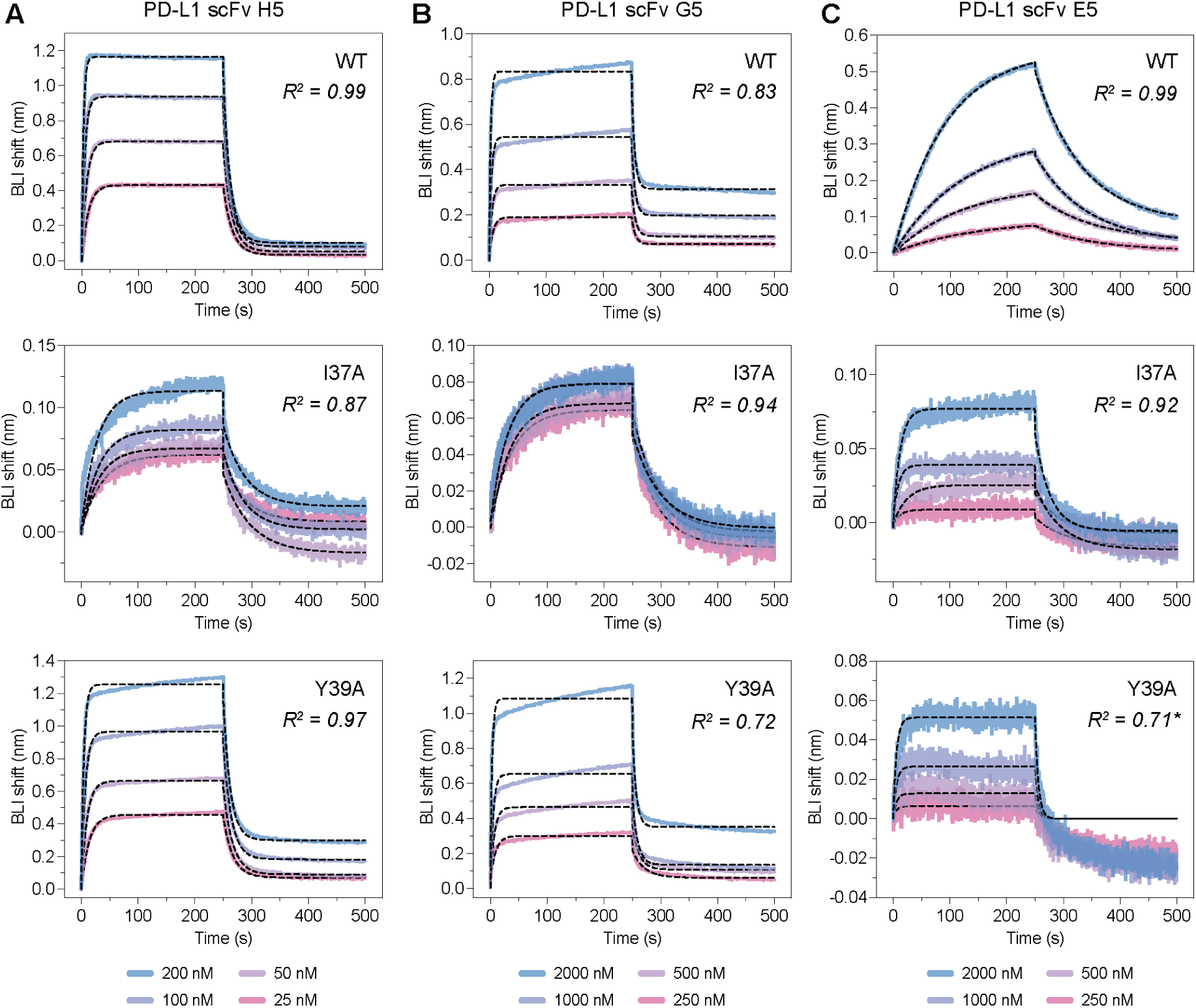
BLI sensorgrams for wild-type and alanine variant PD-L1 binding by designed scFvs. (**A–C**) BLI sensorgrams for designed anti-PD-L1 scFvs H5 (**A**), G5 (**B**), and E5 (**C**) titrated against wild-type PD-L1 (top), and alanine substitution variants I37A (middle) and Y39A (bottom). Binding curves are colored by analyte concentration and kinetic fits are shown as black dashed lines. All curves are fit with a global 1:1 binding model. 𝑅^2^ values are indicated on each plot. An asterisk (*) denotes sensorgrams where the maximum BLI shift at the highest analyte concentration tested did not exceed 0.07 nm; these were classified as non-binders.

**Figure S15.**
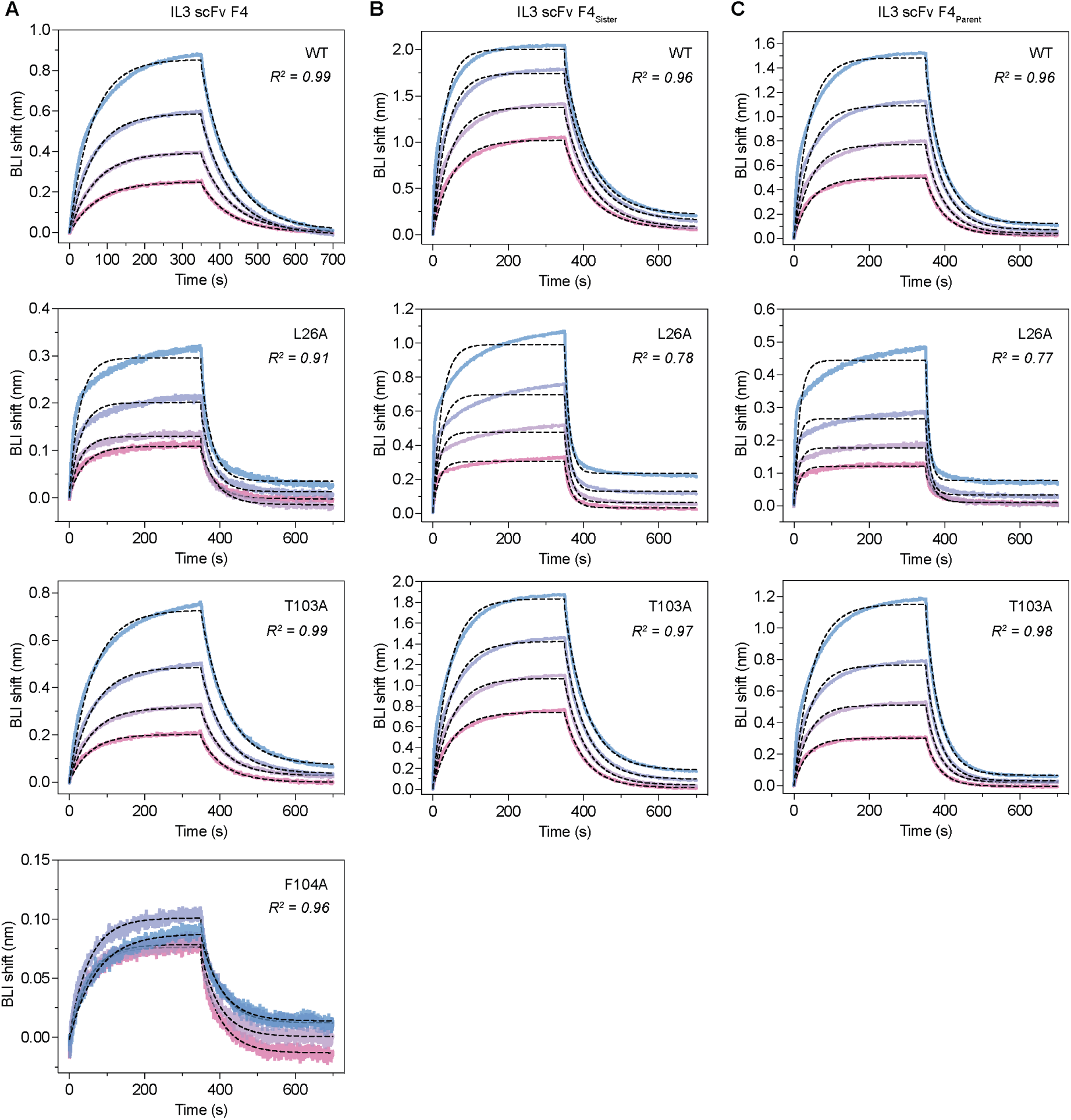
BLI sensorgrams for IL3 alanine variant binding by designed scFv and positive control antibody. (**A**) BLI sensorgrams for the original anti-IL3 F4 Fab titrated against wild-type IL3 and alanine substitution variants L26A, T103A, and F104A. (**B, C**) BLI sensorgrams for related anti-IL3 Fabs F4_Sister_ (**B**) and F4_Parent_ (**C**) titrated against wild-type IL3 and alanine substitution variants L26A and T103A. Binding curves are colored by analyte concentration and kinetic fits are shown as black dashed lines. All curves are fit with a global 1:1 binding model. 𝑅^2^ values are indicated on each plot.

**Figure S16.**
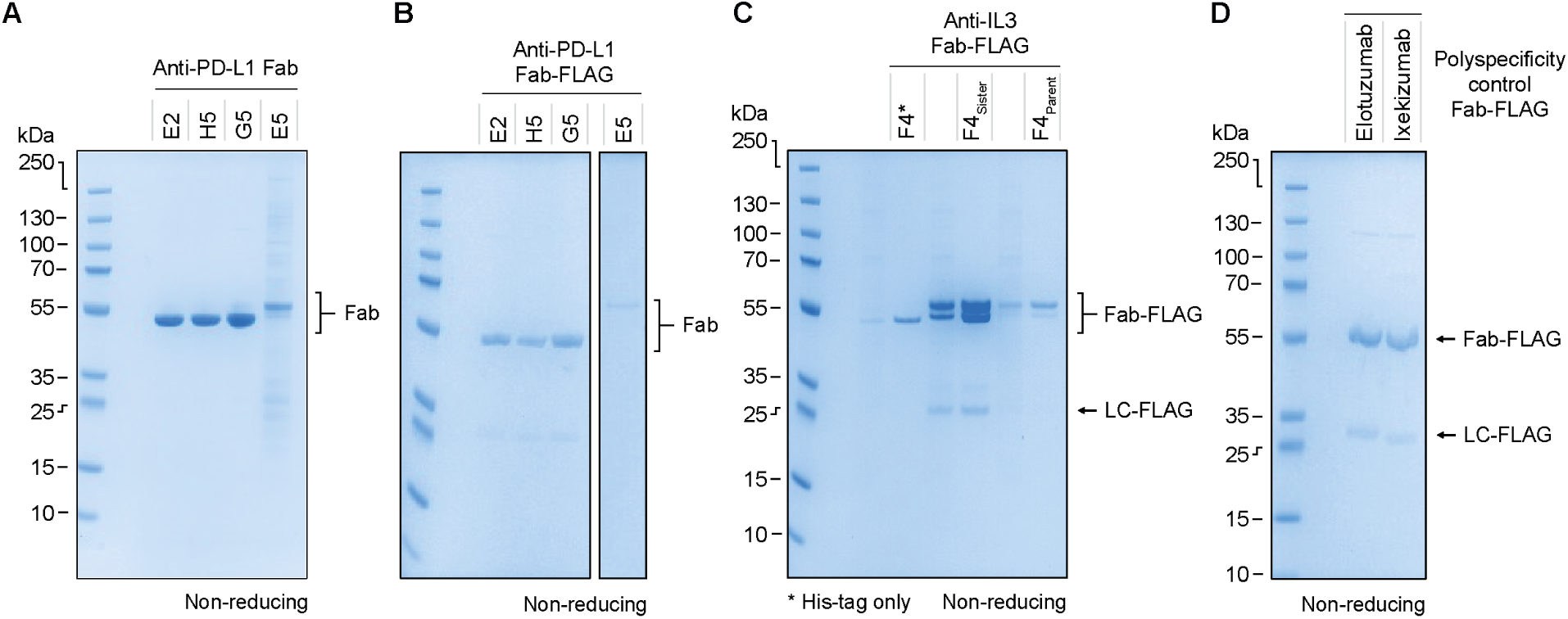
SDS-PAGE analysis of purified designed Fabs and control binders. Non-reducing SDS-PAGE gels of purified Fabs used in this study. (**A**) Anti-PD-L1 Fabs (His-tagged); E2, H5, G5, and E5. (**B**) Anti-PD-L1 Fabs (His- and FLAG-tagged); E2, H5, and G5. (**C**) Anti-IL3 Fabs (His- and FLAG-tagged); F4, F4_Sister_, and F4_Parent_. Intact Fab-FLAG and free light chain-FLAG (LC-FLAG) bands are indicated. (**D**) Polyspecificity assay control Fabs (His- and FLAG-tagged); Elotuzumab and Ixekizumab. Intact Fab-FLAG and LC-FLAG bands are indicated. Molecular weight markers (kDa) are shown on the left of each gel.

**Figure S17.**
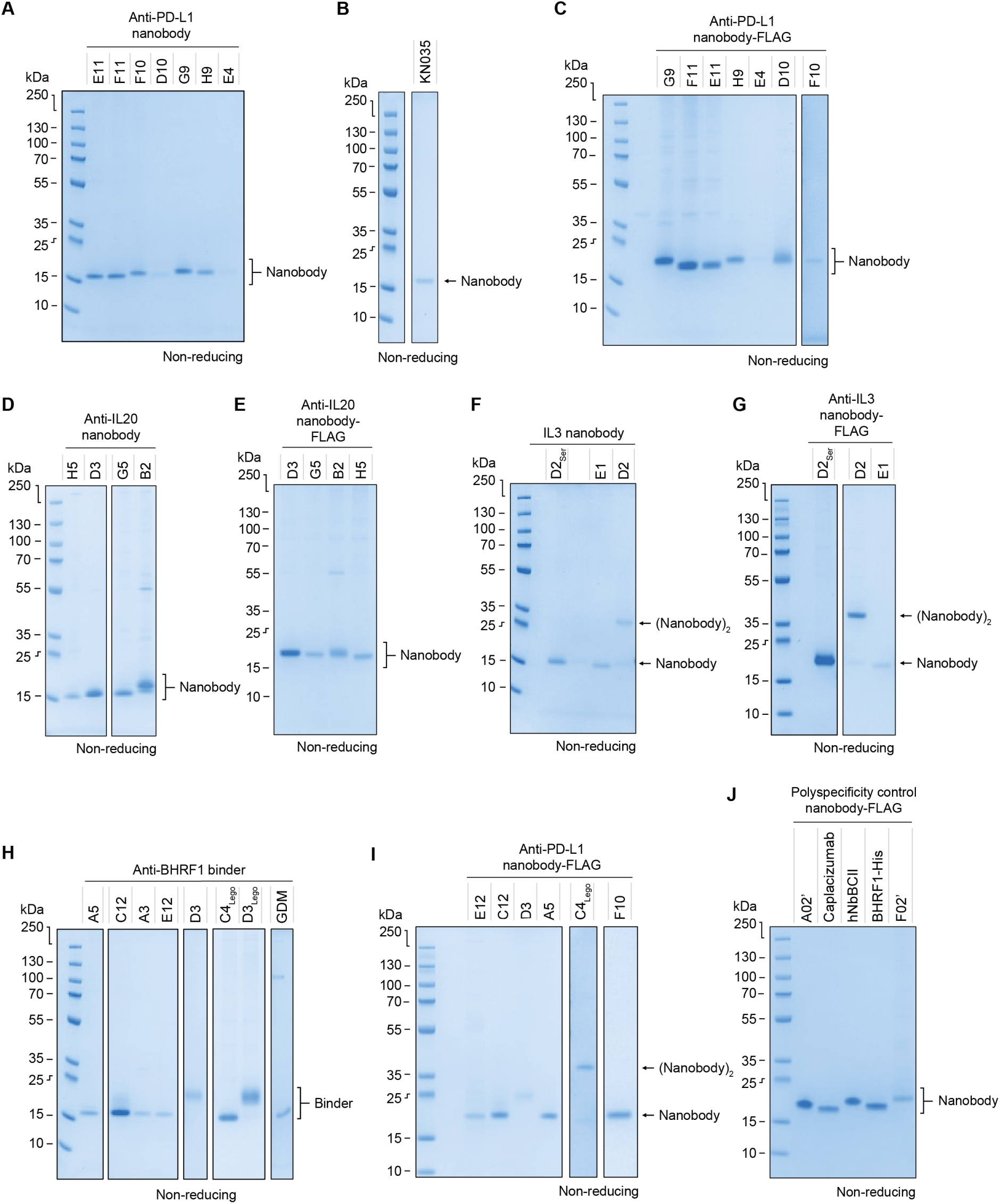
SDS-PAGE analysis of purified designed nanobodies and control binders. Non-reducing SDS-PAGE gels of purified nanobodies and binders used in this study. (**A**) Anti-PD-L1 nanobodies (His-tagged). (**B**) Anti-PD-L1 positive control nanobody KN035. (**C**) Anti-PD-L1 designed nanobodies (His- and FLAG-tagged). (**D**) Anti-IL20 designed nanobodies (His-tagged). (**E**) Anti-IL20 designed nanobodies (His- and FLAG-tagged). (**F**) Anti-IL3 designed nanobodies (His-tagged); monomeric nanobody and disulfide-linked dimer bands are indicated. (**G**) Anti-IL3 designed nanobodies (His- and FLAG-tagged); monomeric and dimeric species indicated. (**H**) Anti-BHRF1 binders (His-tagged), including GDM_BHRF1_35 control. (**I**) Anti-PD-L1 designed nanobodies (His- and FLAG-tagged); monomeric and dimeric species indicated. (**J**) Polyspecificity assay control proteins (His- and FLAG-tagged). Molecular weight markers (kDa) are shown on the left of each gel.

**Figure S18.**
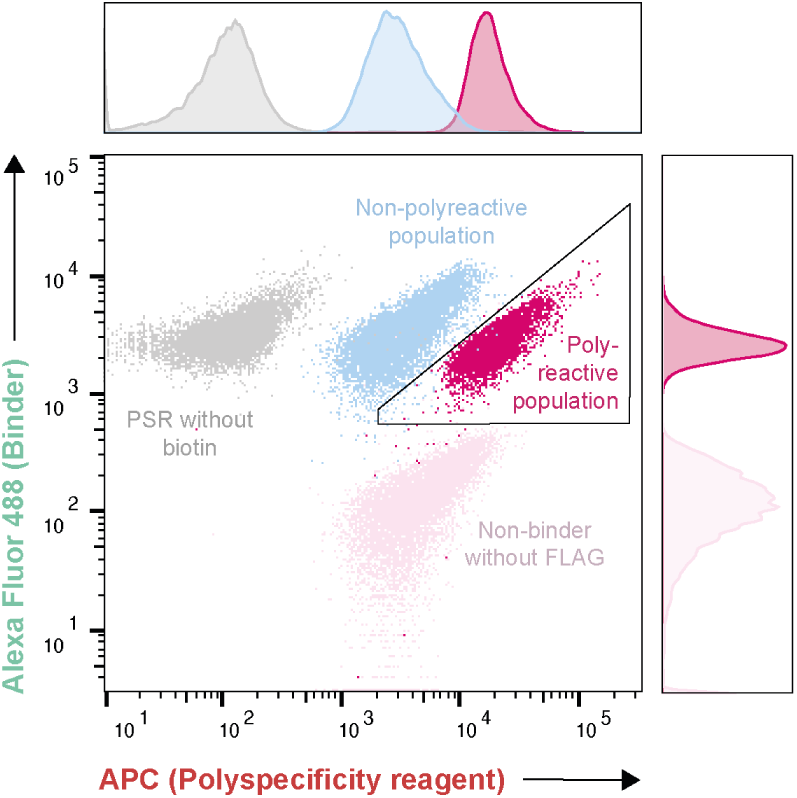
Determining polyreactivity of designed binders with the polyspecificity particle assay. Representative flow cytometry scatter plot from the polyspecificity particle assay. His- and FLAG-taggd binders captured on anti-His magnetic beads are detected on the Alexa Fluor 488 axis (via anti-FLAG antibody); APC fluorescence reports binding to biotinylated polyspecificity reagent (PSR). Control populations are shown for the positive control binder incubated with un-biotinylated PSR (’PSR without biotin’), and for a non-FLAG-tagged, non-binder incubated with biotinylated PSR (BHRF1-His; ‘non-binder without FLAG’). Polygonal gates delineate negative control (non-polyspecific) and positive control (polyspecific) populations, which establish gating boundaries. Marginal density plots show the distribution of each population along both axes.

**Figure S19.**
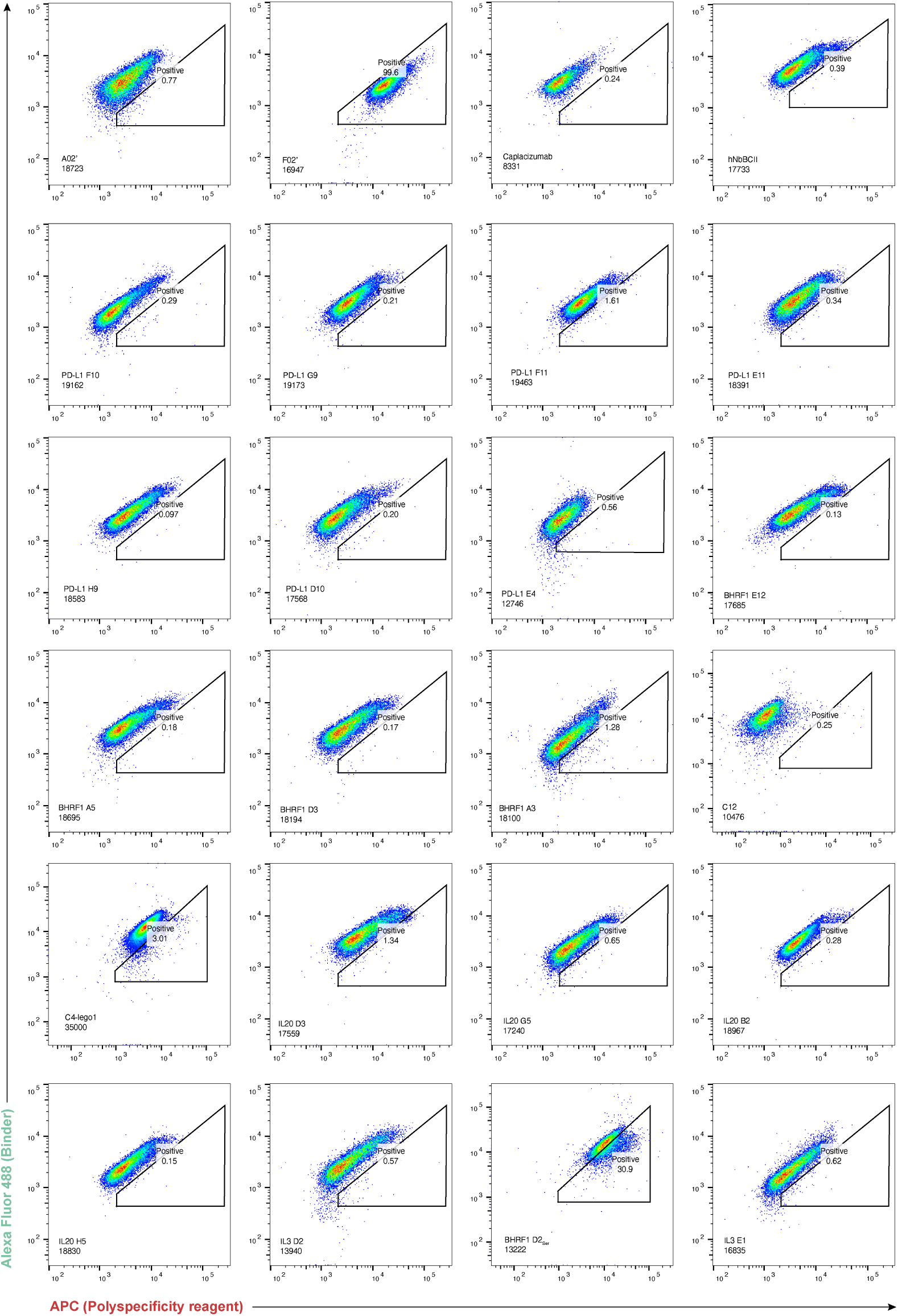
Raw flow cytometry data from the polyspecificity particle assay for designed nanobodies. Representative scatter plots for each nanobody hit, showing Alexa Fluor 488 (binder loading, via anti-FLAG antibody) on the 𝑦-axis and APC (biotinylated polyspecificity reagent binding) on the 𝑥-axis. Each data point represents an individual bead. The polygonal gate defining the polyspecific population was set using the positive control F02’, and the percentage of beads falling within this gate (*Positive*) is indicated for each sample. The number below each label indicates the total bead count acquired. Reference controls are shown in the first row: A02’ and Caplacizumab (non-polyspecific), F02’ (polyspecific), and hNbBCII (humanized nanobody whose framework was used in germinal nanobody design). Designed nanobodies are ordered by target.

**Figure S20.**
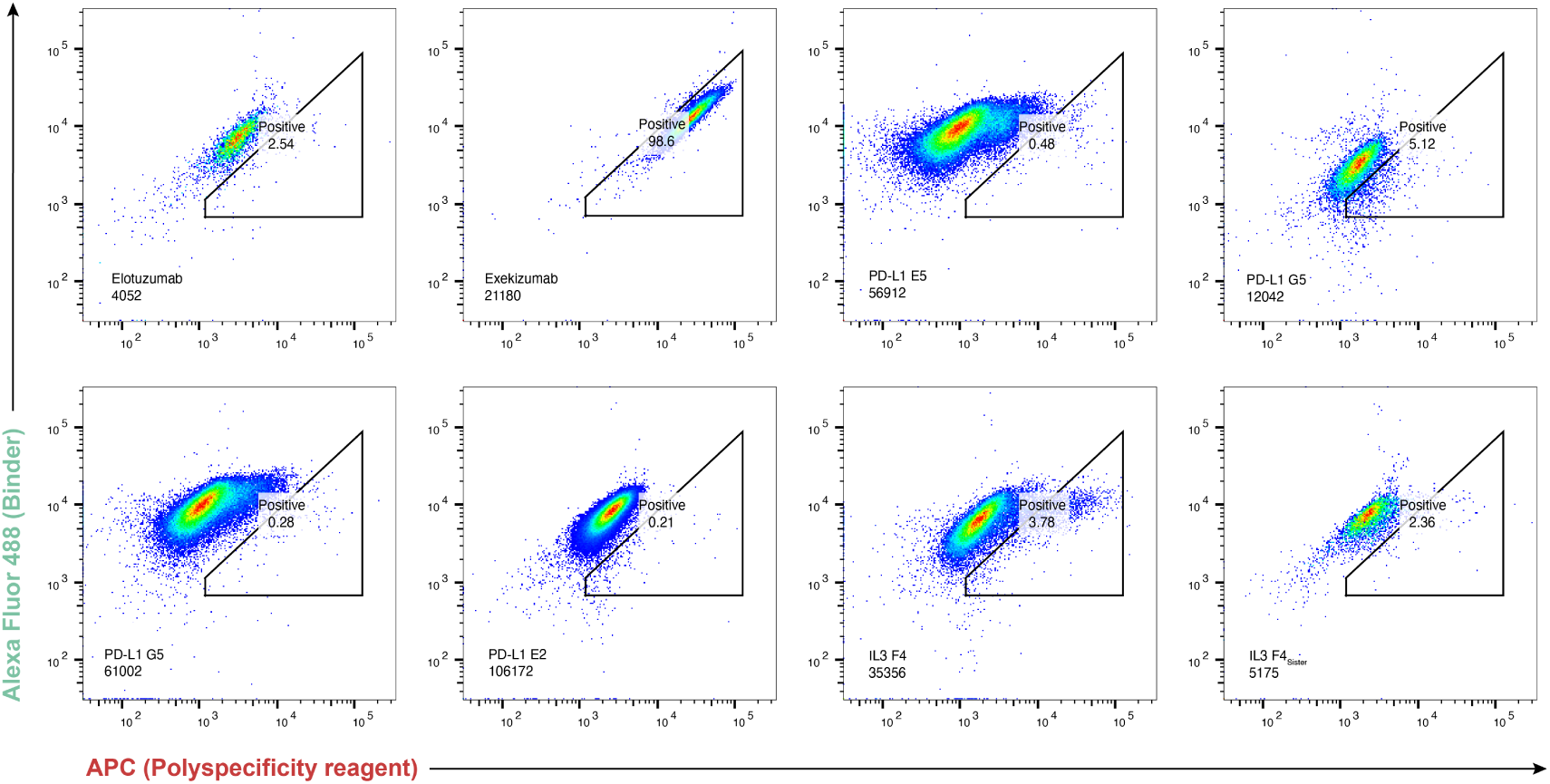
Raw flow cytometry data from the polyspecificity particle assay for designed scFvs. Representative scatter plots for each Fab tested, showing Alexa Fluor 488 (binder loading, via anti-FLAG antibody) on the 𝑦-axis and APC (biotinylated polyspecificity reagent binding) on the 𝑥-axis. Each data point represents an individual bead. The polygonal gate defining the polyspecific population was set using the positive control Ixekizumab, and the percentage of beads falling within this gate (*Positive*) is indicated for each sample. The number below each binder label indicates the total bead count acquired. Elotuzumab and Ixekizumab serve as non-polyspecific and polyspecific reference controls, respectively.

**Figure S21.**
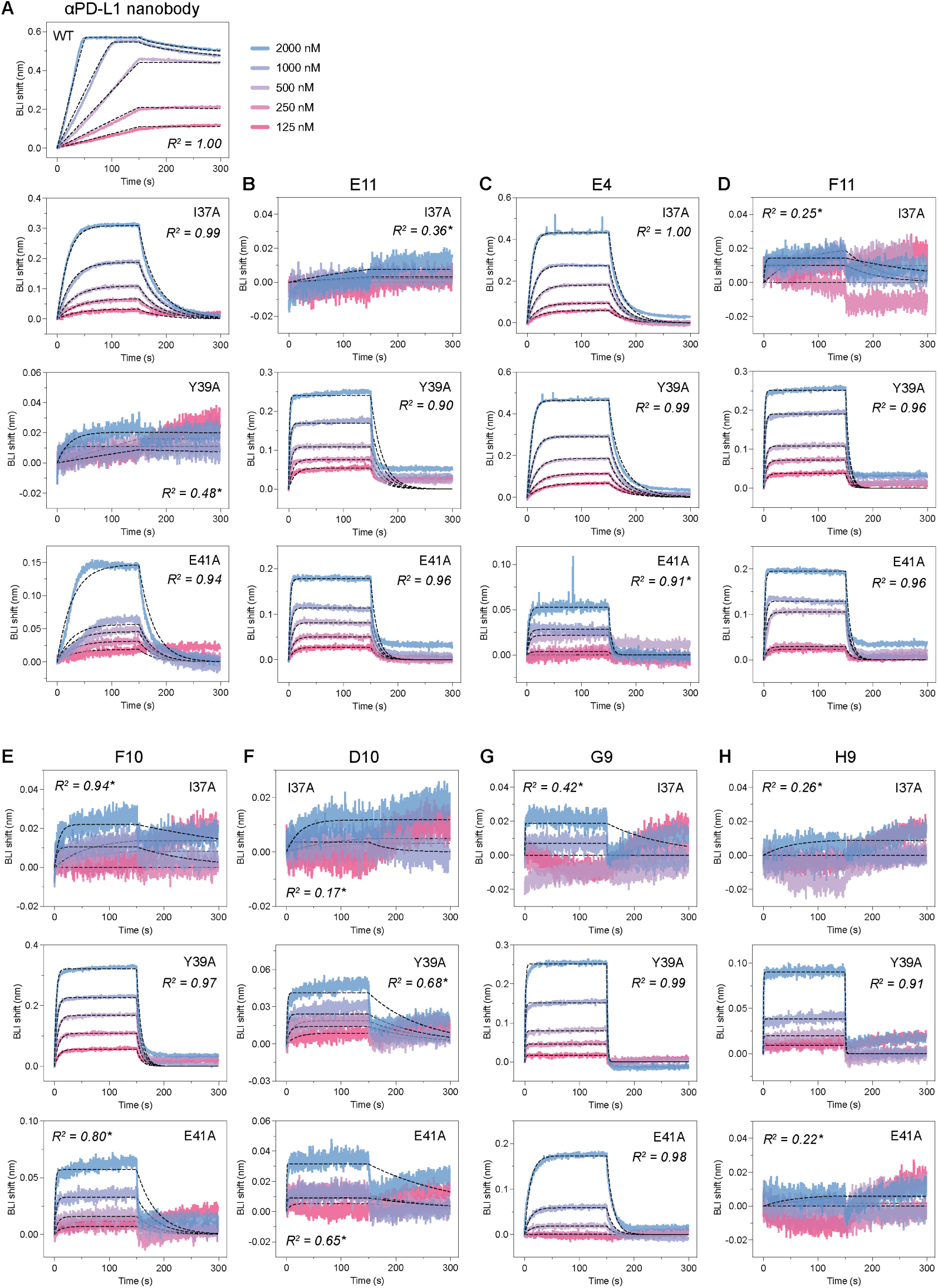
BLI sensorgrams for PD-L1 alanine variant binding by designed and positive control nanobody. (**A**) BLI sensorgrams for a positive control anti-PD-L1 nanobody titrated against wild-type PD-L1 and alanine variants I37A, Y39A, and E41A. (**B–H**) BLI sensorgrams for Germinal-designed anti-PD-L1 nanobodies titrated against PD-L1 alanine variants I37A (top), Y39A (middle), and E41A (bottom). All binding curves are coloured by analyte concentration as indicated in (**A**) and kinetic fits are shown as black dashed lines. All curves are fit with a global 1:1 binding model. 𝑅^2^ values are indicated on each plot. An asterisk (*) denotes sensorgrams where the maximum BLI shift at the highest analyte concentration tested did not exceed 0.07 nm; these were classified as non-binders.

**Figure S22.**
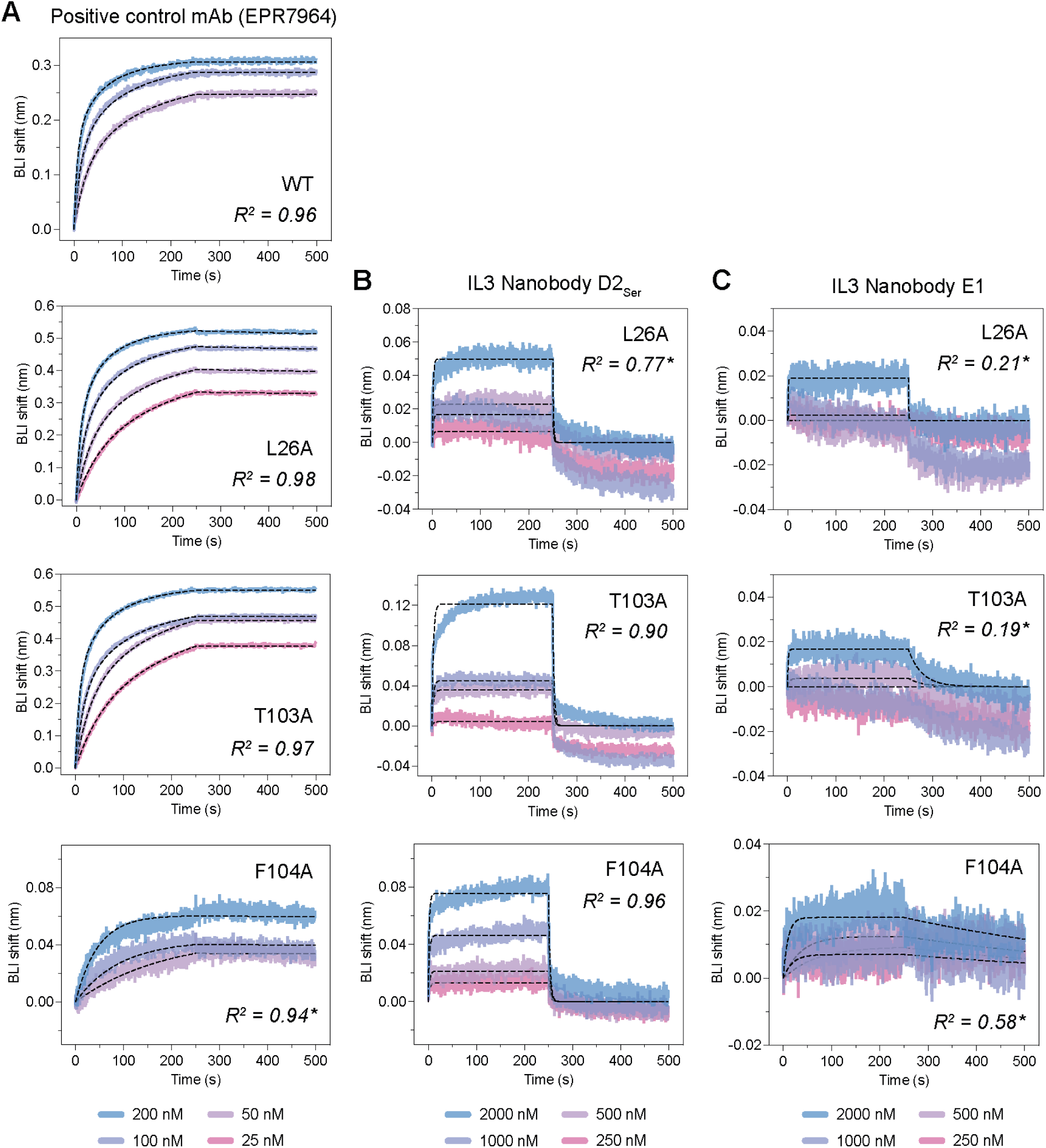
BLI sensorgrams for IL3 alanine variant binding by designed nanobodies and positive control antibody. (**A**) BLI sensorgrams for the positive control anti-IL3 monoclonal antibody EPR7964 titrated against wild-type IL3 and alanine substitution variants L26A, T103A, and F104A. (**B, C**) BLI sensorgrams for designed IL3 nanobodies D2_Ser_ (**B**) and E1 (**C**) titrated against the same IL3 alanine variants. Binding curves are colored by analyte concentration and kinetic fits are shown as black dashed lines. IgG positive control curves are fit with a global 1:2 bivalent analyte model, whilst nanobody curves are fit with a global 1:1 binding model. 𝑅^2^ values are indicated on each plot. An asterisk (*) denotes sensorgrams where the maximum BLI shift at the highest analyte concentration tested did not exceed 0.07 nm; these were classified as non-binders.

**Figure S23.**
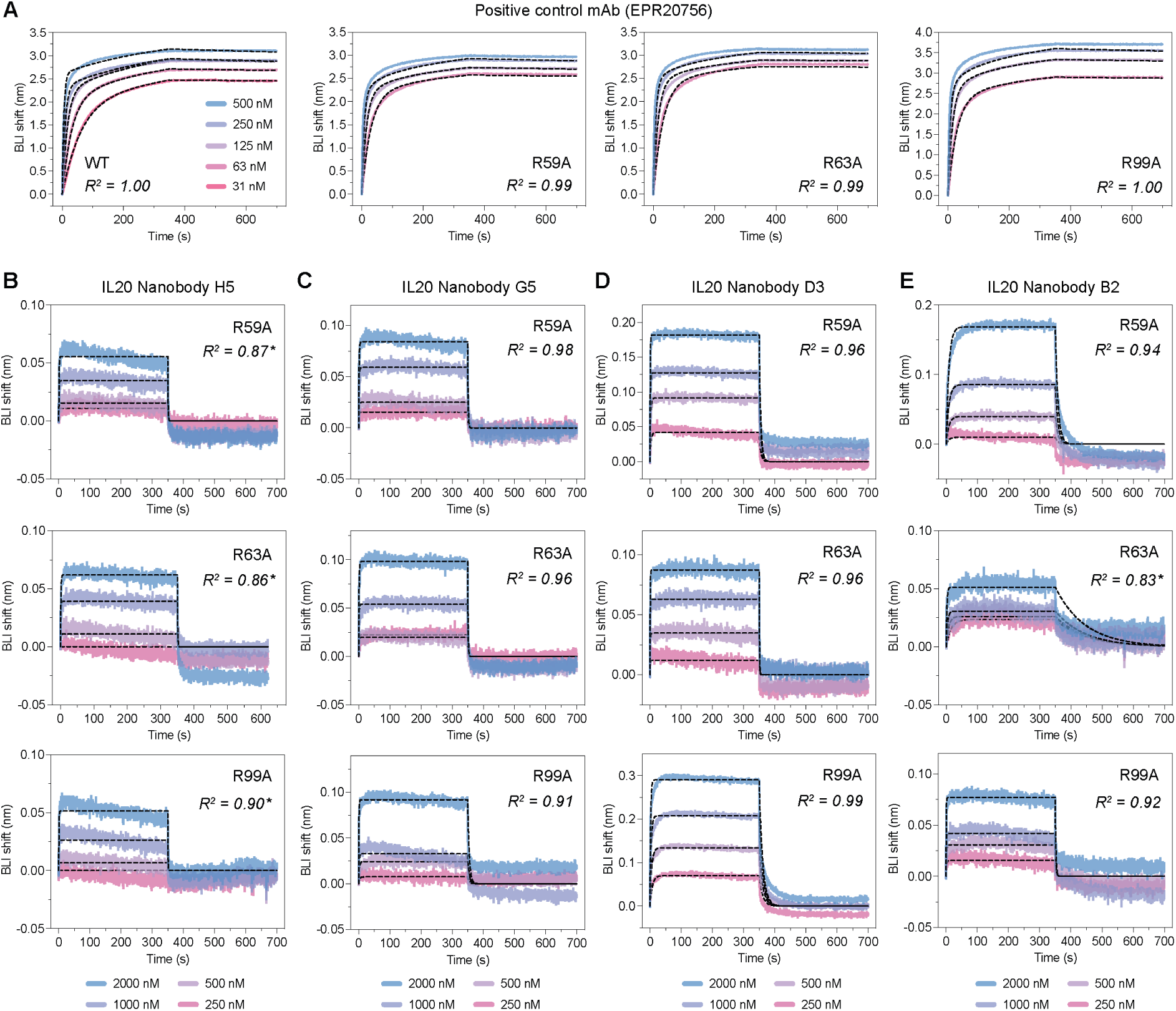
BLI sensorgrams for IL20 alanine variant binding by designed nanobodies and positive control antibody. (**A**) BLI sensorgrams for the positive control anti-IL20 monoclonal antibody EPR20756 titrated against wild-type IL20 and alanine substitution variants R59A, R63A, and R99A. (**B–E**) BLI sensorgrams for designed IL20 nanobodies H5 (**B**), G5 (**C**), D3 (**D**), and B2 (**E**) titrated against IL20 alanine variants R59A, R63A, and R99A. Binding curves are colored by analyte concentration and kinetic fits are shown as black dashed lines. IgG positive control curves are fit with a global 1:2 bivalent analyte model, whilst nanobody curves are fit with a global 1:1 binding model. 𝑅^2^ values are indicated on each plot. An asterisk (*) denotes sensorgrams where the maximum BLI shift at the highest analyte concentration tested did not exceed 0.07 nm; these were classified as non-binders.

**Figure S24.**
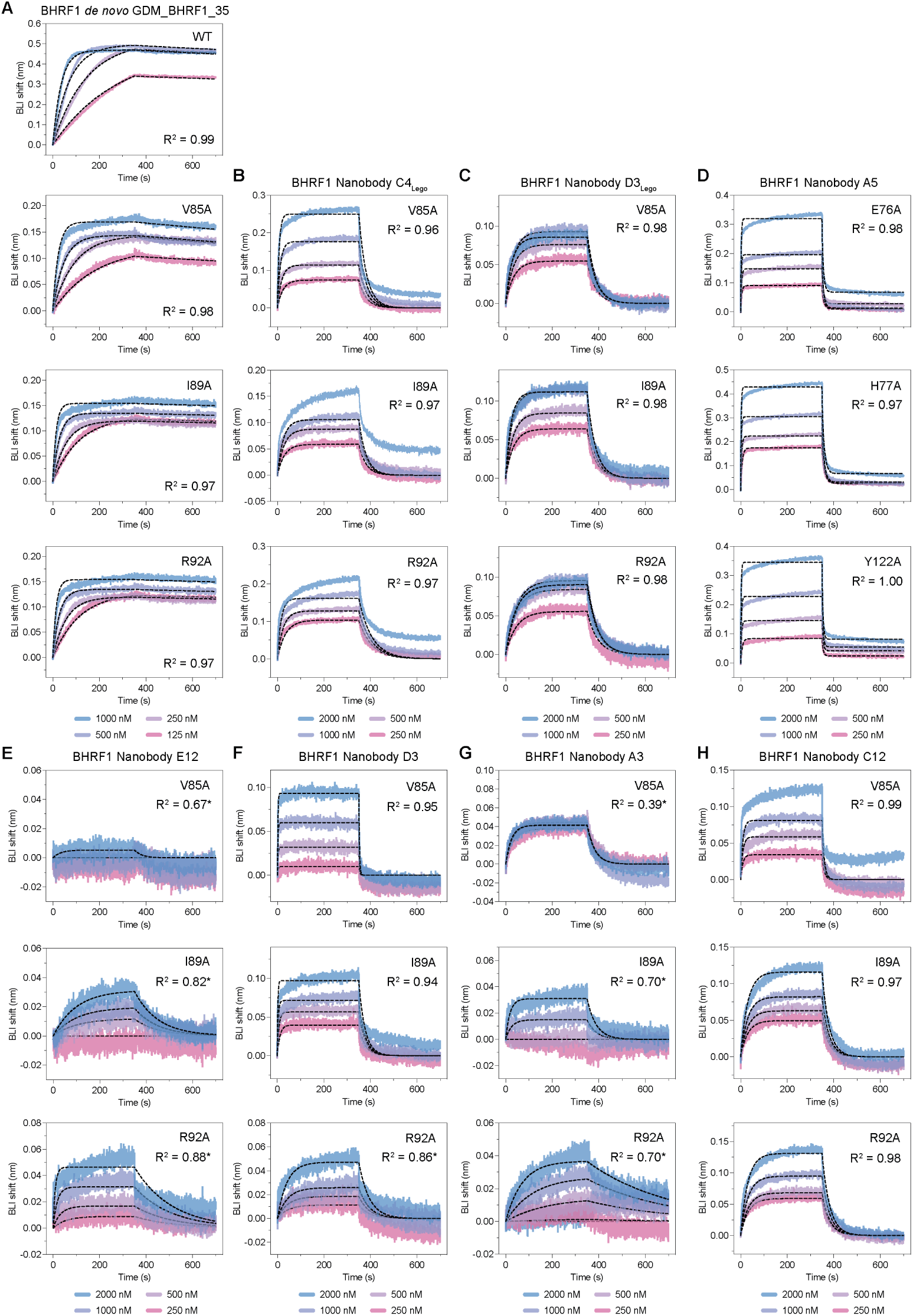
BLI sensorgrams for BHRF1 alanine variant binding by designed nanobodies and positive control binder. (**A**) BLI sensorgrams for the positive control binder GDM_BHRF1_35 against wild-type BHRF1. (**B–D**) BLI sensorgrams for nanobody hits C4_Lego_ (**B**), D3_Lego_ (**C**), and A5 (**D**) against BHRF1 alanine variants V85A, I89A, and R92A (B, C) or equivalent positions E76A, H77A, and Y122A (D). (**E–H**) BLI sensorgrams for nanobody hits E12 (**E**), D3 (**F**), A3 (**G**), and C12 (**H**) against BHRF1 alanine variants V85A, I89A, and R92A. Binding curves are colored by analyte concentration and kinetic fits are shown as black dashed lines. All curves are fit with a global 1:1 binding model. 𝑅^2^ values are indicated on each plot. An asterisk (*) denotes sensorgrams where the maximum BLI shift at the highest analyte concentration tested did not exceed 0.07 nm; these were classified as non-binders.

**Figure S25.**
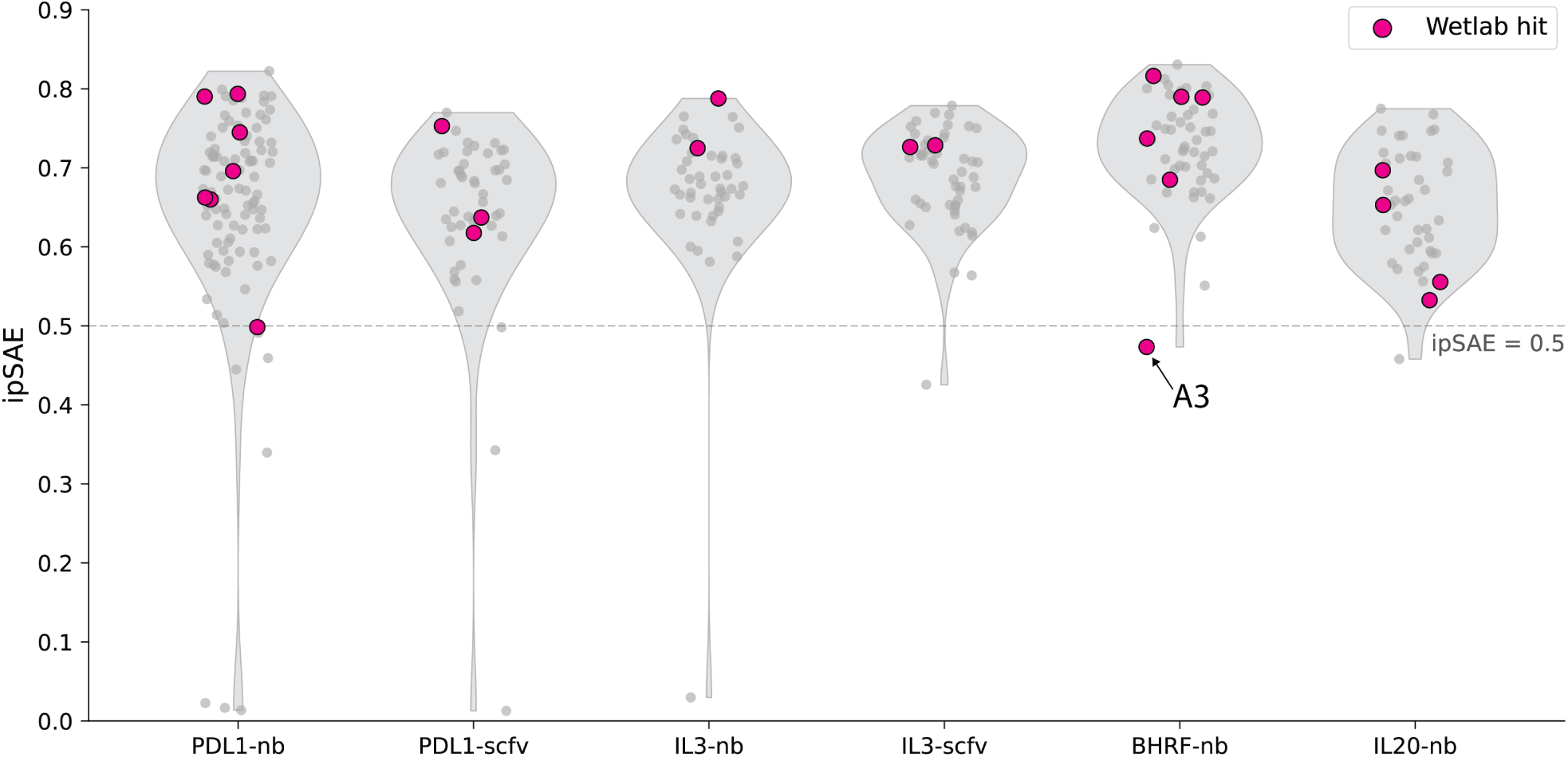
Retrospective ipSAE analysis of experimentally tested designs. Distribution of interface predicted self-assessment estimate (min_ipSAE) scores for all experimentally tested designs across each target–binder format combination. Each point represents an individual design; experimentally validated hits are highlighted in magenta. The dashed line indicates a threshold of 0.5.

**Figure S26.**
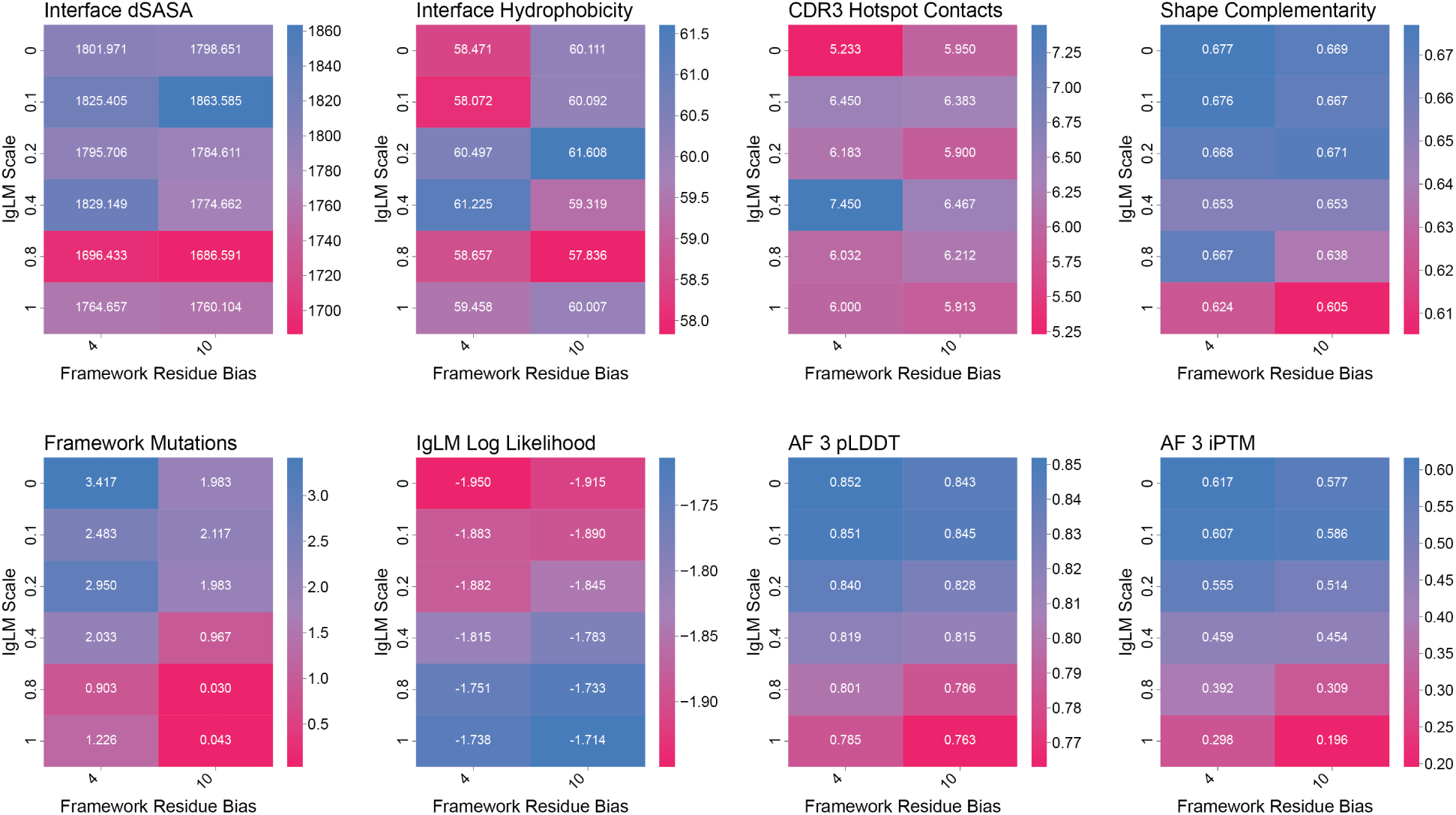
Comparison heatmap of selected metrics across a sweep of framework sequence biases and IgLM scale values. Each cell reports the mean value of the corresponding metric over a small-scale run of 60 trajectories. Metrics in the top row show little dependence on either variable, whereas those in the bottom row exhibit clear trends as Sequence bias and IgLM scale vary.

**Figure S27.**
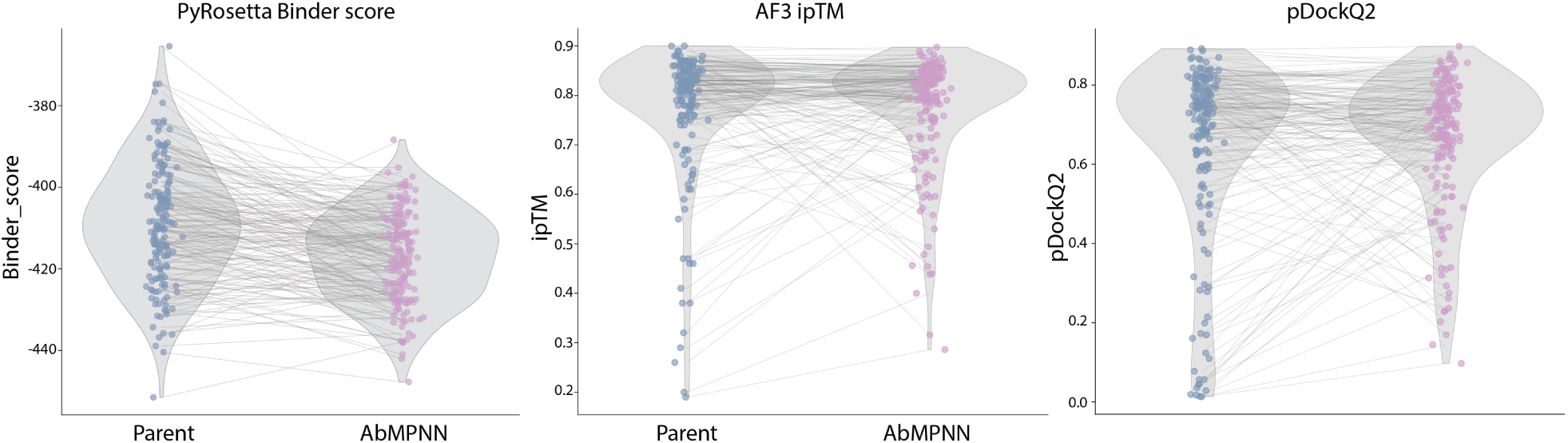
Effect of AbMPNN sequence redesign on key representative metrics. Paired violin plots showing the distribution of PyRosetta Binder score (left), AlphaFold3 interface predicted TM-score (ipTM; middle), and pDockQ2 (right) for Germinal parent designs (blue circles) and their corresponding AbMPNN-redesigned daughter sequences (pink circles). Lines connect paired parent–AbMPNN sequence pairs. Parent designs with high AF3 ipTM and pDockQ2 scores also produce high-confidence daughter designs.

**Figure S28.**
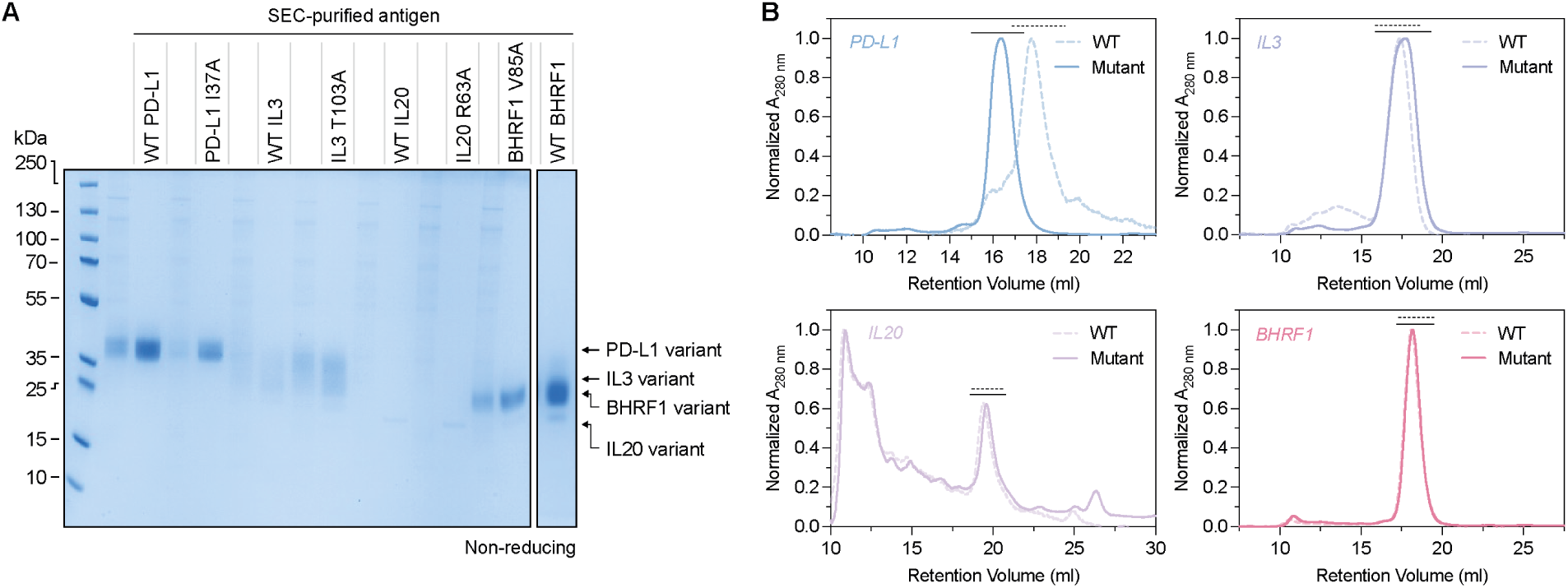
Quality control of purified wild-type and alanine variant antigens. (**A**) Non-reducing SDS-PAGE gel of SEC-purified wild-type and representative alanine substitution variants for each target antigen: PD-L1 (WT and I37A), IL3 (WT and T103A), IL20 (WT and R63A), and BHRF1 (WT and V85A). (**B**) Analytical SEC traces (normalized A_280_) for wild-type (dashed) and representative alanine variant (solid) of each target antigen: PD-L1, IL3, IL20, and BHRF1. Solid or dashed black lines above the main elution peak indicate fractions pooled for downstream experiments.

**Figure S29.**
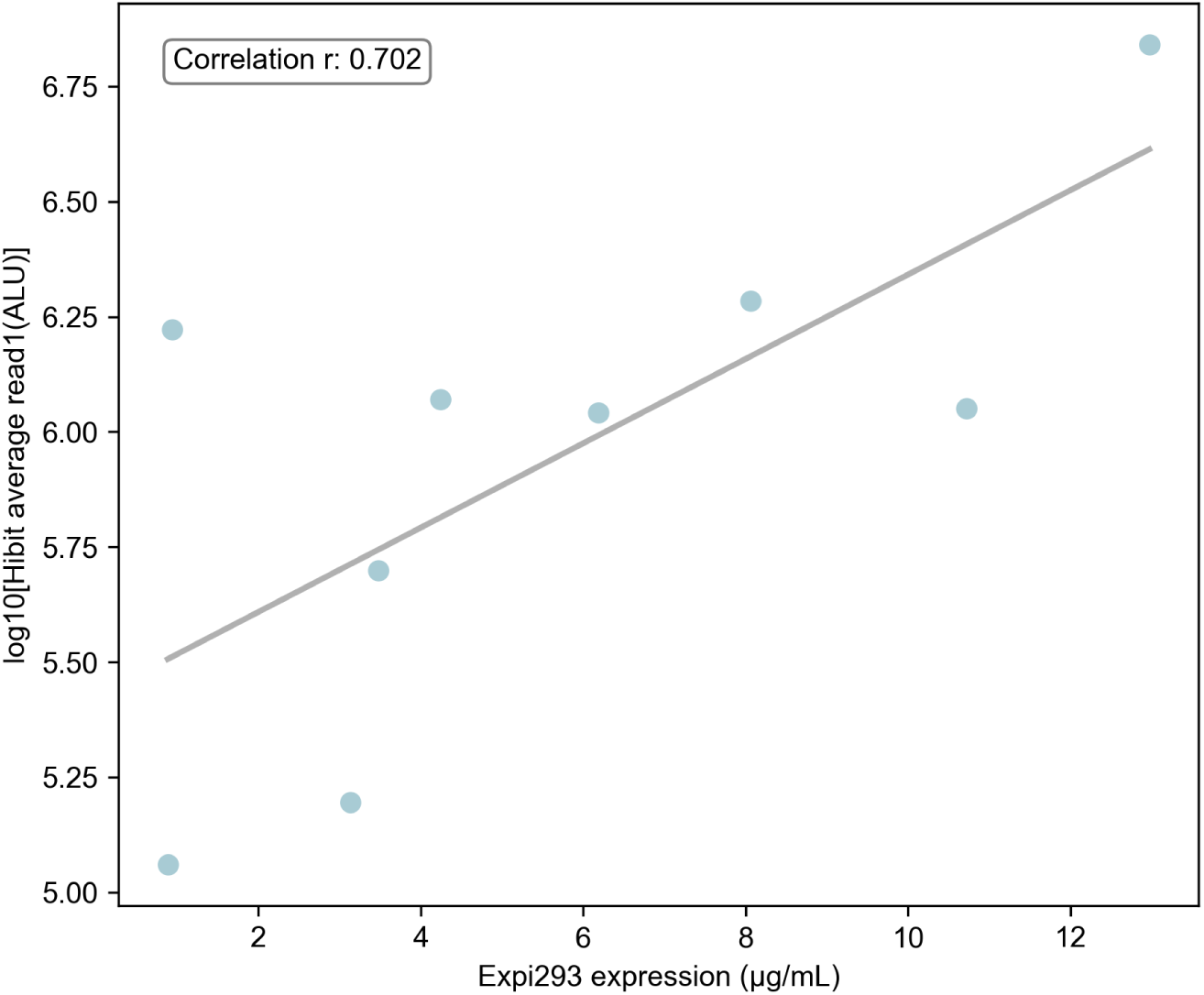
Correlation between luminescence signal (ALU) and Expi293F expression level. The Expi293F expression data were obtained from the same batch of transfections used for the PD-L1 binder designs, with validated protein concentrations and recorded final volumes, to ensure consistent normalization across samples.

**Figure S30.**
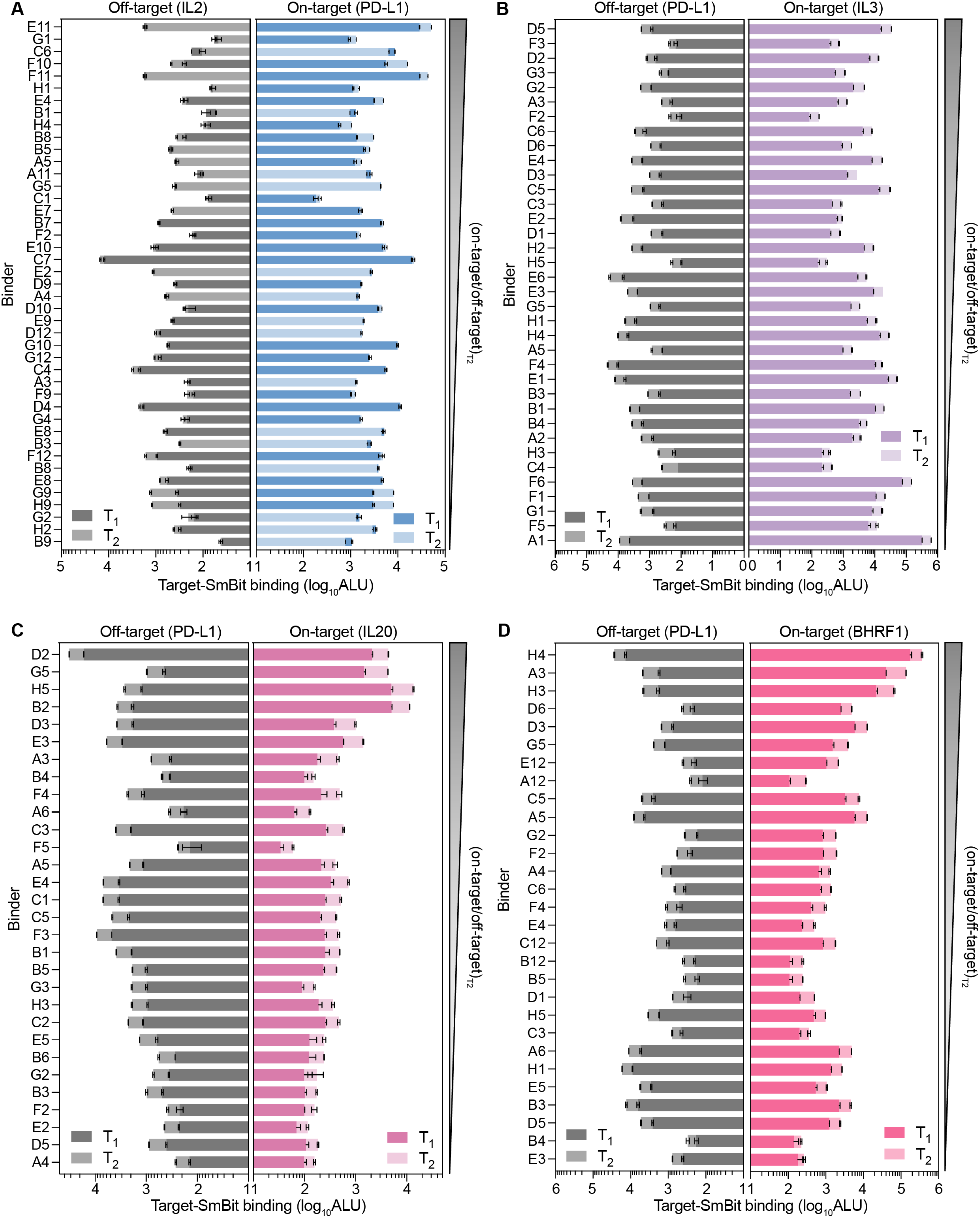
Raw on-target and off-target binding values at two time points. Raw luminescence values from the split luciferase (SmBiT–LgBiT) binding assay for (**A**) PD-L1, (**B**) IL3, (**C**) IL20, (**D**) BHRF1 binders are shown. Each design was tested against its intended on-target antigen (right panels) and an off-target antigen (left panels) at two time points, T1 (dark shade) and T2 (light shade) (8 minutes after T1). Binders are plotted if they pass the expression threshold of >150,000 RLU as determined by the HiBiT assay for expression.

**Figure S31.**
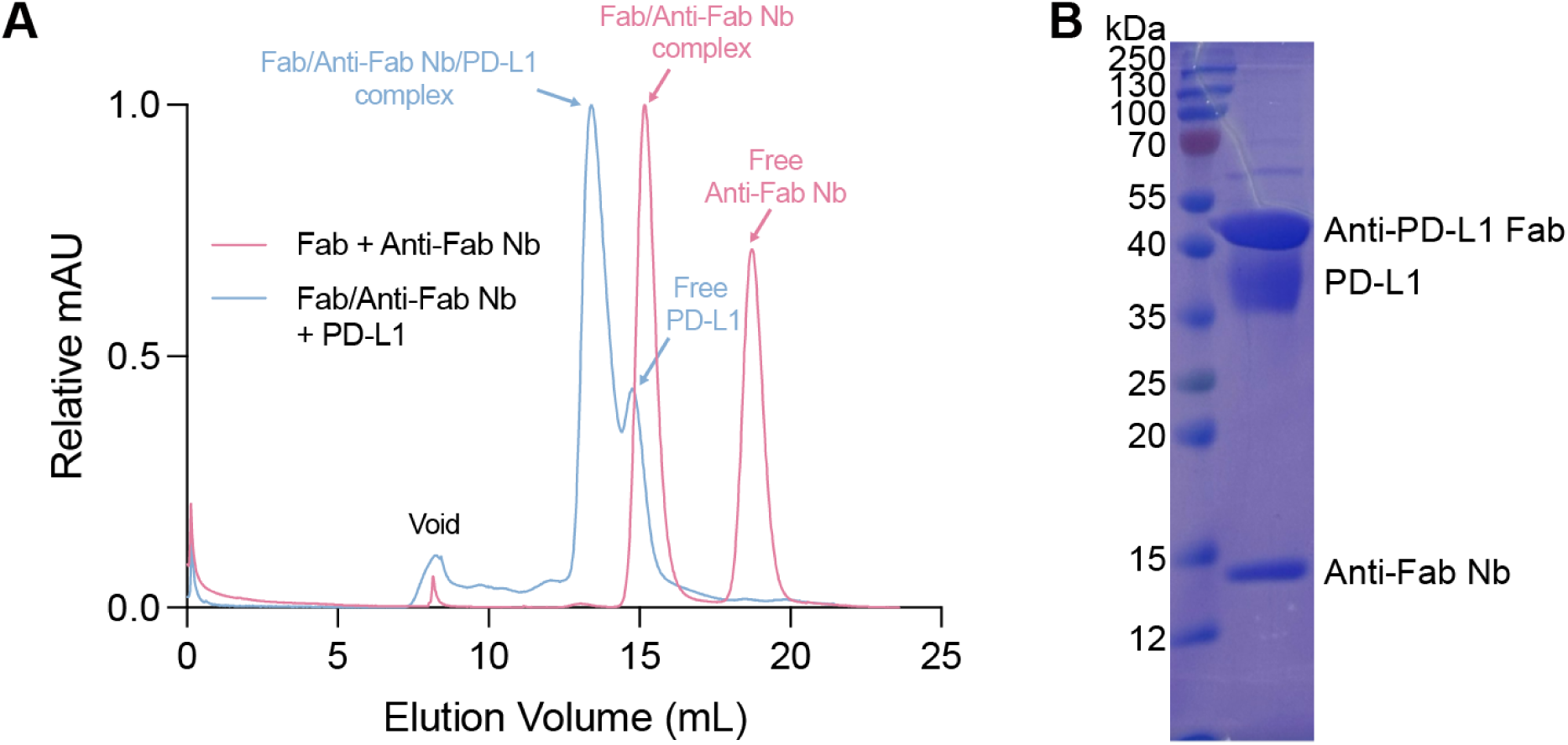
Assembly and purification of the Fab–anti-Fab nanobody–PD-L1 complex for cryo-EM analysis. (**A**) Representative SEC profiles following assembly of the Fab–anti-Fab nanobody complex (blue) and the ternary Fab–anti-Fab nanobody–PD-L1 complex (red). (**B**) Non-reducing SDS-PAGE analysis of the purified Fab–anti-Fab nanobody–PD-L1 complex used for cryo-EM data collection.

**Figure S32.**
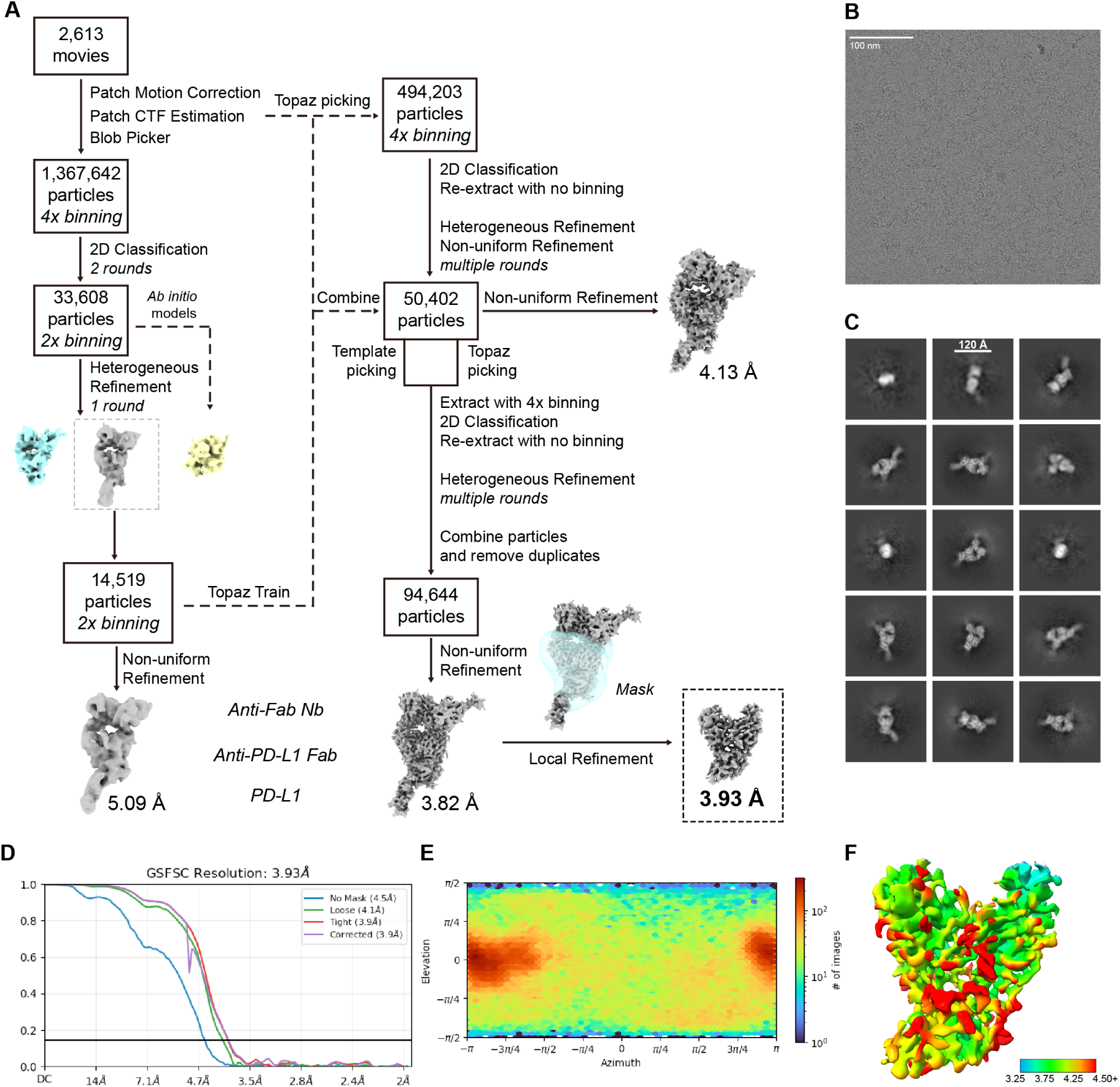
Cryo-EM data processing of the Fab–anti-Fab nanobody–PD-L1 complex. (**A**) Single-particle processing pipeline used for structure determination of the Fab–anti-Fab nanobody–PD-L1 complex. (**B**) Representative cryo-EM micrograph. (**C**) Selected 2D class averages. (**D**) Gold-standard Fourier shell correlation (FSC) curve for the final map after masked local refinement at the Fv–antigen interface; the 0.143 criterion is indicated. (**E**) Angular distribution plot of particle orientations contributing to the final reconstruction. (**F**) Local resolution estimation of the final map, colored by resolution.

## Notes

https://github.com/SantiagoMille/germinal

### Summary of Updates

This revision adds new results including scFv data for IL3 and PDL1, polyreactivity analysis, alanine mutagenesis across all targets, and cryo EM characterization.

