## Supplementary material for "Efficient generation of epitope-targeted *de novo* antibodies with Germinal": supp_file1_plasmidmaps

| Name | Description | Benchling link |
|---|---|---|
| cmv-nanobody-histag | CMV-driven nanobody fused to a HisTag. An IL2 signal peptide is used. | https://benchling.com/s/seq-ZpaqEyScgBkr6YuF4yRU?m=slm-Af2UBJwSDOY6wv02W9Ex |
| cmv-nanobody-lgbit | CMV-driven nanobody fused to LgBiT. An IL2 signal peptide is used. | https://benchling.com/s/seq-f8C1ePIiWOAnwtz623Dk?m=slm-8nTXsQDGY4FICwBHYIGn |
| cmv-antigen-avi-histag | CMV-driven target antigen with C-terminal AviTag and HisTag. | https://benchling.com/s/seq-MViseYn2zOzeNZ9tV6QI?m=slm-1UavHDzfByRO9Qj4zx1r |
| cmv-avi-antigen-histag | CMV-driven target antigen with N-terminal AviTag and C-terminal HisTag. Avi tag is placed after the antigen's signal peptide. | https://benchling.com/s/seq-bxcoEKhTsEQ2cpHJCkCr?m=slm-DaySeYCY1N0eO9gL08OD |
| cmv-antigen-smbit-histag | CMV-driven nanobody fused to SmBit and HisTag (C-term). An IL2 signal peptide is used. | https://benchling.com/s/seq-QWtaxtFdINLliasUkhx9?m=slm-kYdi1kEFxjZmIYNIATxH |
